# Loss of plasma membrane conductance in outside-xylem zone explains non-stomatal control of transpiration

**DOI:** 10.1101/2025.02.15.638441

**Authors:** Piyush Jain, Sabyasachi Sen, Fulton E. Rockwell, Robert J. Twohey, Annika E. Huber, Sahil A. Desai, I-Feng Wu, Tom De Swaef, Mehmet M. Ilman, Anthony J. Studer, N. Michele Holbrook, Abraham D. Stroock

## Abstract

The conventional assumption is that stomatal conductance (*g*_*s*_) dominates the regulation of water and carbon dioxide fluxes between leaves and the atmosphere. Here, a nanoreporter of water status at the mesophyll cell surface and local xylem within intact maize leaves documents significant undersaturation of water vapor in the outside-xylem zone (OXZ) and a large loss of conductance of this zone (*g*_*oxz*_) at moderate xylem water stress (no turgor loss). The ratio of the resistances (1/*g*_*oxz*_)/(1/*g*_*s*_) serves as a predictive phenotype of undersaturation, non-stomatal regulation of transpiration, errors in standard gas exchange analysis, and an increase of intrinsic water use efficiency (*iWUE*). Cell-scale access to water status reveals symplasmic-apoplasmic disequilibrium and informs a biophysical model that can explain experimental observations quantitatively based on localization of variable conductance to the plasma membrane. This work opens new paths of inquiry into the molecular basis and functional consequences of non-stomatal regulation of transpiration.

## Introduction

Leaves play a dominant role in the exchange of water (transpiration, *E*), carbon dioxide (net assimilation, *A*), and energy with the atmosphere (**Fig. 1a**).^1^ Their ability to regulate this exchange in response to changing soil water availability, vapor pressure deficit (*VPD*), and other environmental parameters defines plant productivity, efficiency, and resilience.^2^ Incomplete understanding of these modes of regulation have hindered our ability to predict, manage, and modulate plant function in both ecological^3^ and agricultural contexts.^4^

**Figure 1:**
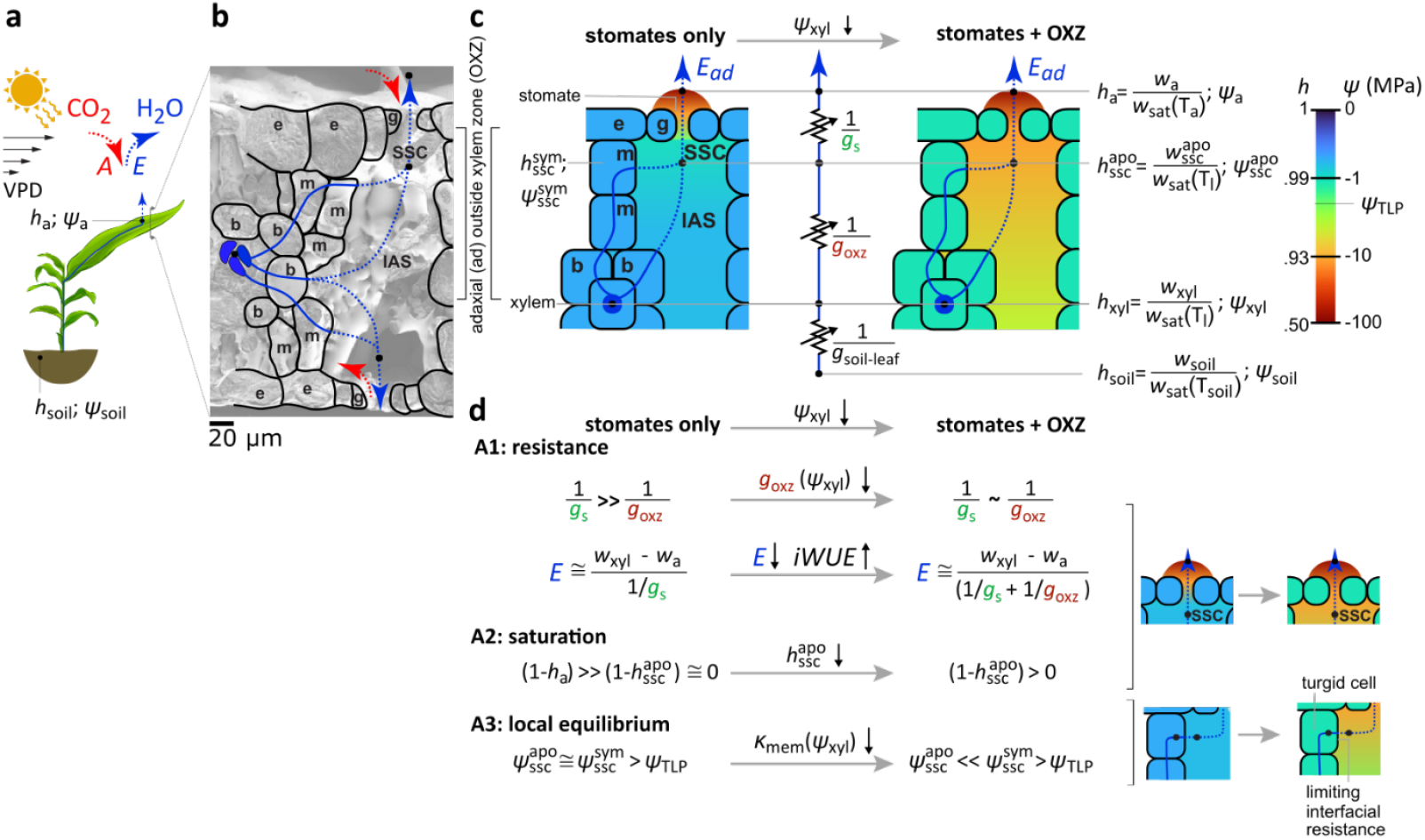
Stomatal and non-stomatal control of transpiration. **a**. Schematic diagram of soil-plant-atmosphere continuum and gas exchange. The transpiration (*E* [mmol/m^2^/s] – blue curve) from high water status in the soil (relative humidity, *h*_*soil*_ ≅ 1 [-]; water potential, *ψ*_*soil*_ ≅ *0* [MPa]) to the low water status in the atmosphere (*h*_*a*_ *<* 1; *ψ*_*a*_ ≪ *0*) is actively regulated in response to environmental signals like soil water availability (*ψ*_*soil*_), thermal loading and light availability, and vapor pressure deficit between the leaf and atmosphere (*VPD*). This regulation also affects assimilation rates (*A* [mmol/m^2^/s] – red arrow). **b-c**. Outside xylem zone (OXZ) and stomates. Cryo-electron micrograph (b) and schematic diagrams (c) of cross-section of a maize leaf identifying xylem (blue shaded), bundle sheath (s), mesophyll (m), epidermal (e) and guard (g) cells, intercellular air spaces (IAS), substomatal cavity (SSC), and stomates. Water flows as a liquid (solid blue curves) through cell-to-cell paths and diffuses as a vapor (dashed blue curves) through the IAS, SSC, and stomates. On its path from the soil to the atmosphere we define three resistances (zigzag lines in (c)) in series: from soil to xylem in the leaf (1/*g*_*soil−leaf*_), from the local xylem to the SSC apoplasm at water status 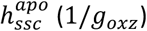, and from the SSC through stomates to the atmosphere (1/*g*_*s*_), where *g*_*soil−leaf*_, *g*_*oxz*_, *g*_*s*_ are conductances per unit area of leaf [mmol/m^2^/s]. Two scenarios are depicted in (c): the resistance of stomates alone control *E* (stomates only – left) and the added resistances of the OXZ provides an additional, non-stomatal control of *E* (stomates + OXZ – right). **d**. Conventional assumptions about regulation of transpiration rate and their violation by large resistance in OXZ. A1: Stomates have been assumed to present the limiting resistance to transpiration (1/*g*_*s*_ *≫* 1/*g*_*oxz*_); if the OXZ presents a resistance of the same order of magnitude (1/*g*_*s*_∼1/*g*_*oxz*_) it will participate in limiting transpiration (1/*g*_*oxz*_ in denominator of expression for *E* on right). A2: If stomates are limiting, the internal water status should remain near saturation (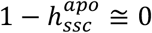-blue in diagram on right); if the OXZ also limits flow, the inside of the leaf will become unsaturated (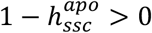-orange in diagram on right). A3: If the SSC remains saturated, then the local apoplasm and symplasm can remain near equilibrium without loss of cell turgor (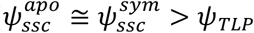, where *ψ*_*TLP*_ is the water potential at the tugor loss point); if the SSC become significantly undersaturated, then large disequilibrium must exist between the apoplasm and symplasm 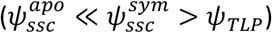 and the interface between these domain must present a large resistance to water flow. The values of water status in the diagrams in c and d are color coded based on the color map in (c).

In transpiration, water moves from the soil to the atmosphere through the plant along a gradient of water potential (*ψ* [MPa]) or equivalent relative humidity (*h = w*/*w*_*sat*_ where *ww* and *w*_*sat*_ are the mole fraction of water vapor and its value at saturation) from the nearly saturated soil (*ψ*_*soil*_ ≅ *0; h*_*soil*_ ≅ 1) to the undersaturated air in the atmosphere (*ψ*_*a*_ ≪ *0; h*_*a*_ *<* 1) (Fig. 1a). Within a leaf (Fig. 1b-c), water passes from the xylem through the outside xylem zone (OXZ) formed of bundle sheath cells (b), mesophyll cells (m), and intercellular air spaces (IAS), to the sub-stomatal cavities (SSC) upstream of the stomatal pores defined by guard cells (g) in the epidermis (e).

Three coupled assumptions define conventional understanding of the physiology that regulates gas exchange: **A1** -stomata present the limiting resistance (1/*g*_*s*_ [mmol/m^2^/s]) to, and so dominant point of control for, transpiration (*E* [mmol/m^2^/s]) and net assimilation (*A* [*μ*mol/m^2^/s]).^5^ This assumption implies that the hydraulic resistances associated with the entire upstream path from soil to the SSC are negligible relative to that of the stomates. With the inherent lack of selectivity of stomates to the outward diffusion of water vapor relative to the inward diffusion of carbon dioxide (CO_2_), this assumption also implies a significant constraint on the modulation of intrinsic water use efficiency (*iWUE* = *A*/*E*) via evolution,^4^ selection,^6^ or engineering^7^ of stomatal properties. **A2** -If A1 holds, the inside of the leaf can remain close to saturation and one can assume that the deviation from saturation remains small relative to the deficit outside the stomates 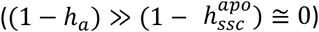. Importantly, in conventional analyses of gas exchange, this assumption is used to close the equations for water and CO_2_ fluxes to estimate stomatal conductance as, 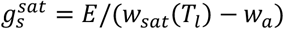, where *w*_*sat*_ is the mole fraction of water vapor at saturation at the temperature of the leaf, *T*_*l*_ (Supplementary Sec. S3; <u>LI-COR</u>).^5^ **A3** -If A2 holds, local equilibrium of water status can be maintained between the symplasm and apoplasm 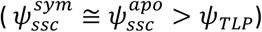 without loss of turgor during active transpiration (i.e., when stomata are open). This assumption of local equilibrium underpins current models of water transport through the OXZ^8,9^ and stomatal regulation.^10,11^

Two early studies with gas exchange using inert gases^12^ and leaves stripped of their epidermis^13^ suggested the emergence of a non-negligible resistance within the OXZ, in violation of assumption A1. Other interrogations of the water relations of leaves using a combination of gas exchange measurements, cell probes, and leaf psychrometers^14,15^ or of the evaporative method and the pressure chamber^16–18^ have been compatible with all three assumptions, without excluding possible violations of one or more of them (Supplementary Sec. S4). These studies with the pressure chamber,^16–18^ and studies by our team with the nanoreporter AquaDust,^19,20^ reported loss of conductance of the OXZ with increasing leaf or xylem water stress without violation of A1. We performed our previous work in maize at low values of vapor pressure deficit relative to the leaf, *VPD* ≅ *0*.*7* − *2* kPa.^19^ These studies also concluded that the resistance upstream of the leaves (1/*g*_*soil*−*leaf*_) remains negligible for moderate drought. Recent measurements employing inline O^18^ discrimination with gas exchange,^21–23^ dual-sided gas exchange,^24,25^ or both^26,27^ have again challenged these assumptions with model-dependent inferences of significant undersaturation within transpiring leaves of a variety of species (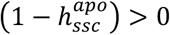 – violating A2), and the suggestion of disequilibrium between symplasm and apoplasm in the OXZ with the emergence of a limiting hydraulic resistance at the symplasm-apoplasm interface (violating A3). These studies report decreasing 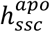 with increasing *VPD* and suggested a mechanistic connection to transport through the plasma membrane.^24,25,27^ The macroscopic and model-dependent nature of these recent studies leave open alternative scenarios that could explain the directly measured processes without involving significant loss of tissue conductance (A1), undersaturation (A2), or symplasmic-apoplasmic disequilibrium (A3).^28,29^

To test the validity of these assumptions and the consequences of their violation we use a nanoreporter, AquaDust, to access localized water status within the OXZ (A2), thereby assessing the conductance of the OXZ relative to that of the stomates (A1), and interrogating the state of symplasmic and apoplasmic equilibrium (A3) in intact, transpiring leaves.^28–31^ We then address the dependence of any observed non-stomatal modulation of gas exchange on water availability (*ψ*_*xyl*_) and demand (*VPD*) and derive a mechanistic, predictive framework based on physiological and physical processes in the OXZ.

## Results

Here we focus on maize (*Zea mays L*. W22 inbred line) infiltrated with AquaDust (**Fig. 2a**). AquaDust exploits Föster Resonance Energy Transfer (FRET) between donor and acceptor dyes bound in a hydrogel matrix.^19,20,32^ Once infiltrated into a leaf, these nanoreporters coat the cell walls surrounding the IAS of the OXZ (red outline in **Fig. 2a**). AquaDust signal from leaf regions with obstructed transpiration (both sides taped – BT) provides an estimate of xylem water status upstream of the OXZ (*h*_*xyl*_ or *ψ*_*xyl*_), and signal from unobstructed transpiring leaf regions (no tape – NT) provides a direct measurement of the water status downstream of the OXZ at the SSC 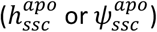.^19,20^ Importantly, the latter quantity provides a direct measure of the water vapor partial pressure in the leaf airspace interior to the stomata, allowing us to evaluate *g*_*s*_ without assumption A2 of near saturation within the leaf (Eq. 1 in Fig. 2a)^5^ and *g*_*oxz*_ (Eq. 2 in Fig. 2a).^19,20^ See Supplementary Note S1-S3 and previous publications for details.^19,20^

**Figure 2:**
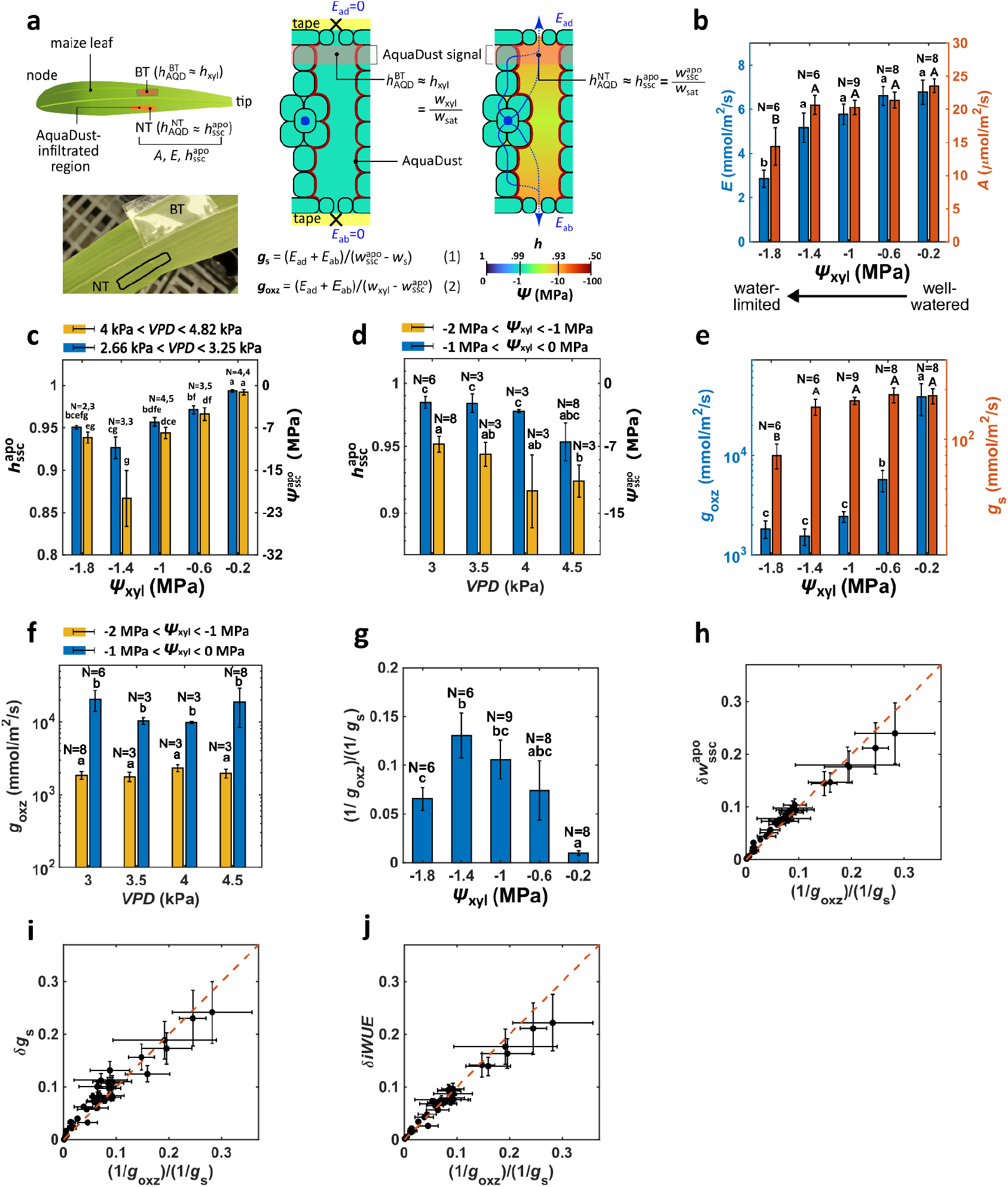
Direct measurement of undersaturation and non-stomatal control of transpiration by the OXZ. **a**. Use of AquaDust to measure the water status upstream (*h*_*xyl*_ and *ψ*_*xyl*_) and downstream 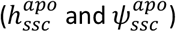 of the OXZ on intact leaves with gas exchange analysis. For a transpiring location on the leaf (labeled ‘NT’ for ‘No-Tape’), the AquaDust signal reports 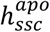; for a non-transpiring location on the leaf (labeled ‘BT’ for ‘Both-sides Taped’), the sub-stomatal cavity is in near equilibrium with the local leaf xylem such that the AquaDust signal reports *h*_*xyl*_. Gas exchange measurements on the NT region provide rates of transpiration (*E* = *E*_*ad*_ *+ E*_*ab*_ [mmol/m^2^/s]) and CO_2_ assimilation (*A* [μmol/m^2^/s]). The conductances of the stomates (Eq. 1) and OXZ (Eq. 2) can be defined with these potentials without an assumption of internal saturation. See previous publications for detailed methods.^1,2^ **b**. Transpiration rates (*E* – blue bars, left axis) and assimilation rates (*A* – red bars, right axis) as a function of *ψ*_*xyl*_ from well-watered (*ψ*_*xyl*_ *=* −*0*.*2 ± 0*.*2* MPa) to near the TLP (*ψ*_*xyl*_ *=* −1.*8 ± 0*.*2* MPa). **c-d**. Saturation of SSC with water availability (*ψ*_*xyl*_ - c) and demand (*VPD* -d). AquaDust measurements of 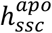 (left axis) and 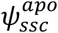 (right axis) as a function of *ψ*_*xyl*_ for low *VPD* (blue bars) and high *VPD* (yellow bars) (c) and as a function of *VPD* for high *ψ*_*xyl*_ (low stress – blue bars) and low *ψ*_*xyl*_ (high stress – yellow bars). **e**. Variations in *g*_*oxz*_ and *g*_*s*_ as a function of *ψ*_*xyl*_. **f**. Variations in *g*_*oxz*_ as a function of *VPD* for well-watered (blue bars) and water-limited (yellow bars) cases. **g**. Ratio of OXZ hydraulic resistance (1/*g*_*oxz*_) to stomatal resistance (1/*g*_*s*_) as a function of *ψ*_*xyl*_. **h-j**. Ratio of resistances, ((1/*g*_*oxz*_)/(1/*g_s*)) as a predictive phenotype of impact of non-stomatal regulation of transpiration. Relative degree of undersaturation of SSC 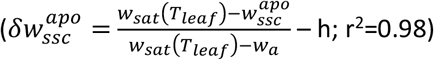, relative error in the conventional assessment of *g*_*s*_ based on assumption of saturation in IAS 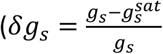 where 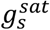 is the standard value of stomatal conductance assuming saturation of SSC – i; r^2^=0.93), and relative gain in intrinsic water-use efficiency 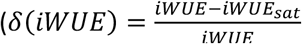 where *iWUE*_*sat*_ is the value assuming saturation of SSC – j; r^2^=0.97). See **Results** and Supplementary Sec. S3 for explanation and calculations. In frames b, c, e, and g, the measurements are binned in ranges of *ψ*_*xyl*_: *0* to −*0*.*4*, −*0*.*4* to −*0*.*8*, −*0*.*8* to −1.*2*, −1.*2* to −1.*6*, and −1.*6* to −*2* MPa. In frames d and f, measurements are binned in ranges of *VPD*: *2*.*66* to *3*.*25, 3*.*25* to *3*.*75, 3*.*75* to *4*.*25*, and *4*.*25* to *4*.*82* kPa. In frames b-g, N represents the number of biological replicates and statistically significant differences at the p<0.05 level are labeled with different letters.

### Undersaturation of leaf airspaces evolves in response to both xylem water status and VPD

For 37 plants and 87 measurements, we varied xylem water status (*ψ*_*xyl*_) and evaporative demand relative to leaf (*VPD*) independently (see Supplementary Fig. S1). With decreasing *ψ*_*xyl*_ for all values of *VPD*, the transpiration rate, *E* [mmol/m^2^/s] and net CO_2_-assimilation rate, *AA* [μmol/m^2^/s] decreased but only fell significantly as *ψ*_*xyl*_ approached the turgor loss point (*ψ*_*TLP*_ ≅ −*2* MPa) (**Fig. 2b**). For intermediate values of *ψ*_*xyl*_ (−1.4±0.2 MPa) we observe the emergence of significant internal undersaturation 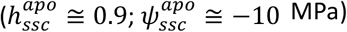 at both high and low *VPD* (**Fig. 2c;** blue bars vs. yellow bars). At the lowest value of 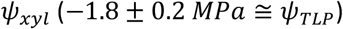, water status in the SSC recovered as the reduction of transpiration due to stomatal closure reduced the dynamic pressure drop across the OXZ. Undersaturation tended to increase modestly (lower apoplastic humidity) with increasing *VPD* across both ranges of xylem stress (*ψ*_*xyl*_,**Fig. 2d;** blue bars vs. yellow bars). Importantly, these direct measurements of undersaturation in the IAS (violation of assumption A2) confirm, qualitatively, previous model-dependent assessments of this phenomenon in maize and other species^21–27^. Quantitatively, we have not observed values of 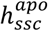 as low as those reported previously 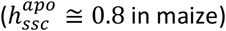.^25^ Exploration of this quantitative difference is a priority^30^ and should yield interesting new insights, for example, on the distribution of sites of evaporation and uptake of carbon dioxide within the OXZ.^33^

### Decline in outside xylem zone conductance evolves in response to xylem water status, not VPD

Evaluating the outside-xylem zone conductance, *g*_*oxz*_ (**Fig. 2e**, blue bars) based on AquaDust (Figs. 2c-2d) and gas exchange (Fig. 2b) data, we observed a dramatic drop (25-fold) that starts well above *ψ*_*TLP*_. In contrast, *g*_*s*_ (Fig. 2e, red bars) only decreased significantly (3-fold) in the most stressed plants (*ψ*_*xyl*_ *=* −1.*8 ± 0*.*2* MPa). While *g*_*oxz*_ responded strongly to *ψ*_*xyl*_ (Fig. 2e), it showed no significant dependence on *VPD* (**Fig. 2f**), revealing that the effects of *VPD* on 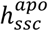 (Fig. 2d) are dynamic, resulting from an increase in the flux rather than a change in OXZ conductance. We observed these same trends in 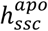 and *g*_*oxz*_ as a function of *ψ*_*xyl*_ in four other inbred genotypes in maize (Supplementary Sec. S5, Figs. S3-S5) and in a C_3_ species (*Vicia Faba L*.; Supplementary Sec. S6, Fig. S6).

### The ratio g_s_/g_oxz_ captures the degree of undersaturation and its effects on gas exchange

To clarify the importance of declines in *g*_*oxz*_ for regulation of gas exchange, in **Fig. 2g**, we plot the ratio of the resistances of the OXZ and the stomates: (1/*g*_*oxz*_)/(1/*g*_*s*_) *= g*_*s*_/*g*_*oxz*_. In well-watered maize (*ψ*_*xyl*_ *=* −*0*.*2 ± 0*.*2* MPa), the stomatal resistance was completely dominant (*g*_*s*_/*g*_*oxz*_ ≅ *0*.*0*1) as conventionally assumed (A1 – Fig. 1d). At intermediate stresses (*ψ*_*xyl*_ *=* −*0*.*6* to −1.*4* MPa) for which *g*_*s*_ remained high as *g*_*oxz*_ dropped (Fig. 2e), the ratio grew, such that the resistance of the OXZ made a non-negligible contribution to the total resistance to transpiration (*g*_*s*_/*g*_*oxz*_ ≅ *0*.1*5*). With the closure of the stomates at the most stressed conditions (*ψ*_*xyl*_ *=* −1.*8 ± 0*.*2* MPa), the importance of the OXZ resistance diminished (*g*_*s*_/*g*_*oxz*_ ≅ *0*.*06*).

To test the value of this ratio as a predictive phenotype of non-stomatal regulation of transpiration, we plot relative degree of undersaturation in internal water status 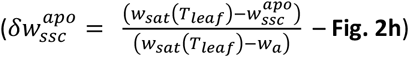, relative error in conventional gas exchange analysis based on assumption 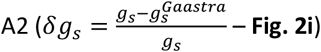, and relative gain in 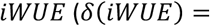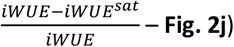 as a function of *g*_*s*_/*g*_*oxz*_ for all biological replicates in the maize inbred W22 (See Supplementary Note S3 for definitions). We note the strong, linear correlation of all of these signatures of non-stomatal regulation with this metric across all values of *ψ*_*xyl*_ and *VPD* tested. We find a further confirmation of the predictive value of *g*_*s*_/*g*_*oxz*_ in our measurements in tomato (Fig. S7) and cotton (Fig. S8): *g*_*s*_/*g*_*oxz*_ remained small (0.015 in tomato and cotton – SI Fig. S7d, SI Fig. S8d), such that even as *g*_*oxz*_ dropped several fold as the plants passed from their well-watered states to near *ψ*_*TLP*_, no substantial undersaturation occurred (SI Fig. S7a, SI Fig. S8a).

The consequences of this non-stomatal control are quantitatively important. For the largest values of *g*_*s*_/*g*_*oxz*_the errors in the conventional estimate of *g*_*s*_ can be as high as *24*% (Fig. 2i). As a test of the appropriateness of this assessment of error, in Fig. S8, we show that the values of δ*g*_*s*_ based on AquaDust (Fig. 2i) agree well with those based on a carbon balance using chlorophyll fluorescence.^34^ We also observe that the error in *iWUE* can grow to ∼ *22*% as the internal limitation on transpiration grows relative to that of the stomates (Fig. 2j). This observation suggests that the underlying processes that lead to a drop in *g*_*oxz*_ act in a selective manner on the flux of water relative to that of CO_2_.

### Undersaturation is accompanied by disequilibrium between symplasm and apoplasm

Motivated by observation of large degrees of undersaturation in apparently turgid leaves, we combined confocal microscopy (Zeiss LSM880; Olympus LMPlanFL N 20x/0.4 objective) with AquaDust to create maps of water status with cell-scale resolution within the mesophyll of intact leaves in the cuvette of a gas exchange system (CIRAS-3, PPSystems) (**Fig. 3a**). In order to maintain robust levels of transpiration under the low light levels required for imaging, measurements were performed on a maize mutant (*slac1-2* null in W22 inbred background) in which stomates remain open except at turgor loss.^35^ Confocal sections (*x*-*y* planes – **Fig. 3b**) of tissue autofluorescence and AquaDust emission were collected to assess pixel-level water status. See Supplementary Sec. S8.

**Figure 3:**
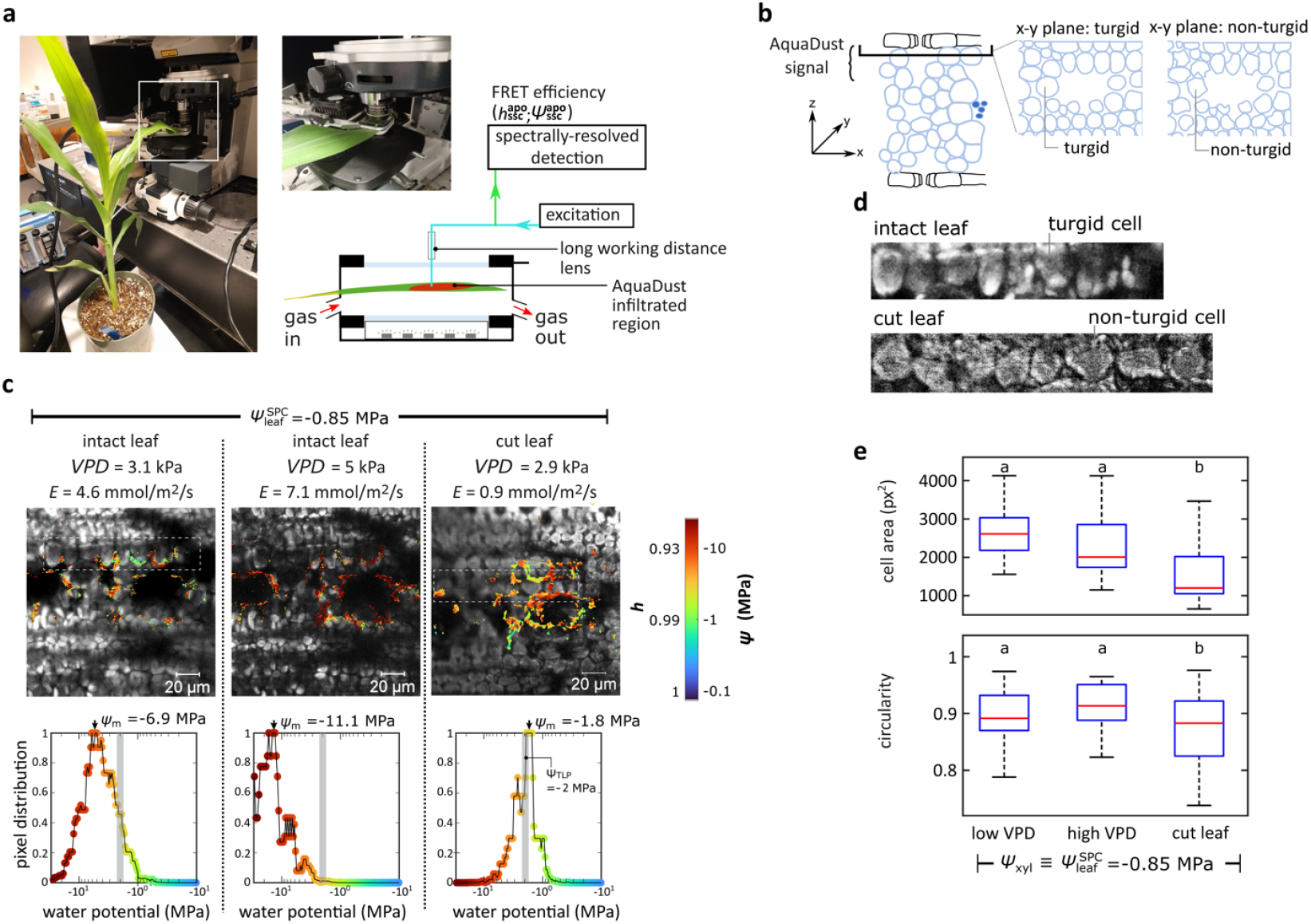
Extreme local gradients of water potential between mesophyll symplasm and SSC. **a**. Spectrally resolved confocal imaging within a gas exchange cuvette in which a section of intact leaf infiltrated with AquaDust is clamped. The native fluorescence of the tissue and a pixel-wise assessment of FRET efficiency (*ζ*(*ψ*)) are captured at a depth of *z* ≅ *24 μm*. **b**. Schematic diagrams indicating the location of the confocal section on a vertical cross-section (x-z – left) and two examples of the cellular structure in the captured horizontal cross-sections (x-y – right): turgid and turgor-loss states. **c-d**. Horizontal confocal sections (x-y) of fluorescence and AquaDust FRET and pixel-level distribution of water potential (c) for three experimental conditions applied to a section of leaf of a maize *slac1-2* null mutant in the gas exchange cuvette: water limited 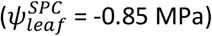 [for low *VPD* (3.1 kPa), high *VPD* (5 kPa), and after excision of leaf (*VPD* = 2.9 kPa). Here, micrographs present AquaDust-reported water potential maps (color map on right) overlaid on intensity of the 686-695 nm-channel in which chloroplasts are visible. Expanded views of a file of cells show turgid and turgor-loss states with intensity in the 686-695 nm-channel (d). In (c), histograms present of pixel-level values of water potential extracted from the images above, with the grey shaded bars representing the bulk tissue water potential measured using a Schölander Pressure Chamber 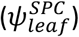. **e:** Cell area and circularity for the three cases with Tukey HSD test (n>20 for each case and p<0.05 for significant difference) using cell outlines. For intact leaf cases, mesophyll cells remain smooth and rounded (left expanded view in (d)), indicating the maintenance of turgor even as AquaDust-reported water potentials in the mesophyll cell wall descended to a mode value, 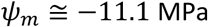, well below the overall water potential of the tissue, 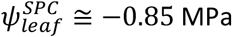. Upon excision from plant, the cells shrank and lost their smooth, rounded shape, indicating loss of turgor (right expanded view in (d)) and AquaDust-reported water potentials recovered to a mode value, *ψ*_*m*_ ≅ −1.*8* MPa, near the turgor-loss point of maize (TLP ≅ -2 MPa). See Supplementary Sec. S8 for methods and additional confocal data.

In a leaf with moderate xylem stress (transpiring leaf, 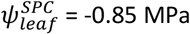), AquaDust measurements of mesophyll cell wall water status from pixels surrounding the cells adjacent to the substomatal cavities (elliptically shaped dark zones) showed a shift toward lower pixel-level water potential (redder color; left-shifted histogram) as *VPD* and *E* increased (*VPD* = 3.1 to 5 kPa and *E* = 4.6 to 7.1 mmol/m^2^/s) (**Fig. 3c**). Notably, the mode values of the distributions of pixel-level water potentials for both values of *VPD* (*ψ*_*s*_ = -6 and -11 MPa) indicate strong undersaturation, consistent with our macroscopic measurements as presented in Fig. 2c. Upon cutting the leaf, the transpiration rate decreased with stomatal closure, and the pixel-level values of water potential shifted toward higher potentials, with *ψ*_*s*_ approaching the range expected for the turgor loss potential, *ψ*_*s*_ ≅ −1.*8* MPa. (see Supplementary Sec. S8, Supplementary Figs. S13-S17).

In the expanded views of the cells’ autofluorescence in **Fig. 3d**, we note a qualitative change in the shape of the cells between the intact (high *VPD* - left) and cut (right) states, with smooth, rounded edges for the intact case and more faceted surfaces in the cut case. Quantitatively, we found a significant drop in both size and circularity of cells between the intact leaf and cut leaf cases (**Fig. 3e** - See Supplementary Sec. S8 for details)) which we interpret as evidence of the presence and loss of turgor respectively. This interpretation is consistent with macroscopic observations of turgidity and the high stomatal conductance of the intact leaves, and the onset of leaf rolling and reduced transpiration in the cut case (Supplementary Fig. S16). Taken together, the cell-scale documentation of strong undersaturation directly surrounding turgid cells in intact leaves undergoing steady transpiration (Figs. 3c-e) provides direct evidence for the violation of assumption A3 of local equilibrium: a large drop in water potential occurred between the turgid symplasm (*ψ*_*sym*_ *> ψ*_*TLP*_ ≅ -2 MPa) and the exterior of the cell wall at which AquaDust reports 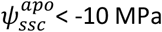 (Fig. 3c). In other words, nearly the entire potential drop from xylem to the sites of evaporation occurred in the passage of water across plasma membrane and cell wall; one or both of these components must present the dominant resistance to the transpiration stream in the OXZ. As discussed below and in Supplementary Sec. S10, these observations also place constraints on the hydraulic properties of the symplastic path through the OXZ.

### Modeling undersaturation selects from among candidate hydraulic architectures

To gain further insights into the physiological basis of these observations, we present a pseudo-one-dimensional model (**Fig. 4a**) of isothermal flow of water through the OXZ in which we can account for a cell-to-cell compartment (blue dashed line), an apoplasmic compartment (red dashed line), the coupling of these compartments through an interface defined by the plasma membrane and cell wall (green dashed line), and the downstream path through stomata. In Supplementary Sec. S10, we derive this model and explore four scenarios informed by the literature; we exclude three of these hypotheses as incompatible with our observations. We present a compatible scenario that predicts the disequilibrium observed in Fig. 3 based on a hypothesis that the plasma membrane presents the limiting, variable resistance within the OXZ, as depicted in **Fig. 4b**.^24,25^

**Figure 4:**
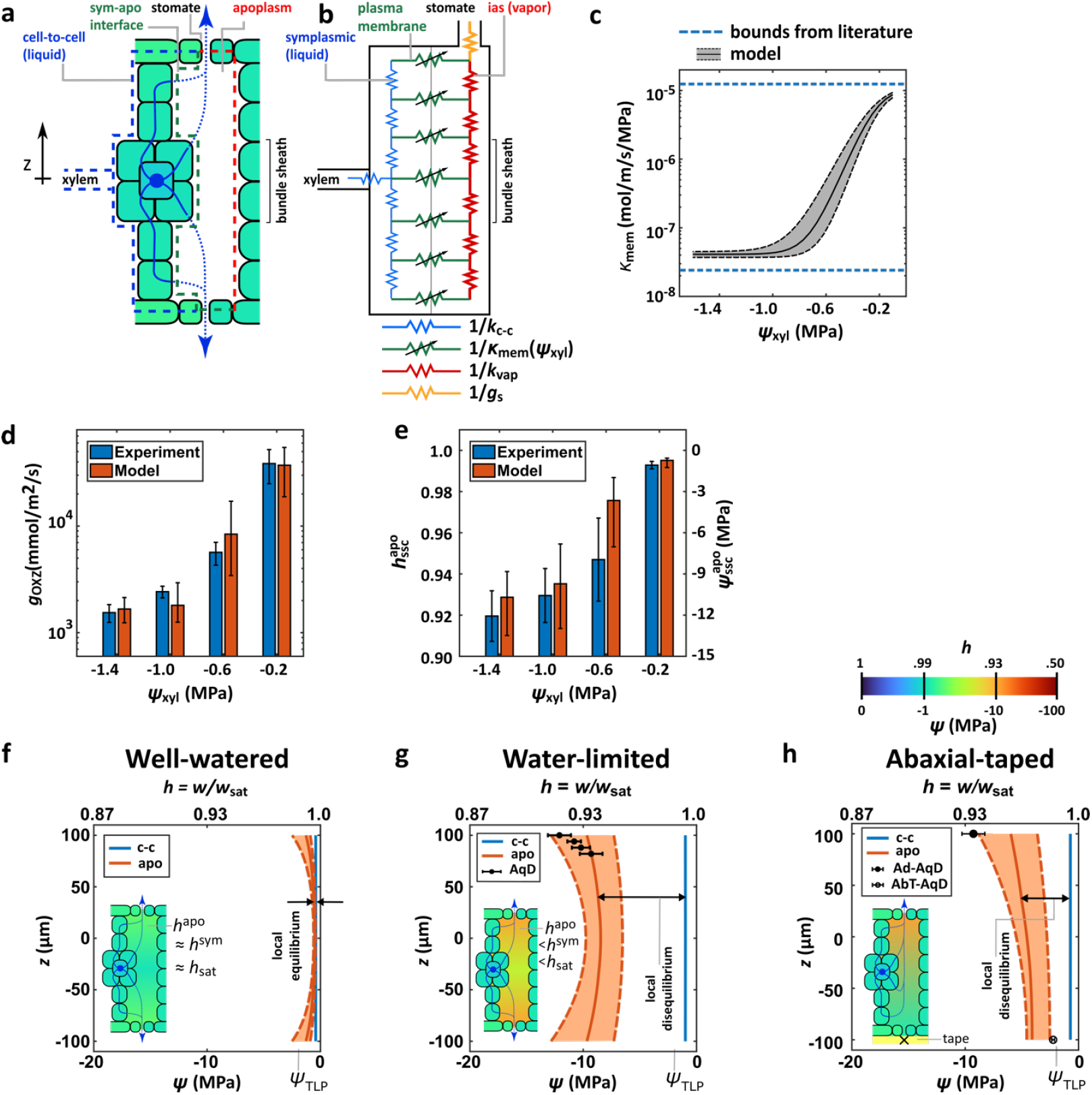
Model of leaf water transport in Outside-Xylem Zone with large loss of conductance in plasma membrane. **a-b**. Two-compartment, pseudo-one-dimensional, continuum, steady-state model of water flow through OXZ introduced here for the hypothetical scenario in which the plasma membrane at the interface between the cell-to-cell and apoplasmic paths presents a limiting, variable conductance, κ_*mem*_. The cell-to-cell path (blue dashed outline; blue resistors in (b) defined by 1/*k*_*c*−*c*_) is coupled to the apoplasmic path (red-dashed outline in (a); red resistors in (b) defined by 1/*k*_*apo*_) through the plasma membrane (green dashed line in (a); green resistors in (b) defined by 1/κ_*mem*_); the apoplasm feeds into a stomatal resistance (orange resistor in (b) defined by 1/*g*_*s*_). Here, the hydraulic conductances, *k*_*c*−*c*_ and *k*_*apo*_ [mol/m/s/MPa] and vapor conductance, *g*_*s*_ [mmol/m^2^/s] are defined per unit area of leaf; the hydraulic conductance, κ_*mem*_ [mol/m/s/MPa] is defined per unit area of the symplasm-apoplasm interface. **c**. Hypothetical functional dependence of conductance of plasma membrane, κ_*mem*_ on *ψ*_*xyl*_ used in the model presented in (b) for predictions in (d)-(h). The upper and lower bounds on κ_*mem*_ are from experiments on maize leaf protoplasts.^29,36^ **d-e**: Comparison of predictions from model based on variable conductance in (c) with experimentally observed *g*_*oxz*_ (d) and 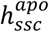 (e) and as a function of *ψ*_*xyl*_ for *VPD = 3*.*8 ± 0*.*2* kPa (mean VPD *±* S.E. for data in Fig. 2). **f-h**: Through-leaf profiles of water potential, *ψ*(*z*) (bottom axes) and *h*(*z*) (top axes – approximating linear dependence of *h* with psi) in cell-to-cell path (*ψ*_*c*− *c*_ – blue curve) and apoplasm (*ψ*_*apo*_ – orange curve) predicted for well-watered (*ψ*_*xyl*_ = -0.4 *±* 0.15 MPa – f), water-limited (*ψ*_*xyl*_ = -0.85 *±* 0.15 MPa; *VPD = 4*.*2* kPa – g), and water-limited with obstructed transpiration from the adaxial surface (*ψ*_*xyl*_ = -1.2 MPa *±* 0.15 MPa; *VPD = 4*.*2* kPa – h). Insets show schematic representations of profiles with colormap (h), with near equilibrium maintained between symplasm and apoplasm at low stress (both blue, near saturation – f) and disequilibrium between cell-to-cell path (blue) and apoplasm (green) at moderate stress (g and h). In both cases, *ψ* _*c* − *c*_ remains above the turgor loss point (TLP ≅ -2 MPa), consistent with data presented in Fig. 3. In g and h, predicted profiles in apoplasm match measurements made with AquaDust in confocal sections (filled circles in adaxial SSC in g and h; open circle in abaxial SSC in h - see Supplementary Sect. S11 for additional details).

The scenario that best fits the observed data includes the following (Fig. 4b): (i) a variable interfacial resistance (green zigzag line with cross arrow indicating variability) dominated by the conductance of the plasma membrane with dependence on the upstream water potential (κ_*mem*_ *ψ*_*xyl*_ [mol/m/s/MPa] per interfacial area); (ii) a cell-to-cell resistance dominated by the conductivity of plasma membranes or plasmodesmata or both (see Supplementary Sec. S10) (*k*_*c−c*_ [mol/m/s/MPa] per leaf area); and (iii) an axial apoplasmic resistance dominated by vapor conductivity (*k*_*vap*_ [mol/m/s/MPa] per leaf area) (See SI Sec. S19). We model κ_*mem*_ (*ψ*_*xyl*_) as a sigmoid that respects the bounds on the hydraulic conductance of plasma membrane from experiments performed on maize leaf protoplasts (dashed blue lines; **Fig. 4c**).^29,36^ In **Fig. 4d**, we present predictions of this model for *g*_*oxz*_ with a range of values informed by the literature for fixed values *k*_*c*_ −_*c*_ and *kk*_*vap*_ and *g*_*s*_ (180 (mmol/m^2^/s)) and imposed *VPD* (3.8*±*0.2 kPa – mean value of VPD in Fig. 2 data). We adjusted the parameters of κ_*mem*_ *ψ*_*xyl*_ to provide a best fit to measured *g*_*oxz*_ for these conditions (Fig. 4d; See Sec. S10 for details and Tables S1-S5 for parameter values). The corresponding predictions of internal 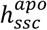 are compatible with our measurements (**Fig. 4e**), providing a first quantitative check of the appropriateness of our model.

For well-watered conditions (**Fig. 4f** – *ψ*_*xyl*_ *=* −*0*.*4 ± 0*.1*5* MPa) for which κ_*mem*_ is high, the symplasmic and vapor paths remain at or near equilibrium, with *ψ*^*apo*^ ≅ *0* (*h*^*apo*^ ≅ 1) throughout. For a water limited condition that corresponds to the experiment in Fig. 3c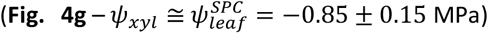, our preferred model predicts strong symplasmic-apoplasmic disequilibrium throughout the thickness and deep undersaturation near the stomates 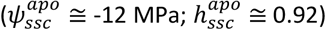. For a case with obstructed transpiration from the abaxial surface (abaxial taped, AbT – *ψ*_*xyl*_ ≅ −1.*2 ± 0*.1*5* MPa), the model predicts a monotonic gradient in the vapor from the non-transpiring abaxial surface (*E*_*ab*_ = 0) to the transpiring adaxial surface (*E*_*ad*_ *>* 0) accompanied by strong disequilibrium (**Fig. 4h**).

Importantly, the predicted disequilibrium in both cases (Figs. 4g-h) is compatible with the maintenance of high symplasmic water potential and the maintenance of turgid mesophyll cells adjacent to deeply undersaturated vapor. Furthermore, the predicted water potential profiles match confocal-based AquaDust measurements at multiple depths within the leaf (Fig. 4g) and macroscopic AquaDust measurements collected at the adaxial and abaxial surfaces (Fig. 4h), providing a second quantitative check of the appropriateness of the preferred model.

## Discussion

Our measurements and model elucidate the physiology underlying the emergence of non-stomatal control of transpiration based on loss of conductance in the OXZ. The direct measurement of undersaturation and symplasm-apoplasm disequilibrium impose important constraints on existing models of leaf hydraulics^8,9^ and stomatal responses.^10,11^ The identification of the ratio (1/*g*_*oxz*_)/(1/*g*_*s*_) as a predictive phenotype for undersaturation, errors in conventional calculations of *g*_*s*_, and favorable variations in *iWUE* creates new opportunities to pursue implications for ecophysiological function and the development of water-use efficient crops. With access to water status at the cellular scale within leaves and a new model of water movement through the OXZ, we have localized the variations in *g*_*oxz*_ to the plasma membrane. This localization resolves important questions about the physiological viability of internal undersaturation^30^ and opens a path to identify the molecular basis of this stress response, with aquaporins as an obvious starting point (see Supplementary Sec. S12 and Fig. S24). Together, these methods, measurements, and models provide a foundation for the necessary reformulation of our existing understanding of leaf gas exchange (assumptions A1-A3) to account for active, non-stomatal management of water-use.

## Methods

### AquaDust synthesis and use

AquaDust nanoparticles (70-100 nm diameter) were formed of hydrogel containing two matrix-bound fluorescent dyes enabling FRET-based sensing of water potential, as described previously.^32^ Supplementary Sec S1 presents details of the synthesis, characterization, injection protocol, calibration, and simultaneous measurement water potential and gas exchange.

### Plant material and growing conditions

Maize (*Zea mays* L. W22 inbred line) plants were grown in a greenhouse under controlled conditions (28 ± 2°C day/20 ± 2°C night temperature, 40% RH day/80-90% RH night) with natural sunlight supplemented to maintain minimum PAR of 1000 μmol m^−2^ s^−1^ during a 16h photoperiod. Plants were grown in potting medium with slow-release fertilizer and measurements were conducted on uppermost newest, fully expanded, leaves at the V9-V10 vegetative growth stage. Additional genotypes and species used for comparative measurements were grown under identical conditions (see Supplementary Sec. S2 for details).

### Confocal Imaging

Intact leaves from slac1-2 null mutants (W22 background) were infiltrated with AquaDust and imaged using a Zeiss LSM880 confocal microscope with a long working distance 20x objective (LMPlanFL N 20x/0.4, Olympus) while maintaining the leaves in a gas exchange cuvette (CIRAS-3, PPSystems). Two-channel imaging was performed using 488 nm and 561 nm excitation to collect AquaDust FRET signals and tissue autofluorescence. Water potential was calculated pixel-by-pixel from FRET efficiency using calibrated response curves. Cell morphology analysis was performed using ImageJ on autofluorescence images. Detailed imaging protocols, calibration procedures, and analysis methods are provided in Supplementary Sec S8.

## Acknowledgements

We thank Jacob L. Wszolek, Nicholas Kaczmar, Amy L. Soriano, Scott A. Anthony, and Jeffrey T. Persky for maintaining the plants in the greenhouse and growth chamber, Glenn Swan for technical support and Graham Farquhar, Diego Marquez, Lukas Cernusak, Florian Busch, Ying Sun, Vesna Bacheva, and Warren Zipfel for valuable discussions. This research was performed with support from the Center for Research on Programmable Plant Systems under National Science Foundation Grant No. DBI-2019674, the Air Force Office of Scientific Research under Grant No. FA9550-18-1-0345, the Harvard MRSEC, DMR-2011754, and a Star-Friedman Challenge award (Harvard University) and an OECD-CRP fellowship award (TAD/CRP PO 0500109311). Imaging data were acquired through the Cornell Institute of Biotechnology’s Imaging Facility, with NIH S10OD018516 funding for the shared Zeiss LSM880 confocal/multiphoton microscope. Cryo-imaging was performed in part at the Harvard University Center for Nanoscale Systems (CNS); a member of the NNCI supported by the National Science Foundation under NSF award no. ECCS-2025158. This work was also performed in part at the Cornell Nanoscale Facility, an NNCI member supported by NSF Grant NNCI-2025233.

## Contributions

P.J., S.S., F.E.R., and A.D.S conceived the project. P.J., S.S., F.E.R., A.E.K., S.A.D., I-F.W., T.D.S., R.J.T., and M.M.I. performed the experiments. S.S., F.E.R., and A.D.S. developed the model. R.J.T. and A.J.S. developed the germplasm. P.J., S.S., F.E.R., and A.D.S. analyzed the data. P.J., S.S., F.E.R. and A.D.S. wrote the manuscript. All authors reviewed and revised the manuscript. P.J. and S.S. contributed equally to this paper.

## Conflict of interest

P.J. and A.D.S. are listed as inventors on US Patent 11,536,660 titled, ‘In situ sensing of water potential’ owned by Cornell University.

## Data Availability Statement

All the data used in this study has been included in the main text or in the supplementary information files.

## Code Availability Statement

All the code used in this study has been included in the supplementary information files.

## Notes

### Competing Interest Statement

Piyush Jain and Abraham D. Stroock are listed as inventors on US Patent 11,536,660 titled, 'In situ sensing of water potential' owned by Cornell University.

