## Supplementary Information for "Loss of plasma membrane conductance in outside-xylem zone explains non-stomatal control of transpiration"

Water transport across plasma membrane: 32
Coupled governing equations: 32
Boundary and interfacial conditions: 33
Transpiration rate and conductance of OXZ: 34
Solution technique 34
Predictions for four hypothetical scenarios 35
Preferred hypothesis: Large symplasmic conductance and both mesophyll and bundle sheath plasma membranes undergo loss of conductivity to water under drought stress (Fig. 4 in main text) 35
Alternative hypothesis 1: Local equilibrium between cell-to-cell path and apoplasm and cell-to-cell path becomes less permeable to water under drought stress 37
Alternative hypothesis 2: Cell wall is the dominant hydraulic pathway, and it becomes less permeable to water under drought stress 38
Alternative hypothesis 3: Bundle-sheath membrane is the dominant hydraulic resistance in the outside-xylem tissue 39
Neglect of temperature-gradient-driven vapor transport: 40
Table S2: Transport variables 40
Table S3: Thermodynamic parameters 41
Table S4: Leaf anatomical parameters 41
Table S5: Transport parameters 42
Figure S19. Pseudo 1-D model of OXZ. 46
Figure S20. Alternative hydraulic architecture 1: ‘local equilibrium’. 48
Figure S21. Alternative hydraulic architecture 2: ‘vulnerable cell wall’. 50
Figure S22. Alternative hydraulic architecture 3: ‘vulnerable bundle sheath’. 52
S11. Depth profile of AquaDust measurements from confocal microscopy 53
Figure S23. Cellular scale measurement of $\psi_{\mathrm{scc}}^{apo}$ at varying depths. 53
S12. Response of hydraulic conductance of outside-xylem zone $g_{\mathrm{OXZ}}$ to Mercury (II) Chloride. 54
Figure S24. Response of hydraulic conductance of outside-xylem zone $g_{\mathrm{OXZ}}$ to Mercury (II) Chloride. 55
S13. Standard Operating Procedure (SOP) for handling Mercury (II) Chloride ($HgCl_2$) solution for gas-exchange and AquaDust experiments 55
Introduction. 55
Figure S25. Photo of experiment in progress with an excised maize leaf with the cut end dipped inside artificial xylem sap solution with 0.5 mM $HgCl_2$. 55
Supplies. 55
Personal Protective Equipment (PPE). 55
Procedure. 56

### S1. AquaDust synthesis and use

**Synthesis and mechanism of response:**
AquaDust is a nano-scale probe of water potential comprising nanometer-scale hydrogel spheres with Forster Resonance Energy Transfer (FRET)-paired fluorophores incorporated into the polymer network. In the FRET process, excitation of one fluorophore, the donor dye, leads to energy transfer to the second fluorophore, the acceptor dye, with an efficiency that is a function of the distance between the donor and acceptor dyes. Here, we have covalently linked FRET pairs into a polyacrylamide network, resulting in 70-100 nm diameter AquaDust nanoparticles. Changes in the water potential in the cell wall (the part of the tissue to which AquaDust is localized) result in the shrinking or swelling of the gel, which changes the efficiency of energy transfer (FRET) between the two dyes. These changes in efficiency are detected as changes in the relative intensity of the emission peaks of the donor and acceptor dyes in the fluorescence spectrum of the leaf. Please see Jain et al., 2021[1] for more details on synthesis and mechanism of response.

**Injection of AquaDust solution in leaves:**
AquaDust (20 $\mu$L per injection) was injected into maize leaves through a syringe that was gently pressed against the leaves two days before measurement to allow the aqueous buffer, in which AquaDust was dissolved, sufficient time to transpire. Successful injections were characterized by a dark green colorization of the leaf tissue (approximately 2 $cm^2$) caused by AquaDust suspension filling the leaf air spaces. To avoid AquaDust extrusion out of open stomates during injection, plants were removed from the greenhouse and placed in shade to induce stomatal closure. If stomates were shut too tightly for AquaDust injection, then leaf surfaces were gently scratched with a razor blade around the injection site. After AquaDust injection, leaves were rinsed off with water to clean the leaf from AquaDust on the leaf surface. Then, plants were returned to the greenhouse and left for ∼24 hours before measurements were taken.

**Measurement of AquaDust signal:**
To exclude artefacts in water potential data from potential tissue damage during injection, local leaf water potential measurements with AquaDust were performed at least 1 cm away from the injection site. Local water potentials in the mesophyll were measured by recording the fluorescence intensity of AquaDust. The fluorescence intensity was collected by clamping onto the leaf with a fiber optic point probe. Excitation light was emitted by a mercury lamp light and passed through a narrow-band optical filter. The reflected light was passed through a long-pass cutoff filter and transmitted to a spectrometer. The spectra were saved using OceanView software. We recorded between four and ten spectra from one infiltration zone by moving the optical probe across the infiltration zone. The mean FRET was calculated by averaging over multiple spectra. No distinction was made in the data collected from the infiltrated zone upstream/downstream of the spot on the leaf where the syringe made contact to infiltrate AquaDust. The fluorescence spectra from a leaf not

infiltrated with AquaDust were saved as background signals and subtracted from the spectra collected from AquaDust-infiltrated leaves. Further details on the synthesis, characterization, injection, and calibration of AquaDust have been described in detail previously.[1–3]

**S2. Plant growth and experimental conditions**

Species: Maize (*Zea mays L.*; Inbred lines W22, Mo17, B73, Tx303, CML322, *slac1;2* null mutant in W22 background). Growing conditions: Maize plants were grown from seed in a 1:1 ratio of Cornell Mix[4] and Turface (Turface Athletics MVP Profile, Products, LLC; Buffalo Grove, IL, USA) in 15 cm diameter standard pots. Plant material was grown under controlled conditions in a greenhouse with a 12-hour photoperiod, 30 C day and 24 C night temperature, 50% relative humidity, and a photosynthetic photon flux density of 1000 µmol/m$^2$/s photosynthetic photon flux density (PPFD). Plants were kept well-watered during the growth period. Maize plants were fertilized twice a week with Jack's 21-5-20 Fertilizer (JR Peters Inc, Allentown, PA) at 200 ppm. Experiments were conducted when maize plants were between 8-10 weeks old (vegetative stage V9-V10). One day before measurements, plants were moved into a growth chamber that simulated the planned experimental conditions of PPFD, temperature, and relative humidity, and the zone infiltrated with AquaDust that was meant for the measurement of local leaf xylem water potential ($\psi_{\mathrm{xyl}}$) was taped (see S3 for details).

**S3. Experimental procedure: Measuring outside xylem zone conductance and corrected stomatal conductance with AquaDust in a transpiring maize leaf**

The methods described here correspond to the data shown in main text Figure 2. In order to calculate the hydraulic conductance of the outside-xylem zone ($g_{\mathrm{oxz}}$) and the stomatal conductance to water ($g_{\mathrm{s}}$), we measured gas exchange variables ($CO_2$ assimilation rates ($A$), transpiration rates ($E$), etc.) followed by measurements of local leaf water potential ($\psi$) with AquaDust on the adaxial surface of the leaf. The plants were infiltrated with AquaDust on Day 0, and the experiment was conducted during the subsequent three-day period. We infiltrated two regions of the maize leaf with AquaDust and subjected them to the following treatments using an optically transparent tape that is impermeable to gas exchange (low fluorescence optical transparent tape - Nunc Sealing, Thermo Scientific Inc., Waltham, MA, USA): (i) BT - Taping both sides of the leaf such that $E_{ab} + E_{ad} \approx 0$, and (ii) NT- No taping. On the first day, well-watered plants were probed. On two consecutive days, plants were sampled with a range of varying decreasing stem water potentials (increasing drought stress) above the turgor loss point in maize. Drought stress was imposed by withholding water on days two and three, allowing us to vary $\psi_{\mathrm{xyl}}$ indirectly. On each experiment day, we measured gas exchange variables from both regions (NT and BT) of different maize plants by clamping a leaf into a gas exchange cuvette (CIRAS-3, PPSystems, Amesbury, MA, US). Measurements were taken after gas exchange parameters stabilized (typically in 15-20 minutes). The gas exchange cuvette was programmed with the following settings: $CO_2$ concentration was set at 460 ppm, cuvette temperature = 36 C, Photosynthetically active radiation PAR = 600-750 $\mu$mol/m$^2$/s (Blue light = 40 $\mu$mol/m$^2$/s; Remaining PAR = Red light), and vapor pressure deficit ($VPD_{\mathrm{leaf}}$) across the following range was used: [2.66 kPa to 4.82 kPa]; flow within the cuvette was set to be 300 cc/min.

For the data reported in Fig. 2, 87 measurements on 37 plants were performed for various values of $\psi_{\mathrm{xyl}}$ and $VPD$in an uncorrelated manner, as illustrated in Fig. S1.

After gas exchange measurements were taken, we measured the local adaxial leaf water potential ($\psi_{\mathrm{ssc}}^{\mathrm{apo}}$; $h_{\mathrm{ssc}}^{\mathrm{apo}}$) on the same leaf on which gas exchange was measured, with an optical point probe, and finally, the water potential at the BT location ($\psi_{\mathrm{xyl}}$; $h_{\mathrm{xyl}}$). The effectiveness of the taped region in providing a faithful representation of $\psi_{\mathrm{xyl}}$ has been described earlier.[2,3]

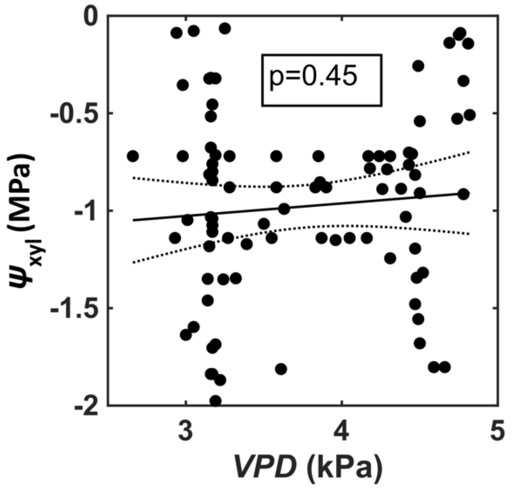

**Fig. S1. Lack of correlation between *VPD* and $\psi_{\mathrm{xyl}}$.**
In our controlled-environment experiments, we independently regulate $\psi_{\mathrm{xyl}}$ and *VPD* by changing the water potential in the soil and by changing the vapor pressure in the cuvette of a gas exchange system, respectively. This way we could isolate the effects of $\psi_{\mathrm{xyl}}$ and *VPD* and attribute the physiological origin of the phenomenon of non-stomatal control of transpiration to a reduction in $\psi_{\mathrm{xyl}}$ and not to rising VPD.

**Corrected stomatal conductance and intrinsic water use efficiency ($iWUE$).**
With the data obtained in this manner, we calculate the effective outside xylem zone conductance as:

$$g_{\mathrm{oxz}} = \frac{E^{\mathrm{ad}}+E^{\mathrm{ab}}}{w_{\mathrm{xyl}}-w_{\mathrm{ssc}}^{\mathrm{ad}}} \qquad \text{(S1)}$$

The stomatal conductance is estimated from the equations in the LICOR LI6800 manual with the following modification: in the expression that represents the total conductance of a leaf to water vapor flux

$$g_{\mathrm{s}} = \frac{E\left(1-\frac{w_{\mathrm{i}}+w_{\mathrm{a}}}{2}\right)}{w_{\mathrm{i}}-w_{\mathrm{a}}} \qquad \text{(S2)}$$

the value of $w_i$ (= $w_{\mathrm{ssc}}^{\mathrm{apo}}$) measured with AquaDust is used instead of the saturation vapor pressure at the leaf temperature, $w_{\mathrm{sat}}(T_{\mathrm{leaf}})$. The stomatal conductance, $g_{\mathrm{s}}$, and the outside xylem zone conductance, $g_{\mathrm{oxz}}$, are both expressed in [mmol/m$^2$/s]. Fig. S2 presents the variation of both correct stomatal conductance, $g_{\mathrm{s}}$ and uncorrected values, $g_{\mathrm{s}}^{\mathrm{sat}}$.

With a corrected estimate of stomatal conductance informed by $\psi_{\mathrm{ssc}}^{\mathrm{apo}}(h_{\mathrm{ssc}}^{\mathrm{apo}})$ in hand, we can define an error in the standard assessment of stomatal conductance with assumption of near saturation inside leaf: $E = g_{\mathrm{s}}^{\mathrm{sat}}(w_{\mathrm{leaf}}^{\mathrm{sat}} - w_{\mathrm{a}})$, where $g_{\mathrm{s}}^{\mathrm{sat}}$ is the standard definition of stomatal conductance with the standard assumption (A2 in main text) due to Gaastra of internal saturation.[5]

From (S1) and (S2), and by ignoring the boundary layer resistance, we have:

$$E = g_{\mathrm{tot}}\left(w_{\mathrm{xyl}} - w_{\mathrm{a}}\right) = \frac{g_{\mathrm{s}}}{1+\frac{g_{\mathrm{s}}}{g_{\mathrm{oxz}}}}\left(w_{\mathrm{xyl}} - w_{\mathrm{a}}\right) \cong \frac{g_{\mathrm{s}}}{1+\frac{g_{\mathrm{s}}}{g_{\mathrm{oxz}}}}(w_{\mathrm{leaf}}^{\mathrm{sat}} - w_{\mathrm{a}})$$

$$g_{\mathrm{s}}^{\mathrm{sat}} \cong \frac{g_{\mathrm{s}}}{1+\frac{g_{\mathrm{s}}}{g_{\mathrm{oxz}}}} \cong g_{\mathrm{s}}(1 - \frac{g_{\mathrm{s}}}{g_{\mathrm{oxz}}})$$

We thus find the following approximate relationship between the relative error in stomatal conductance and the ratio $\frac{g_{\mathrm{s}}}{g_{\mathrm{oxz}}}$ as plotted in Fig. 2i:

$$\delta g_{\mathrm{s}} = \frac{g_{\mathrm{s}} - g_{\mathrm{s}}^{\mathrm{sat}}}{g_{\mathrm{s}}} \cong \frac{g_{\mathrm{s}}}{g_{\mathrm{oxz}}} \quad \text{(S3)}$$

Undersaturated measurements of $\psi_{\mathrm{ssc}}^{\mathrm{apo}}$ also tell us that improvements to intrinsic water use efficiency ($iWUE$) under drought stress have a non-stomatal contribution that can be captured by comparing the true, measured intrinsic water use efficiency ($iWUE$) to a hypothetical $WUE$ that would have occurred with the same true stomatal conductance if the intercellular air spaces were saturated with vapor ($iWUE^{\mathrm{sat}}$). If inside of leaf remained saturated, we have:

$$iWUE^{\mathrm{sat}} = \frac{A}{E} = \frac{A}{g_{\mathrm{s}}(w_{\mathrm{leaf}}^{\mathrm{sat}} - w_{\mathrm{a}})}$$

$$iWUE = \frac{A}{g_{\mathrm{tot}}(w_{\mathrm{xyl}} - w_{\mathrm{a}})} \cong \frac{A}{g_{\mathrm{tot}}(w_{\mathrm{leaf}}^{\mathrm{sat}} - w_{\mathrm{a}})}$$

$$\frac{iWUE^{\mathrm{sat}}}{iWUE} = \frac{g_{\mathrm{tot}}}{g_{\mathrm{s}}} = \frac{1}{1+\frac{g_{\mathrm{s}}}{g_{\mathrm{oxz}}}} \cong 1 - \frac{g_{\mathrm{s}}}{g_{\mathrm{oxz}}}$$

We further predict the following approximate relationship to $\frac{g_{\mathrm{s}}}{g_{\mathrm{oxz}}}$ as plotted in Fig. 2j:

$$\delta(iWUE) = \frac{iWUE - iWUE_{\mathrm{sat}}}{iWUE} = 1 - \frac{g_{\mathrm{tot}}}{g_{\mathrm{s}}} = \frac{\frac{g_{\mathrm{s}}}{g_{\mathrm{oxz}}}}{1+\frac{g_{\mathrm{s}}}{g_{\mathrm{oxz}}}} \cong \frac{g_{\mathrm{s}}}{g_{\mathrm{oxz}}} \quad \text{(S4)}$$

**Relative undersaturation.**
From Eqs. S1 and S2, we have:

$$g_{\mathrm{s}}\left(w_{\mathrm{ssc}}^{\mathrm{apo}} - w_{\mathrm{a}}\right) = g_{\mathrm{oxz}}\left(w_{\mathrm{xyl}} - w_{\mathrm{ssc}}^{\mathrm{apo}}\right) \cong g_{\mathrm{oxz}}\left(w_{\mathrm{sat}}(T_{\mathrm{leaf}}) - w_{\mathrm{ssc}}^{\mathrm{apo}}\right)$$

Solving for $w_{\text{ssc}}^{\text{apo}}$, we have:

$$w_{\text{ssc}}^{\text{apo}} \cong \frac{g_{\text{s}} w_{\text{a}} + g_{\text{OXZ}} w_{\text{leaf}}^{\text{sat}}}{g_{\text{s}} + g_{\text{OXZ}}}$$

We can define a relative degree of undersaturation as follows. Measured values of this parameter are plotted in Fig. 2h:

$$\delta w_{\text{ssc}}^{\text{apo}} \equiv \frac{w_{\text{sat}}(T_{\text{leaf}}) - w_{\text{ssc}}^{\text{apo}}}{w_{\text{sat}}(T_{\text{leaf}}) - w_{\text{a}}} = \frac{1}{1 + \frac{g_{\text{OXZ}}}{g_{\text{s}}}} \cong \left(\frac{g_{\text{s}}}{g_{\text{OXZ}}}\right) \quad \text{(S5)}$$

**Kelvin equation.**

We use the Kelvin equation to relate water potential and relative humidity and partial molar water vapor fraction:[6]

$$\psi = \frac{RT}{\bar{v}} \ln\left(\frac{w}{w_{sat}}\right) \quad \text{(S6)}$$

The version of the Kelvin equation that corresponds to xylem water status is: $\psi_{\text{xyl}} = \frac{RT}{\bar{v}} \ln\left(\frac{w_{xyl}^{*}}{w_{sat}}\right)$.

The version of the Kelvin equation that corresponds to the water status in the intercellular airspaces next to sites of evaporation in mesophyll cell walls is: $\psi_{\text{ssc}}^{\text{apo}} = \frac{RT}{\bar{v}} \ln\left(\frac{w_{\text{ssc}}^{\text{apo}}}{w_{\text{sat}}}\right)$.

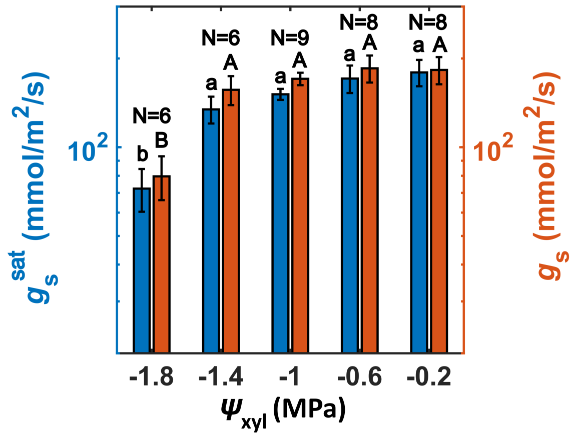

**Fig. S2. Estimates of stomatal conductance with and without AquaDust.**

Corrected estimate of stomatal conductance ($g_{\text{s}}$) using AquaDust measurement of intercellular air space water potential, $\psi_{\text{ssc}}^{\text{apo}}$, and estimate of stomatal conductance using the conventional assumption of saturation in the leaf intercellular air space ($g_{\text{s}}^{\text{sat}}$). The conventional method always underestimates the value of stomatal conductance.

**S4: Previous methods to measure OXZ conductance and associated assumptions:**

**Cell probe.**

This method evaluated the total symplastic water potential within epidermal cells ($\psi_{\text{sym-epi}}$) with the cell probe and near the xylem with a psychometer ($\psi_{\text{xyl}}$) in intact, transpiring leaves.[7,8] They then evaluated the "within leaf hydraulic conductance", $K =$

$E/(\psi_{\mathrm{xyl}} - \psi_{\mathrm{sym-epi}})$. None of their reported values of $K$ implied that this conductance could contribute to the limitation of transpiration. This technique did not provide access to the water status in the intercellular air spaces adjacent to the stomates ($\psi_{\mathrm{ssc}}^{\mathrm{apo}}$ or $h_{\mathrm{ssc}}^{\mathrm{apo}}$), and thus did not directly test for significant undersaturation (i.e., violation of assumption **A2**). With their use of symplastic water potential in the epidermis as the downstream potential driving flow through the leaf, they depended on the assumption of cell-apoplasm equilibrium (assumption **A3**). To the extent that $\psi_{\mathrm{ssc}}^{\mathrm{apo}}$ may have been more negative that $\psi_{\mathrm{sym-epi}}$, their assessment of $K$ would have been an overestimate (underestimate of the resistance) relative to the assessment provided in this study ($g_{\mathrm{oxz}}$).

**Pressure chamber.**
A common practice for the assessment of leaf and OXZ conductance is to use the pressure chamber to assess the terminal water potential in the transpiring leaf.[9] As we have argued and demonstrated previously,[2,3] the pressure chamber potential is higher (less negative) than that in the intercellular airspaces adjacent to the stomates (based on AquaDust) and provides higher values of the conductance of the OXZ than those provided in this study. As such, methods based on the pressure chamber measurements may miss the emergence of significant undersaturation (assumption **A2**) and of non-negligible resistance in the OXZ (assumption **A1**).

**S5. Loss of conductance of OXZ in additional inbred genetic backgrounds of maize.**
We use AquaDust and gas exchange data to evaluate the internal undersaturation, $h_{\mathrm{i}}$, OXZ conductance, $g_{\mathrm{oxz}}$ (blue bars), and stomatal conductance, $g_{\mathrm{s}}$ (red bars) in four additional maize inbred lines (B73: 13 plants and 37 measurements; Tx303: 8 plants and 32 measurements; CML322: 10 plants and 34 measurements; and Mo17: 9 plants and 32 measurements). The environmental conditions are described in the Methods Section. We observe a significant change in $g_{\mathrm{oxz}}$ across all genotypes (as large as ~5-fold in Mo17). In contrast, $g_{\mathrm{s}}$ only decreased significantly in two of the five genotypes, that too approximately 1.2-fold. The data shown in Fig. S3 and Fig. S4 is divided into two ranges of $\psi_{\mathrm{xyl}}$: $-0.8$ MPa $< \psi_{\mathrm{xyl}} < 0$ MPa (Well-watered) and, $-2$ MPa $< \psi_{\mathrm{xyl}} \leq -0.8$ MPa (Water-limited) given the lower number of replicates used relative to the W22 inbred genotype shown in the main text. For the sake of a complete comparison, we have included the data from main text Fig. 2 (W22 inbred) as a fifth panel in these figures. In summary, the trends are qualitatively similar across genotypes. Data represent the mean value ($\pm$S.E.) from N biological replicates (plants). Different letters (a vs. b; A vs. B) indicate statistically significant differences at the p<0.05 level.

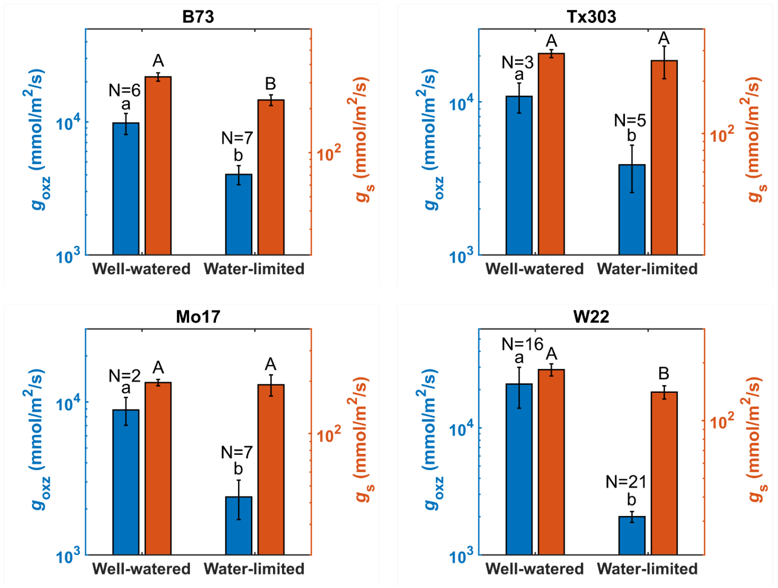

**Fig. S3. Loss of hydraulic conductance of outside-xylem zone as a function of $\psi_{xyl}$ in multiple inbred lines in maize.**

The data is divided into two ranges of $\psi_{xyl}$: $-0.8$ MPa $< \psi_{xyl} < 0$ MPa (well-watered) and, $-2$ MPa $< \psi_{xyl} \leq -0.8$ MPa (water-limited). Significant differences are observed between the treatments. Data represent the mean value ($\pm$S.E.) from N biological replicates. Different letters (a vs. b; A vs. B) indicate statistically significant differences at the p<0.05 level.

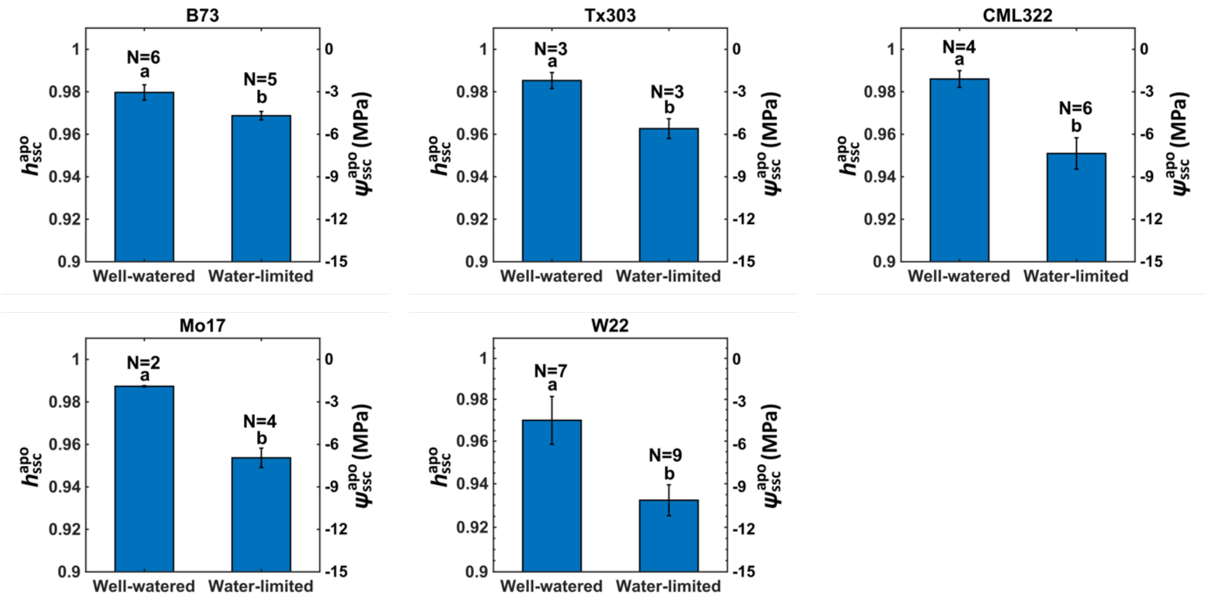

**Fig. S4. Internal relative humidity, $h_i$ as a function of $\psi_{xyl}$ in multiple inbred lines in maize for $VPD$ in the range 2 kPa to 3.5 kPa.**

The data is divided into two ranges of $\psi_{xyl}$: $-0.8$ MPa $< \psi_{xyl} < 0$ MPa (well-watered) and, $-2$ MPa $< \psi_{xyl} \leq -0.8$ MPa (water-limited). Significant differences are observed in $RH_i$ between the treatments. Data represent the mean value ($\pm$S.E.) from N biological replicates. Different letters (a vs. b) indicate statistically significant differences at the p<0.05 level.

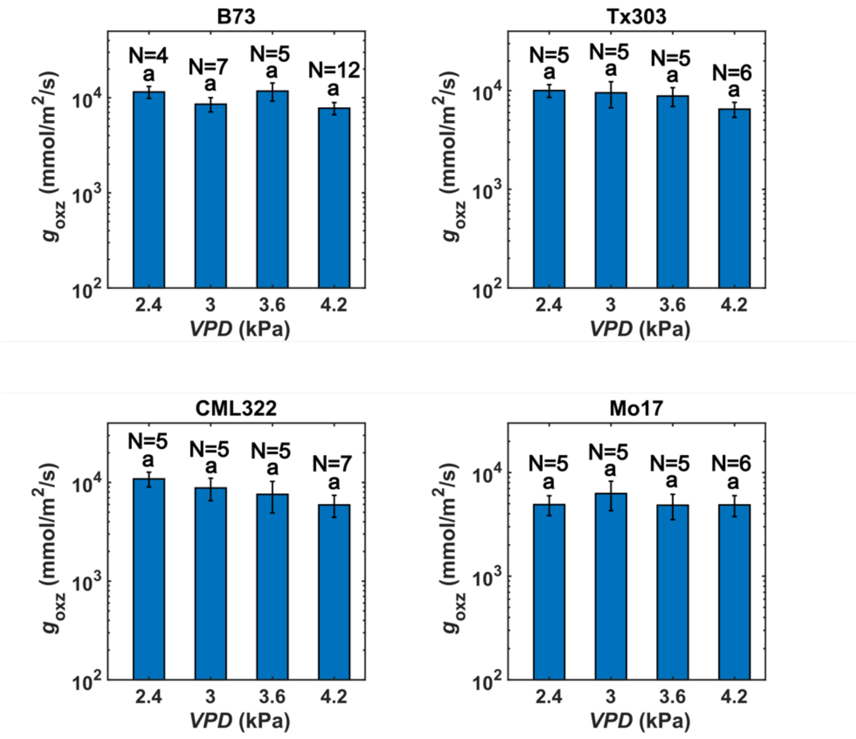

**Fig. S5. OXZ conductance, $g_{\text{oxz}}$ as a function of $VPD$ across genetic backgrounds in maize.** For the same plants shown in Fig. S3 and Fig. S4, we use AquaDust and gas exchange data to evaluate the internal conductance, $g_{\text{oxz}}$ in four additional maize inbred lines (B73, Tx303, CML322, and Mo17) as a function of $VPD$ for $\psi_{\text{xyl}}$ in the range $-1$ MPa to $-2$ MPa. We observe no dependence of $g_{\text{oxz}}$ on $VPD$, similar to what was shown in the main text for W22. Data represent the mean value ($\pm$S.E.) from N biological replicates. Different letters (a vs. b) indicate statistically significant differences at the p<0.05 level.

**S6. Loss of conductance of OXZ in C3 species: broad bean (*Vicia faba*), tomato (*Solanum Lycopersicum*), and upland cotton (*Gossypium hirsutum*).**

Data collected in broad bean (11 measurements across 5 plants) grown in Growth Chamber with conditions reported in Sec. S2 and $VPD_{\text{leaf}} \cong 2$ kPa are reported in Fig. S6. We observe significant undersaturation in the intercellular air spaces as xylem water potential decreases, similar to observations in maize. Transpiration declines with more negative $\psi_{\text{xyl}}$, whereas the assimilation rate shows a non-significant decrease, consistent with observations of non-stomatal control of transpiration in Vicia faba. The ratio $(1/g_{\text{oxz}})/(1/g_{\text{s}})$ grows with drought, indicating the growing importance of the outside-xylem zone in regulating transpiration under drought stress conditions. Data collected in tomato (15 measurements across 15 plants) as reported previously[2] and reanalyzed for comparison here (Fig. S7). In tomato, internal relative humidity, $h_{\text{ssc}}^{\text{apo}}$ stays very near saturation ($h_{\text{ssc}}^{\text{apo}} \geq 0.99$) (a) even as $g_{\text{oxz}}$ drops $\sim$ 3-fold (b) as it passes from its well-watered state to near TLP ($\psi_{\text{TLP}} \cong -0.8$ MPa). Fig. S7-c presents rates of transpiration and assimilation. Despite the drop in $g_{\text{oxz}}$ with xylem stress, $g_{\text{s}}/g_{\text{oxz}}$ remains small (0.015) (d); no substantial undersaturation occurred (a) because the upstream resistance of the OXZ never competed with the that of the stomates. Data collected in cotton (18 measurements across 8 plants) is also shown here (Fig. S8). Similar to tomato, internal relative humidity,

$h_{ssc}^{apo}$, in cotton stays very near saturation ($h_{ssc}^{apo} \geq 0.99$) (a) even as $g_{oxz}$ drops ~ 9-fold (b) as it passes from its well-watered state to water-limited state. Fig. S8-c presents rates of transpiration. Despite the drop in $g_{oxz}$ with xylem stress, $g_s/g_{oxz}$ remains small (0.015) (d); no substantial undersaturation occurred (a) because the upstream resistance of the OXZ never competed with the that of the stomates.

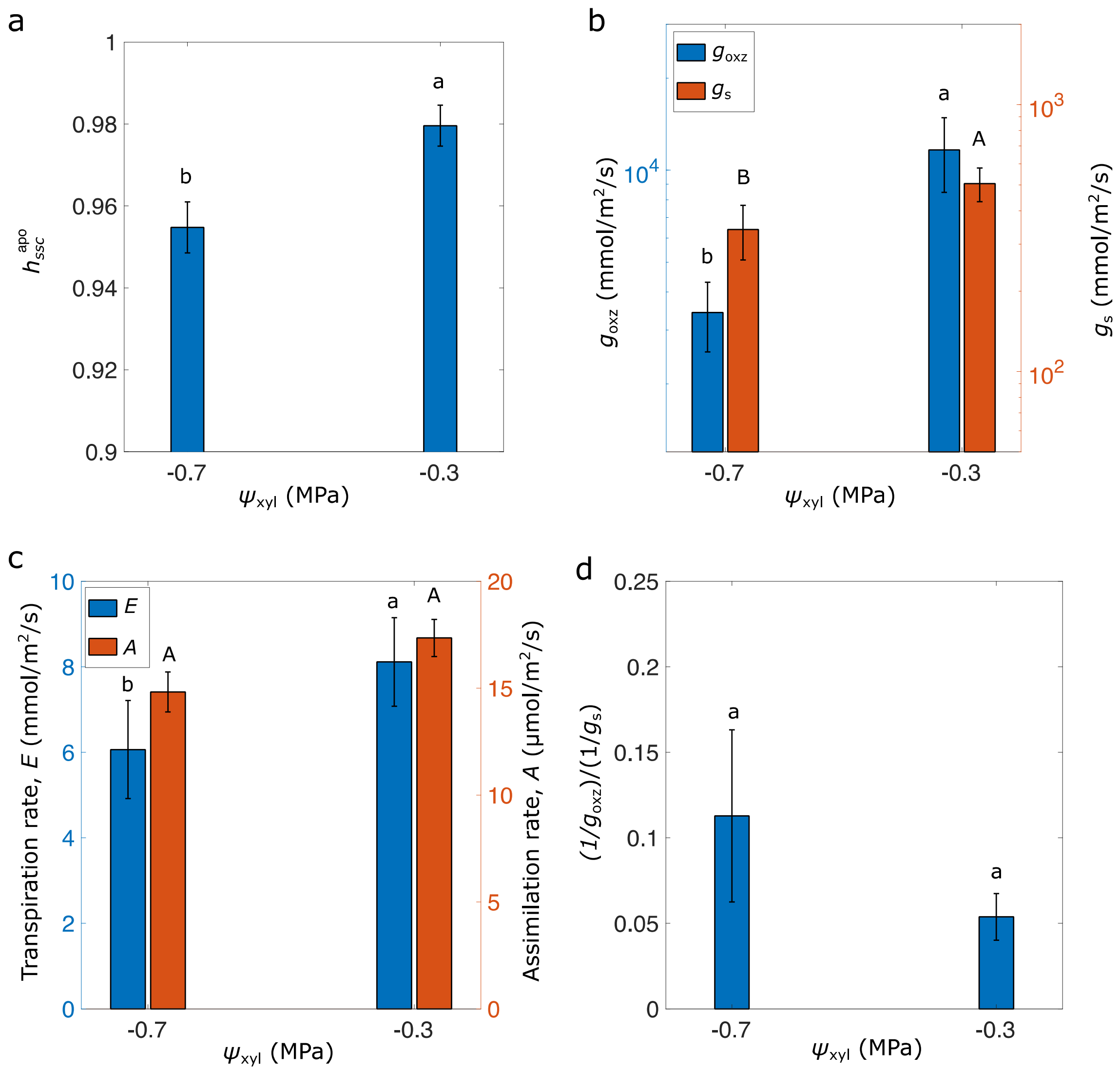

**Fig. S6. Non-stomatal control of transpiration in a C3 species *Vicia faba*.**
**a**. Relative humidity in the intercellular air spaces, reported as $h_{ssc}^{apo}$, as a function of xylem water potential ($\psi_{xyl}$). **b**. Outside-xylem zone conductance ($g_{oxz}$) and stomatal conductance ($g_s$) as a function of xylem water potential ($\psi_{xyl}$). **c**. Transpiration rate (E) and $CO_2$ assimilation rate (A) as a function of $\psi_{xyl}$. **d**. Ratio of outside-xylem zone hydraulic resistance ($1/g_{oxz}$) to stomatal resistance ($1/g_s$) as a function of $\psi_{xyl}$.

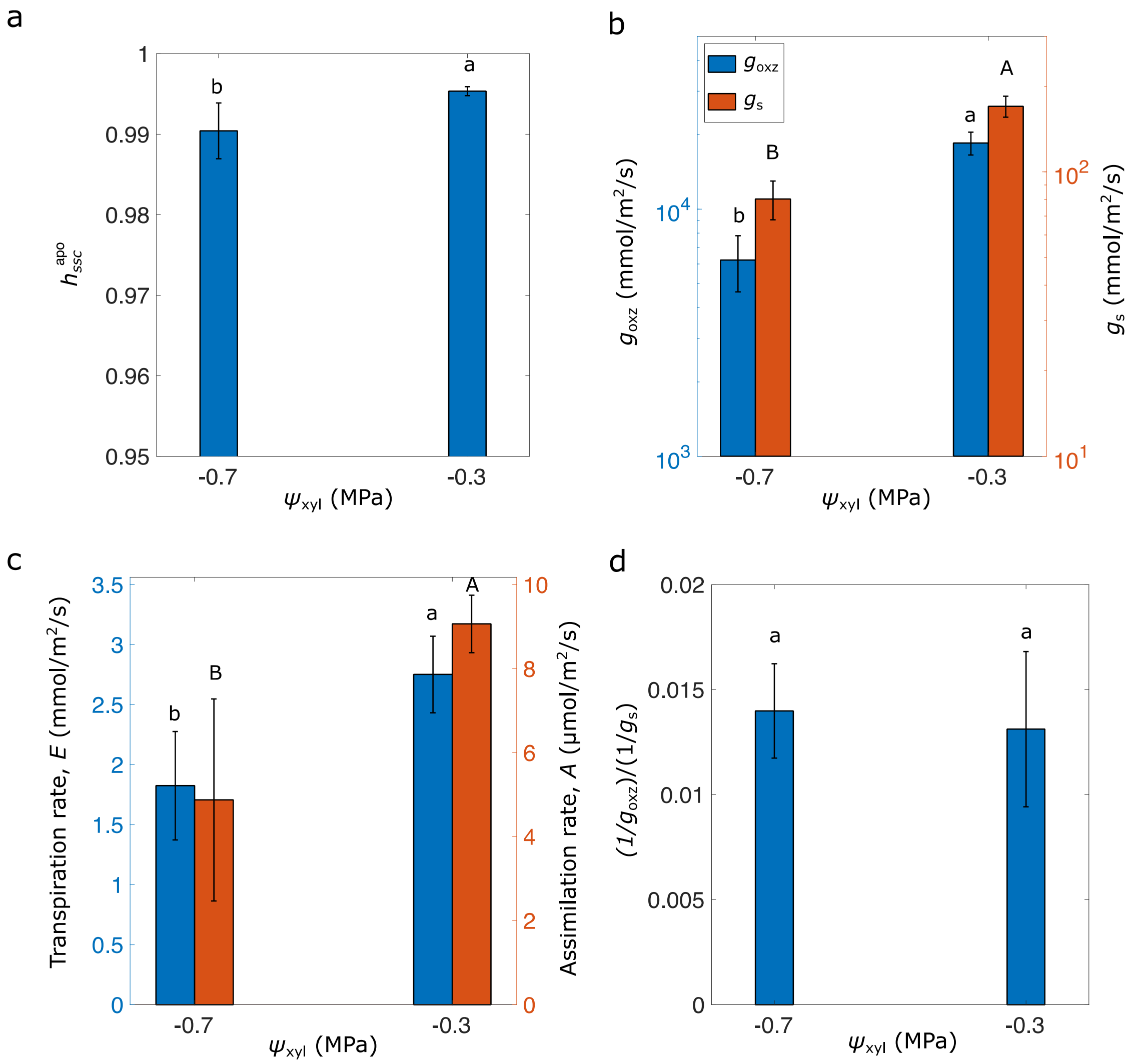

**Fig. S7. Loss of conductance of OXZ in a C3 species tomato (*Solanum Lycopersicum*).**
**a**. In tomato, internal relative humidity, $h_{\mathrm{ssc}}^{\mathrm{apo}}$ stays very near saturation ($h_{\mathrm{ssc}}^{\mathrm{apo}} \geq 0.99$) even as $g_{\mathrm{oxz}}$ drops ~ 3-fold (b) as it passes from its well-watered state to near TLP ($\psi_{\mathrm{TLP}} \cong -0.8$ MPa in tomato). Despite the drop in $g_{\mathrm{oxz}}$ with xylem stress, $g_{\mathrm{s}}/g_{\mathrm{oxz}}$ remains small (0.015) (d); no substantial undersaturation occurred (a) because the upstream resistance of the OXZ never competed with the that of the stomates.

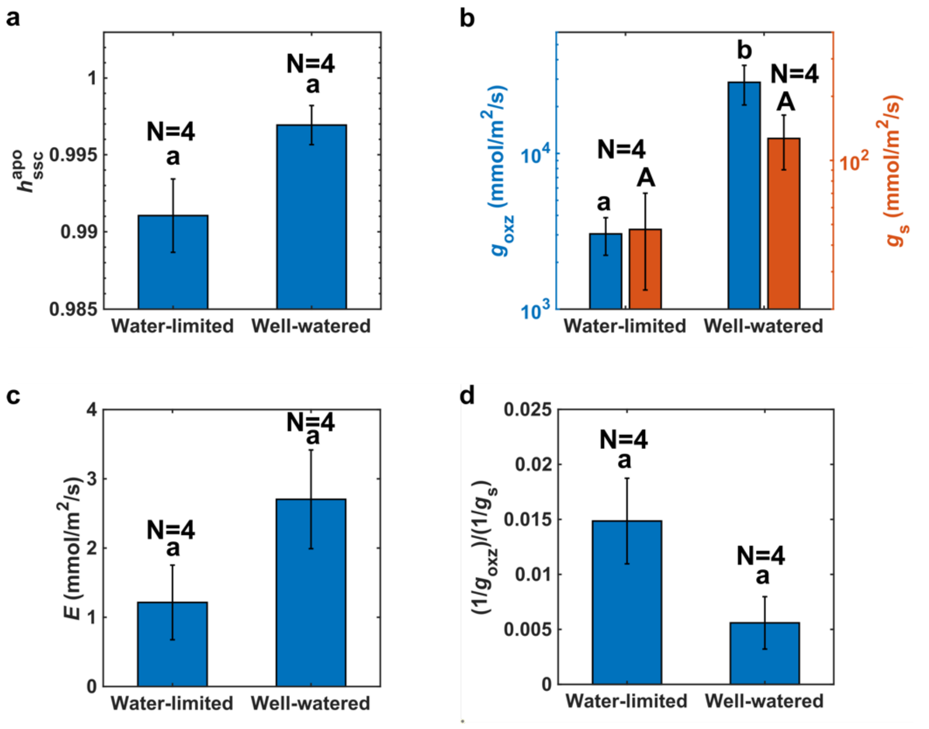

**Fig. S8. Loss of conductance of OXZ in a C3 species cotton (*Gossypium hirsutum*).**
**a**. In cotton, internal relative humidity, $h_{\rm ssc}^{\rm apo}$ stays very near saturation ($h_{\rm ssc}^{\rm apo} \geq 0.99$) even as $g_{\rm oxz}$ drops $\sim$ 9-fold (b) as it passes from its well-watered state ($\psi_{\rm xyl} > -0.5$ MPa) to water-limited state ($\psi_{\rm xyl} < -0.5$ MPa). Despite the drop in $g_{\rm oxz}$ with xylem stress, $g_{\rm s}/g_{\rm oxz}$ remains small (0.015) (d); no substantial undersaturation occurred (a) because the upstream resistance of the OXZ never competed with the that of the stomates.

**S7. Corrected stomatal conductance calculation based on chlorophyll fluorescence:**
We first estimate $c_{\rm i}$ using the biochemical model for photosynthesis as described in Caemmerer et al. [10] There are two different sets of equations that are needed to fit the $A - c_{\rm i}$ curve: (1) Enzyme limited rate equations and (2) Light and electron transport limited rate equations. Knowing the value of $A$ from gas exchange equipment, we compare whether that $A$ lies on the enzyme-limited regime or the light-limited regime using the A-ci curve and then use that specific set of equations to calculate $c_i$. In most of our experiments, our measurements are light-limited. Then, we use the standard equation used by gas exchange equipment to then calculate stomatal conductance, $g_s$, with a known value of $c_i$. In conventional scenario, this equation is used to calculate $c_i$ with a known $g_s$. This standard equation accounts for boundary layer and increase in mass flow rate of airstream due to $E$. Parameters related to enzyme kinetics are borrowed from literature and tabulated in Table S1.

Briefly, the enzyme limited equations are as follows:

$$A_c = min(V_p - R_{\rm m} + g_{bs}c_m, V_{\rm cmax} - R_{\rm d} \qquad \text{(S7)}$$

Where $\quad V_p = \min\left(\frac{V_{pmax}\, c_m}{c_m + K_p},\ V_{pr}\right)$

$$A_c = g_m(c_i - c_m) \tag{S8}$$

Using two equations and two unknowns in Eqs. S7 and S8, we estimate corrected stomatal conductance based on C4 biochemical model $c_{\mathrm{i}}$ and $c_{\mathrm{m}}$ from the Eqs. S7-S8 when the assimilation is enzyme limited.

The light limited equations are as follows:

$$A_j = \frac{\left(1-\frac{\Gamma^*}{c_s}\right)(1-x)J_t}{3(1+7\Gamma^*/3c_s)} - R_{\mathrm{d}} \tag{S9}$$

$$A_j = \frac{xJ_t}{2} + g_{bs}(c_s - c_m) - R_m \tag{S10}$$

$$A_j = g_m(c_i - c_m) \tag{S11}$$

We measured $J_t$, $A_j$, $c_a$ using PAM + Gas exchange and we solved for $c_s$, $c_m$ & $c_i$ using Eqs. S9-S11, when assimilation rate is light-limited. With a known $c_i$, we use Eq. S12 to estimate $g_{\mathrm{s}}^{\mathrm{C4-model}}$ using measured values of $A$ and $E$.

$$A_{\mathrm{measured}} = \frac{1}{\frac{1.585}{g_{\mathrm{s}}^{\mathrm{C4-model}}} + \frac{1.37}{r_{\mathrm{bl}}}} (C_a - C_i) - E\frac{C_i + C_a}{2} \tag{S12}$$

See Table S1 below for definitions and quantitative estimates for different variables used in the above equations.

As reported in Fig. S9, the strong correlation between corrected stomatal conductance using the C4 model of photosynthesis ($g_{\mathrm{s}}^{\mathrm{C4-model}}$) and corrections for unsaturation as measured using AquaDust ($g_{\mathrm{s}}^{\mathrm{corr}}$) across varying soil water stress conditions (Fig. S9a) validates both approaches for correcting stomatal conductance measurements. The error bars on the C4 model predictions, accounting for variations in mesophyll conductance (gm), net assimilation rate (A), and partitioning factor (x), suggest that the model is robust across a reasonable range of internal leaf gas exchange parameters. This agreement between two independent correction methods strengthens confidence in both approaches.

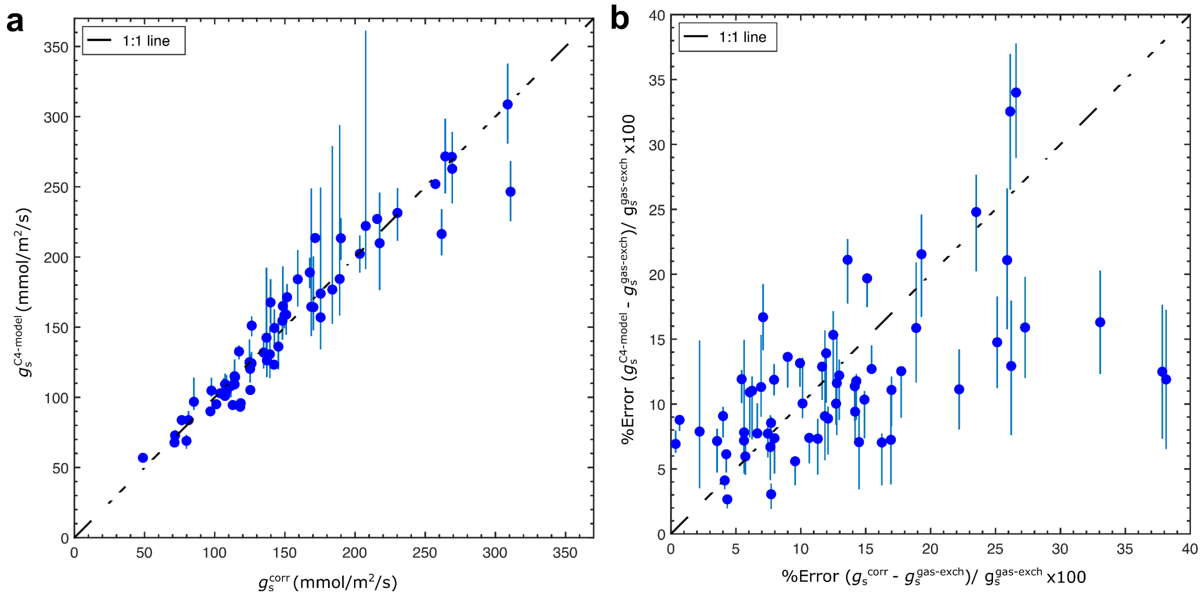

**Fig. S9. Comparison of corrected stomatal conductance using the C4 model of photosynthesis ($g_s^{\mathrm{C4-model}}$) and corrections for unsaturation as measured using AquaDust ($g_s^{\mathrm{corr}}$).**

**a**. Plot of corr ected stomatal conductance using C4 model predictions for stomatal conductance ($g_s^{\mathrm{C4-model}}$) in maize leaves compared with the corrected conductance based on unsaturation ($g_s^{\mathrm{corr}}$) reported by AquaDust under varying soil water stress (*VPD*=2 kPa; N = 26 biological replicates; $r^2$=0.87, RMSE=21.5mmol/m$^2$/s). Error bars on the C4 model predictions are calculated by varying mesophyll conductance ($g_\mathrm{m}$) from 0.6 to 1.4 times its nominal value, net assimilation rate (*A*) by ± 0.3 times, and partitioning factor *x* between 0.3 and 0.4. These uncertainties account for potential variabilities in internal leaf gas exchange parameters. **b**. Percentage error between the stomatal conductance estimated by gas exchange equipment using the conventional assumption of saturation ($g_s^{\mathrm{sat}}$), against $g_s^{\mathrm{C4-model}}$ and stomatal conductance estimate accounting for unsaturation as measured using AquaDust ($g_s$) shows the relative deviation of the corrected stomatal conductance from the Gaastra saturation model predictions across varying soil water stress ($r^2$=0.69, RMSE=8.1%).

**Table S1: Parameters used for $C_4$ biochemical model for photosynthesis to estimate $g_s$ based on carbon stream and chlorophyll fluorescence measurements.**

| Parameter | Value | Definition | Reference |
|---|---|---|---|
| $J_t$ | Measured using PAM Fluorescence | Total electron transport rate | |
| $V_{pmax}$ | 528 μmol m$^{-2}$ s$^{-1}$ | Maximum PEPC activity | Kolbe, Ph.D. Thesis (2018) |
| $V_{cmax}$ | 60 μmol m$^{-2}$ s$^{-1}$ | Maximum Rubisco activity | |
| $V_{pr}$ | 80 μmol m$^{-2}$ s$^{-1}$ | PEP regeneration rate | |
| $K_p$ | 15.4 Pa | Michaelis constant of PEPC for $CO_2$ | DiMario and Cousins (2019) |
| $g_{bs}$ | 0.003 $\mu$mol m$^{-2}$ s$^{-1}$ Pa$^{-1}$ | Bundle sheath conductance to $CO_2$ | Alonso-Cantabrana *et al.* (2018) |

| | | | |
|---|---|---|---|
| $R_d$ | 0.01×$V_{cmax}$ | Leaf mitochondrial respiration | Farquhar *et al.* (1980) |
| $R_m$ | 0.5 $R_d$ | Mesophyll mitochondrial respiration | |
| $x$ | 0.4 (optimal) | Partitioning factor of electron transport rate | |
| $g_m$ | Estimated by fitting the enzyme limited rate equations to A-ci curve | Mesophyll conductance to $CO_2$ | Ubierna *et al.* (2017) |
| $\Gamma^*$ | Estimated from A-ci curve, typically between 1-30 $\mu$mol/mol | $CO_2$ compensation point | |
| $C_s$ | | Bundle-sheath CO2 concentration | |
| $C_m$ | | Mesophyll CO2 concentration | |
| $C_i$ | | Intercellular CO2 concentration | |
| $C_a$ | | Cuvette atmospheric CO2 concentration | |
| $r_{bl}$ | 0.4 mmol $m^{-2}$ $s^{-1}$ | Boundary layer resistance – gas exchange equipment specific | |

**S8. Confocal imaging methods**

These methods relate to the data presented in Figure 3b of the main text.

**Experimental setup:**

An intact leaf from a slac1;2 null mutant (in a maize W22 inbred background) was infiltrated with AquaDust a day prior to imaging. Plants were sampled with a range of varying values of $\psi_{\text{xyl}}$ by withholding soil water supply above the turgor loss point. These intact leaves were placed under a laser scanning confocal microscope (Zeiss LSM880 upright) inside the cuvette head of a gas exchange equipment (CIRAS-3, PPSystems). We varied $VPD_{\text{leaf}}$ locally by controlling the relative humidity of the circulating air stream inside the gas exchange cuvette. Imaging was performed with a Long Working Distance 20x objective (Olympus LMPlanFL N 20x).

**Extracting AquaDust-reported water potentials pixel-by-pixel from confocal images (corresponding to main text Figure 3):**

In confocal sections within the plane of the leaf (Fig. 3c) – x-y images are collected at z = 0, 6, 12, and 18 µm beneath the inner surface of the epidermis. The autofluorescence signal provided information on the morphology of the cells to assess their state of turgor; the AquaDust emission allowed assessment of water status at each pixel based on the FRET efficiency ($\zeta(\psi)$). For every location on the leaf imaged with the confocal microscope, two images are collected. In the first image, the sample was excited with a 561 nm laser (close to acceptor dye excitation maximum) and emission from 575 nm – 700 nm was collected in fourteen 9 nm wide channels. This acceptor dye emission image was a first filter that allowed us to locate AquaDust pixels in the image. In the second image, the sample was excited with a 488 nm laser and emission from 495 nm – 700 nm was collected in twenty-two 9 nm wide channels. The second image served two roles: we took advantage of our knowledge of the spectral signature of AquaDust to test if the pixels that passed the first filter satisfied the spectral signature test as well. For the set of pixels ultimately selected as AquaDust pixels, the FRET efficiency ($\zeta$) was calculated on a pixel-by-pixel basis. The conversion from $\zeta$ to water potential ($\psi$) used the calibration dataset described in the next paragraph.

**Calibration of AquaDust signal under a confocal microscope**

Calibration of AquaDust fluorescence intensity as a function of $\psi_{\mathrm{xyl}}$ was performed by collecting AquaDust measurements and corresponding Scholander Pressure Chamber measurement ($\psi_{leaf}^{SPC}$) in leaves with various degrees of soil water stress. Blocking the transpiration from leaves enabled local equilibrium of leaf tissue being imaged with AquaDust and $\psi_{\mathrm{xyl}}$. We achieved this by placing an oil droplet over the maize leaf and imaged it using Apochromat 20x/1.0 oil immersion objective. We obtained the fluorescence intensity from AquaDust at different depths by exciting the donor dye with 488 nm laser line and collecting the emission in the Lambda imaging mode within a range of 500 – 700 nm with a laser scanning confocal microscope (Zeiss LSM880). Pixels in the image corresponding to AquaDust nanoreporter were isolated by thresholding based on fluorescence intensity obtained by directly exciting acceptor dye with 568 nm laser line. The average spectra were constructed as shown in **Figs. S9-S11**. Using the fluorescence intensity collected in 9 nm bins from 500 nm to 700nm, we calculate the FRET Efficiency at each of these pixels using the following equation:

$$\boldsymbol{\zeta(i,j)} = \frac{\boldsymbol{I_{488}^{575}(i,j) - 0.09\left(I_{488}^{520}(i,j)\right)}}{\boldsymbol{I_{488}^{520}(i,j) + I_{488}^{575}(i,j) - 0.09\left(I_{488}^{520}(i,j)\right)}} \qquad \text{(S13)}$$

Where, $\zeta(i,j)$ is the FRET Efficiency, $I_{488}^{575}(i,j)$ is the fluorescence emission at $575 \pm 4.5$ nm with excitation using 488 nm laser line, and $I_{488}^{520}(i,j)$ is the fluorescence emission at $520 \pm 4.5$ nm with the excitation using 488 nm laser line at pixel corresponding to $(x,y) = (i,j)$ in the 2D plane. The subtraction of 9% of intensity from $I_{488}^{575}(i,j)$ accounts for cross-bleeding of donor (Oregon Green 488) fluorescence at the 575 nm.

The corresponding water potential, $\psi(i,j)$ is estimated from $\zeta(i,j)$ using 1:1 monotonic function that relate these two variables by accounting for gel swelling using modified Flory-Rehner model and dipole-plane FRET model for donor-acceptor interaction. Details of the model are described in Jain et al.[1]

After calibration using dark-adapted, non-transpiring leaves, we proceed to collect measurements in transpiring leaves under confocal microscope. Leaves of slac1-2 null mutants were infiltrated with AquaDust. We took advantage of the lack of stomatal closure in slac1-2 null mutant plants in the dark, allowing large, steady-state transpiration rates during the image acquisition ($E$ ranges from $4 - 8$ mmol/m$^2$/s). Plants were sampled with a range of values of $\psi_{\mathrm{xyl}}$ by withholding soil water supply above the turgor loss point. These intact leaves were placed under the confocal microscope inside the cuvette head of a gas exchange equipment (Walz GFS-3000 gas exchange system, Heinz Walz GmbH). We varied

$VPD_{\mathrm{leaf}}$ locally by controlling the relative humidity of the circulating air stream inside the gas exchange cuvette (Fig. S13-S15). To examine the recovery of water potential in absence of transpiration, each leaf was cut until transpiration dropped below ~1 mmol/m$^2$/s (Fig. S16). For constant $\psi_{\mathrm{xyl}}$, we observe linear decline in $h_{\mathrm{ssc}}^{\mathrm{apo}}$ with increasing $VPD_{\mathrm{leaf}}$ and $E$ demonstrating no significant change in $g_{\mathrm{oxz}}$ (slope) in response to $VPD_{\mathrm{leaf}}$, and complete recovery of $h_{\mathrm{ssc}}^{\mathrm{apo}}$ to $TLP$ in response to lack of transpiration after the leaf was cut (Fig. S17).

**Circularity and cell size estimation:**

These methods relate to the data presented in Figure 3e of the main text. Cell circularity and size measurements were conducted using ImageJ (Version 1.53) through three independent manual outlining of individual cells with the freehand selection tool, followed by spline fitting to ensure smooth boundary delineation. Cell area was directly computed from the pixel counts within each outlined region, with spatial calibration based on the image scale (pixels/µm). Circularity (C) was calculated using the relationship $C = 4\pi(A/P^2)$, where A [m$^2$] represents the cell area and P [m] the perimeter of the spline-fitted outline. This dimensionless parameter ranges from 0 to 1, with 1 indicating a perfect circle and lower values reflecting increasingly irregular shapes. This methodology provides a quantitative framework for assessing both cell morphology and size distribution, though it inherently assumes that the 2D projections adequately represent the cellular geometry. The spline-fitting procedure may introduce minor deviations from the true cell boundaries, but these are generally negligible compared to the errors that would be introduced by using unsmoothed, freehand outlines (ImageJ Documentation, 2024).

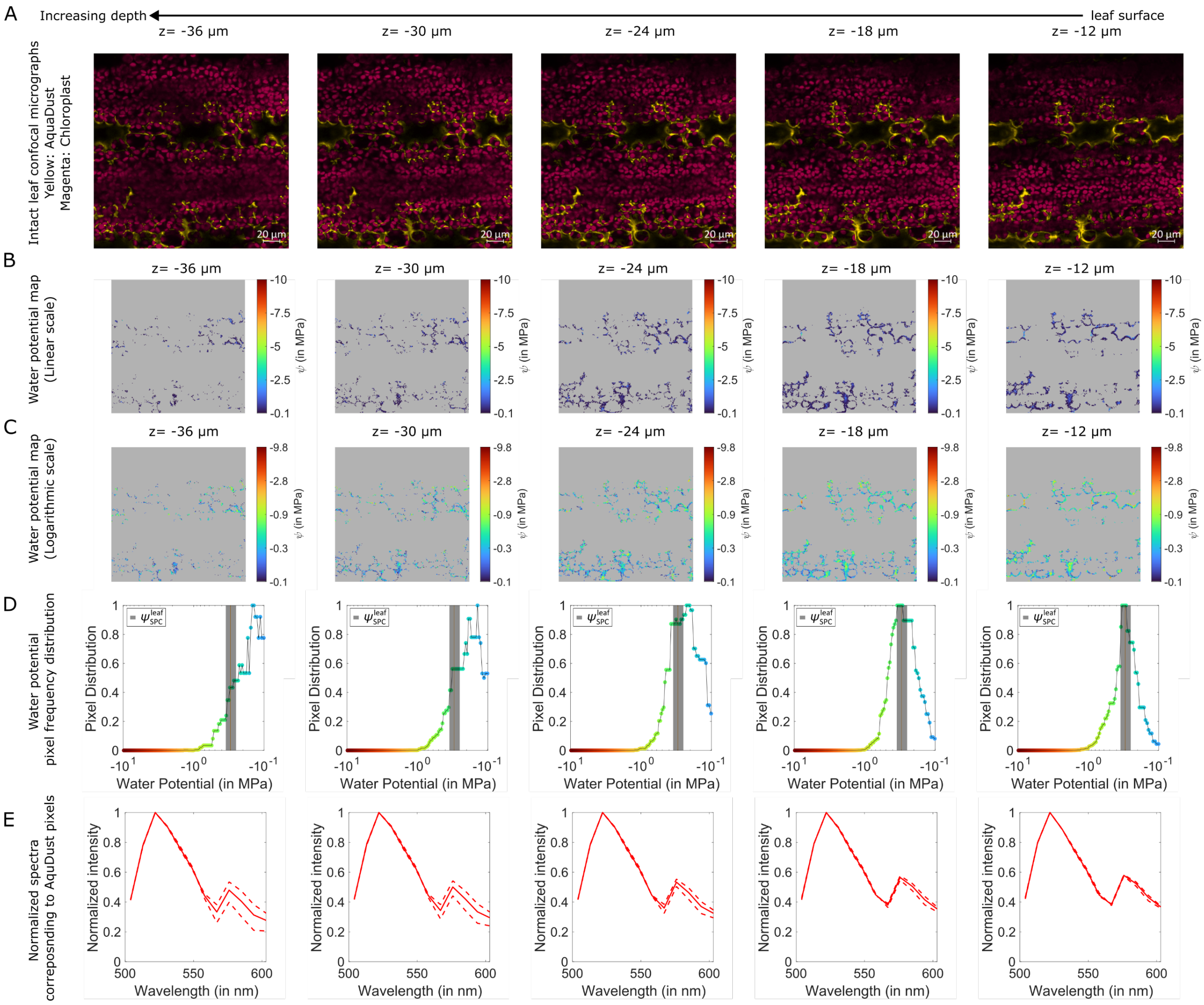

**Figure S10. Calibration of AquaDust measurements under confocal microscope against pressure chamber $\psi_{leaf}^{SPC} = -0.3$ MPa**

(A) Confocal micrographs of the maize leaf with a top-view across varying leaf depths. AquaDust signal is false colored as yellow and chloroplast signal is false colored as magenta. Thresholding is used to segment AquaDust and non-AquaDust pixels and corresponding water potential is calculated as described by Eq. S1. Water potential is shown as colormap in (B) linear scale and (C) logarithmic scale for clarity. (D) Frequency distribution of pixels at different water potential values, peak corresponds to the mode of the distribution. (E) Bold line and dashed lines represent average and standard deviation of spectra respectively normalized to emission corresponding to the peak emission from the donor dye.

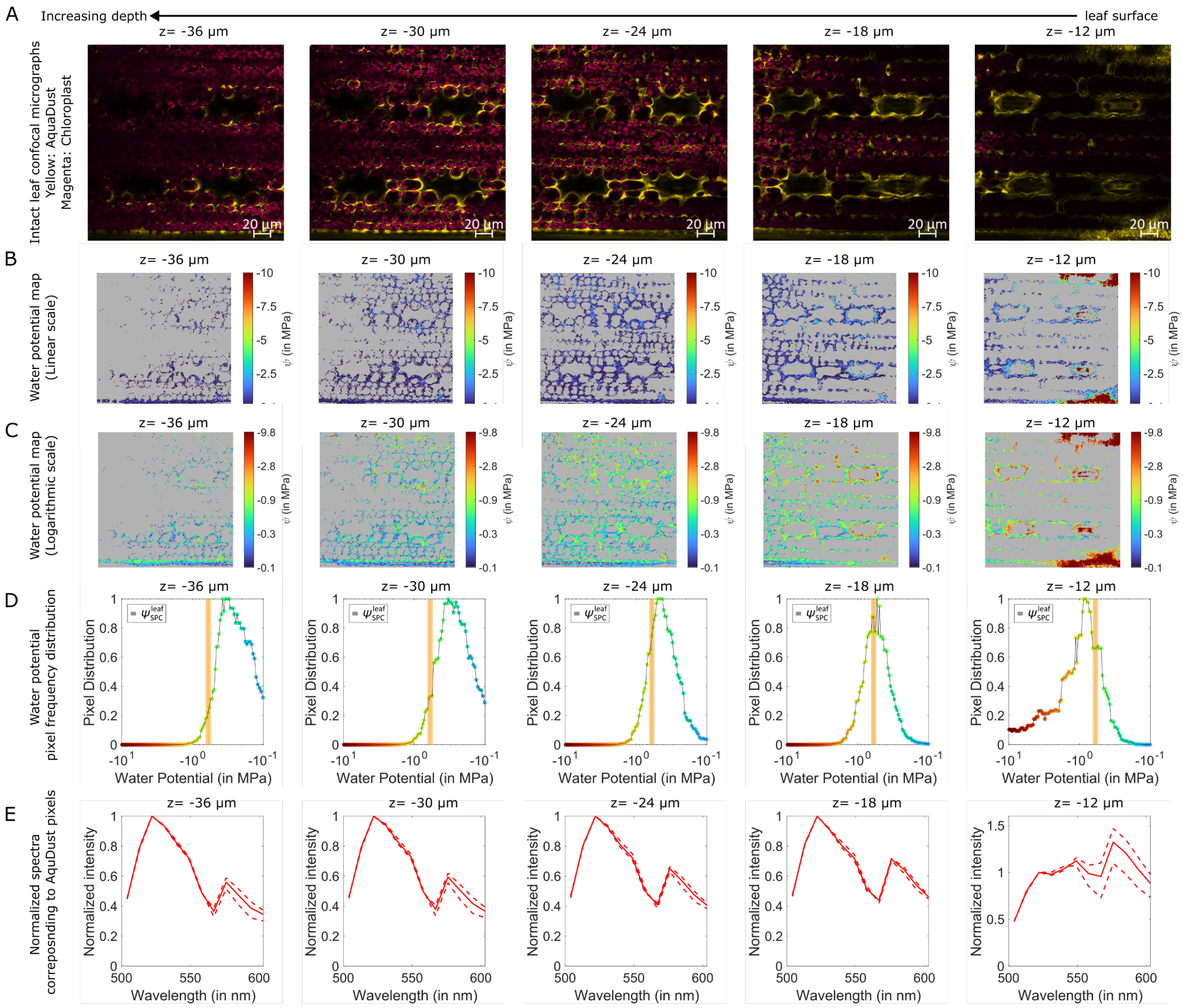

**Figure S11. Calibration of AquaDust measurements under confocal microscope against pressure chamber $\psi_{leaf}^{SPC} = -0.6$ MPa.**

(A) Confocal micrographs of the maize leaf with a top-view across varying leaf depths. AquaDust signal is false colored as yellow and chloroplast signal is false colored as magenta. Thresholding is used to segment AquaDust and non-AquaDust pixels and corresponding water potential is calculated as described by Eq. S1. Water potential is shown as colormap in (B) linear scale and (C) logarithmic scale for clarity. (D) Frequency distribution of pixels at different water potential values, peak corresponds to the mode of the distribution. (E) Bold line and dashed lines represent average and standard deviation of spectra respectively normalized to emission corresponding to the peak emission from the donor dye.

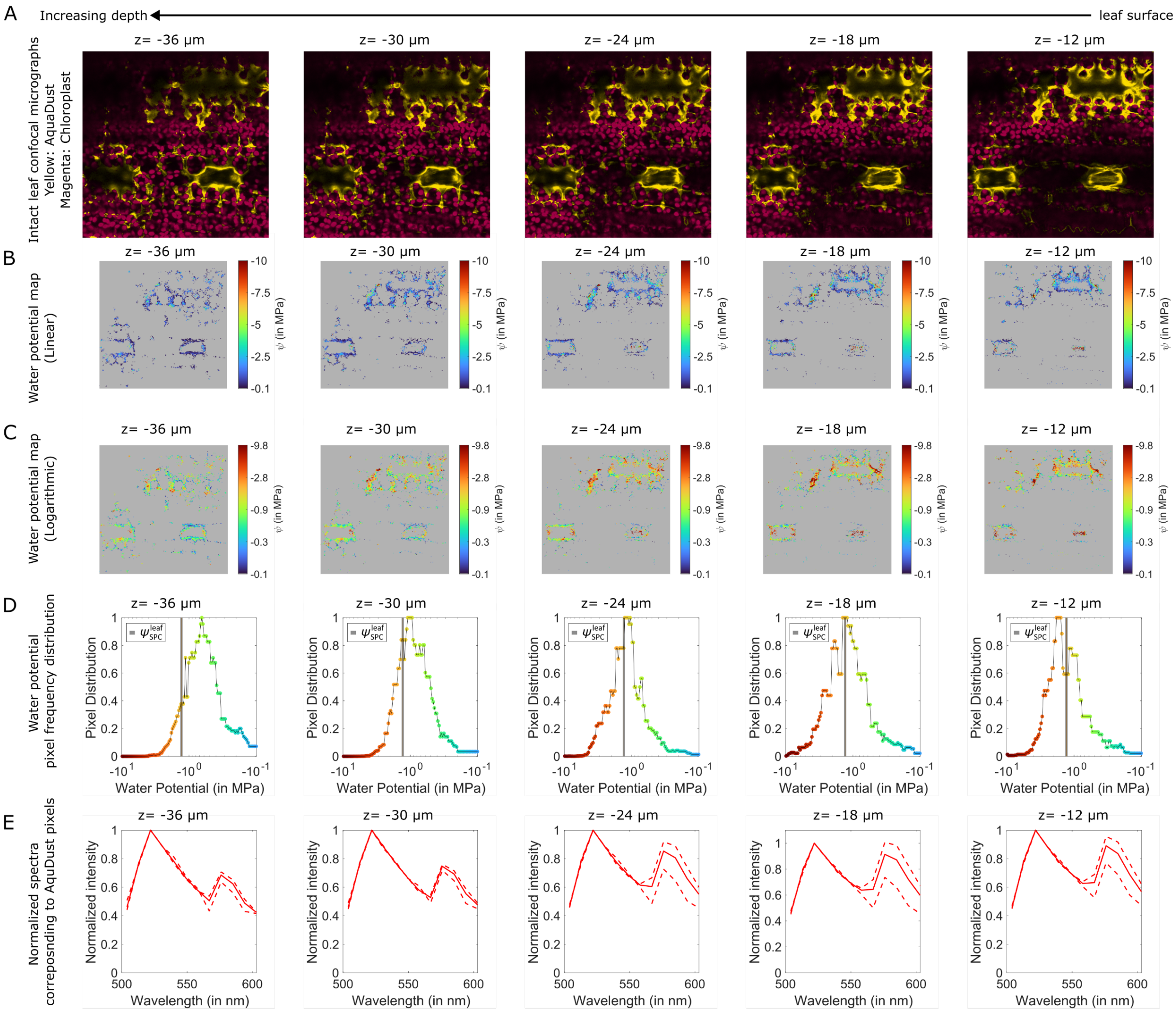

**Figure S12. Calibration of AquaDust measurements under confocal microscope against pressure chamber $\psi_{leaf}^{SPC} = -1.3$ MPa.**

(A) Confocal micrographs of the maize leaf with a top-view across varying leaf depths. AquaDust signal is false colored as yellow and chloroplast signal is false colored as magenta. Thresholding is used to segment AquaDust and non-AquaDust pixels and corresponding water potential is calculated as described by Eq. S1. Water potential is shown as colormap in (B) linear scale and (C) logarithmic scale for clarity. (D) Frequency distribution of pixels at different water potential values, peak corresponds to the mode of the distribution. (E) Bold line and dashed lines represent average and standard deviation of spectra respectively normalized to emission corresponding to the peak emission from the donor dye.

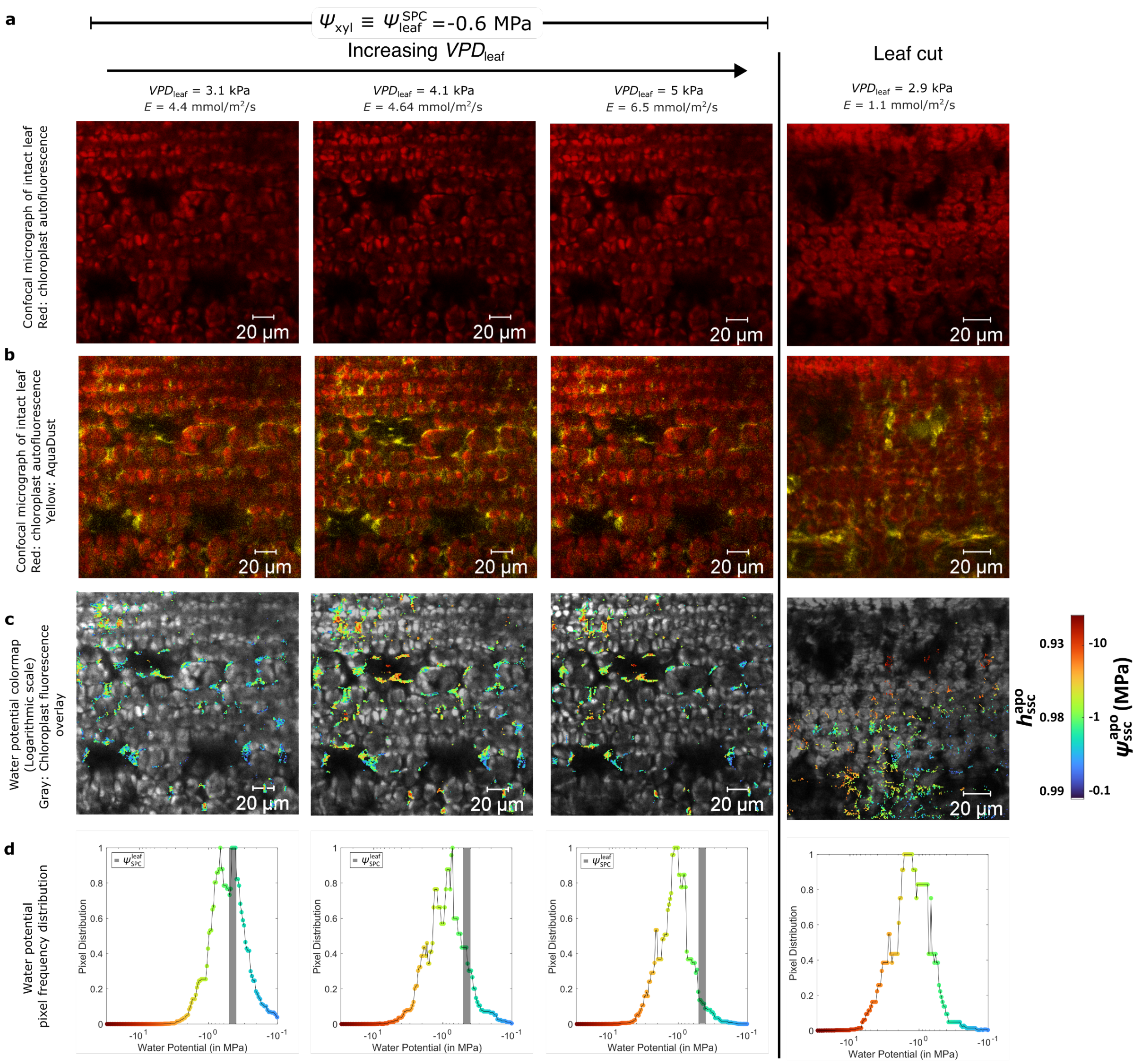

**Figure S13. Cellular scale measurement of $\psi_{ssc}^{apo}$ at constant depth of $z = -24$ µm in response to constant $\psi_{xyl} = -0.6$ MPa and varying $VPD_{leaf}$ and the leaf is cut at the end to force stomatal closure and equilibrium of the tissue.**

(A) Autofluorescence from chloroplast. (B) AquaDust is false colored as yellow and chloroplast is false colored as magenta. Thresholding is used to segment AquaDust and non-AquaDust pixels and corresponding water potential is calculated as described by Eq. S1. (C) Water potential is shown as color map in logarithmic scale with chlorophyll fluorescence as gray overlay to show cell boundaries. (D) Frequency distribution of pixels at different water potential values, peak corresponds to the mode of the distribution.

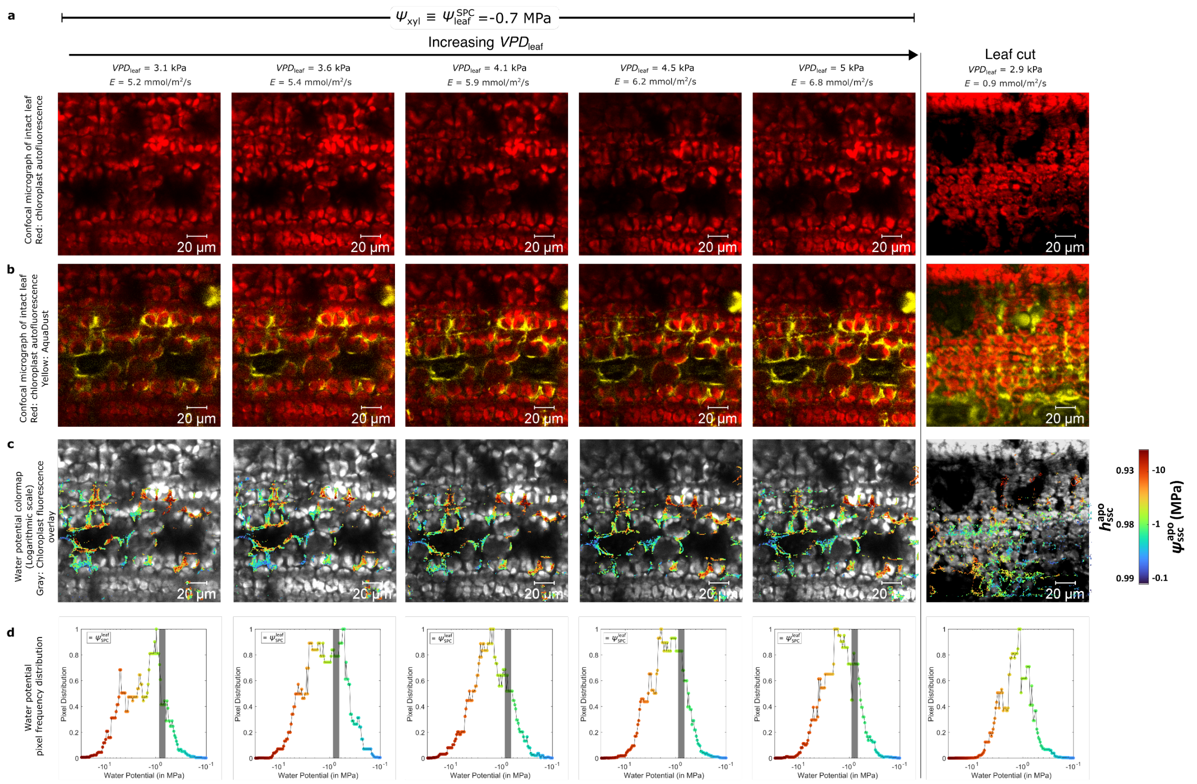

**Figure S14: Cellular scale measurement of $\psi_{ssc}^{apo}$ at constant depth of $z = -24$ µm in response to constant $\psi_{xyl} = -0.75$ MPa and varying $VPD_{leaf}$ and the leaf is cut at the end to force stomatal closure and equilibrium of the tissue.**

(A) Autofluorescence from chloroplast. (B) AquaDust signal is false colored as yellow and chloroplast signal is false colored as magenta. Thresholding is used to segment AquaDust and non-AquaDust pixels and corresponding water potential is calculated as described by Eq. S1. (C) Water potential is shown as colormap in logarithmic scale with chlorophyll fluorescence as gray overlay to show cell boundaries. (D) Frequency distribution of pixels at different water potential values, peak corresponds to the mode of the distribution.

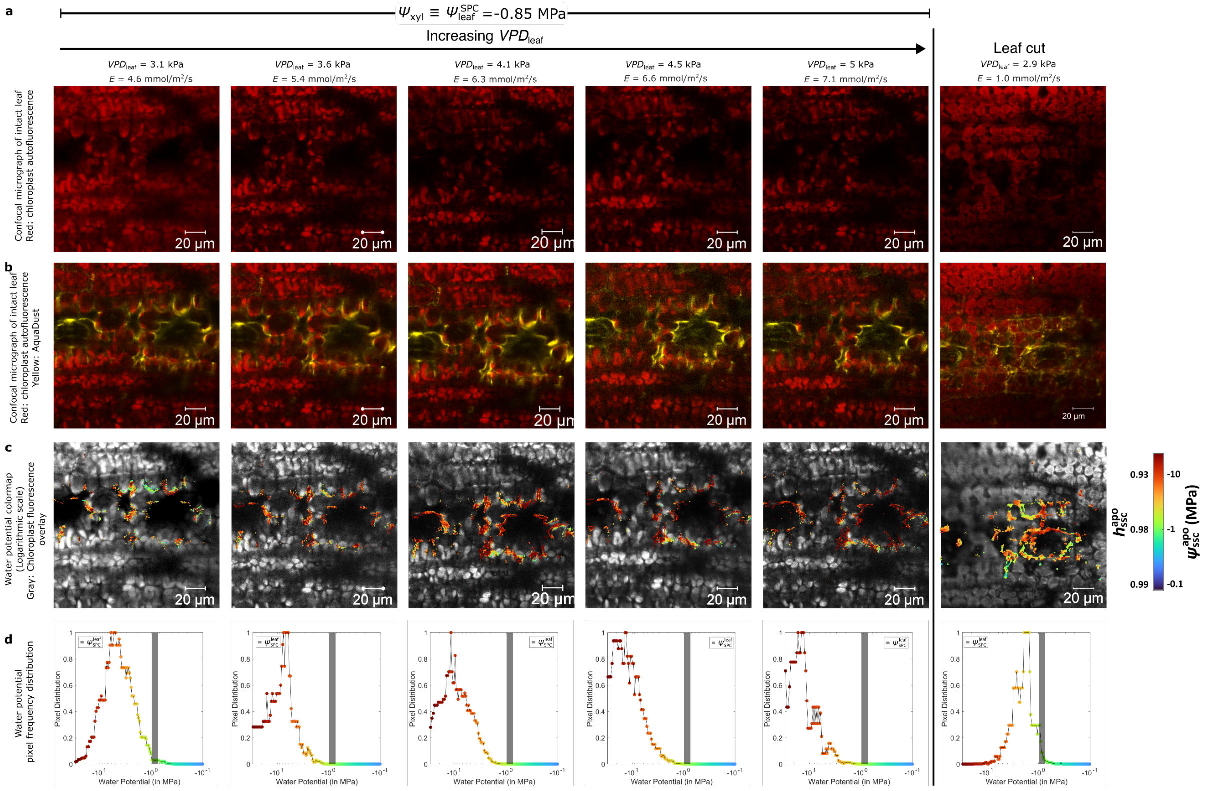

**Figure S15: Cellular scale measurement of $\psi_{ssc}^{apo}$ at constant depth of $z = -24$ µm in response to constant $\psi_{xyl} = -0.85$ MPa and varying $VPD_{leaf}$ and the leaf is cut at the end to force stomatal closure and equilibrium of the tissue.**

(A) Autofluorescence from chloroplast. (B) AquaDust signal is false colored as yellow and chloroplast signal is false colored as magenta. Thresholding is used to segment AquaDust and non-AquaDust pixels and corresponding water potential is calculated as described by Eq. S1. (C) Water potential is shown as color map in logarithmic scale with chlorophyll fluorescence as grey overlay to show cell boundaries. (D) Frequency distribution of pixels at different water potential values, peak corresponds to the mode of the distribution.

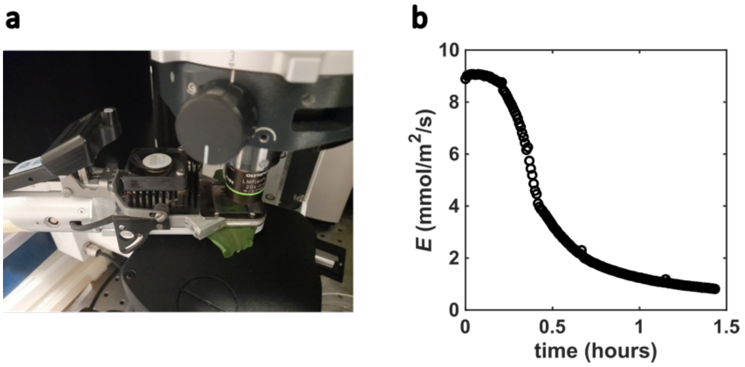

**Figure S16. Leaf rolling and gas exchange data in a leaf cut under confocal microscope.**

(a) Leaf was cut at time=0 hours. (a) Leaf rolling observed at time∼ 1 hour. (b) Transpiration rate was measured continuously as a function of time and measurements corresponding to cut leaf case, reported in Fig. S13-S15, were performed at time∼ 1 hour.

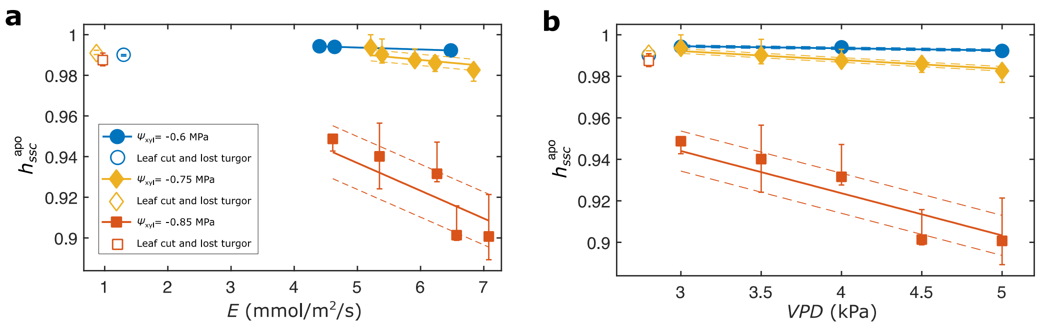

**Figure S17. Gas exchange data in a leaf cut under confocal microscope.**
Relationship between internal undersaturation ($h_{\mathrm{ssc}}^{\mathrm{apo}}$) as extracted from the peak of pixel distribution shown in Fig. S13-S15 with error bars representing the standard error of water potential from all pixels plotted against transpiration rate ($E$) for $\psi_{\mathrm{xyl}} = -0.6$ MPa ($h_{\mathrm{ssc}}^{\mathrm{apo}} = 1 - 0.001E,\ R^2 = 0.99$), $\psi_{\mathrm{xyl}} = -0.75$ MPa ($h_{\mathrm{ssc}}^{\mathrm{apo}} = 1 - 0.003E,\ R^2 = 0.68$), and $\psi_{\mathrm{xyl}} = -0.85$ MPa ($h_{\mathrm{ssc}}^{\mathrm{apo}} = 1 - 0.014E,\ R^2 = 0.74$); (h) vapor pressure deficit for $\psi_{\mathrm{xyl}} = -0.6$ MPa ($h_{\mathrm{ssc}}^{\mathrm{apo}} = 1 - 0.001E,\ R^2 = 0.85$), $\psi_{\mathrm{xyl}} = -0.75$ MPa ($h_{\mathrm{ssc}}^{\mathrm{apo}} = 1 - 0.004E,\ R^2 = 0.95$), and $\psi_{\mathrm{xyl}} = -0.85$ MPa ($h_{\mathrm{ssc}}^{\mathrm{apo}} = 1 - 0.020E,\ R^2 = 0.86$).

### S9. Measurement of bulk leaf osmotic potential in response to drought stress of water transport in the outside-xylem zone.

We evaluated osmotic potential ($\boldsymbol{\psi}_{\mathbf{osm}}$) through destructive osmolality measurements using a Wescor Vapro 5520 osmometer (Wescor Inc., Logan, UT) on leaf tissue samples. Measurements were conducted on 21 maize plants each for WW and WL case, with tissue sampling systematically performed across the tip of the growing leaf. The osmometer was calibrated using standard solutions prior to each measurement series to ensure measurement accuracy and stability. Tissue samples were collected coincident with continuous environmental monitoring via datalogger systems and AquaDust-based water potential measurements ($\boldsymbol{\psi}$). Each 5-cm long leaf tissue sample was immediately stored in a 20 mL plastic syringe at -18°C. Samples were thawed at room temperature two hours before measurement. A droplet of leaf sap was pressed out of the syringe onto a filter paper, which was inserted in the osmometer. The osmometer output (c, in mmol kg-1) was then converted to osmotic potential using the van’t Hoff equation:

$$\psi_{osm} = -icRT$$

Where i is the van’t Hoff factor (set to 1), R is the universal gas constant (8.31 J mol-1 K-1) and T the leaf temperature at the time of sampling (~303 K).

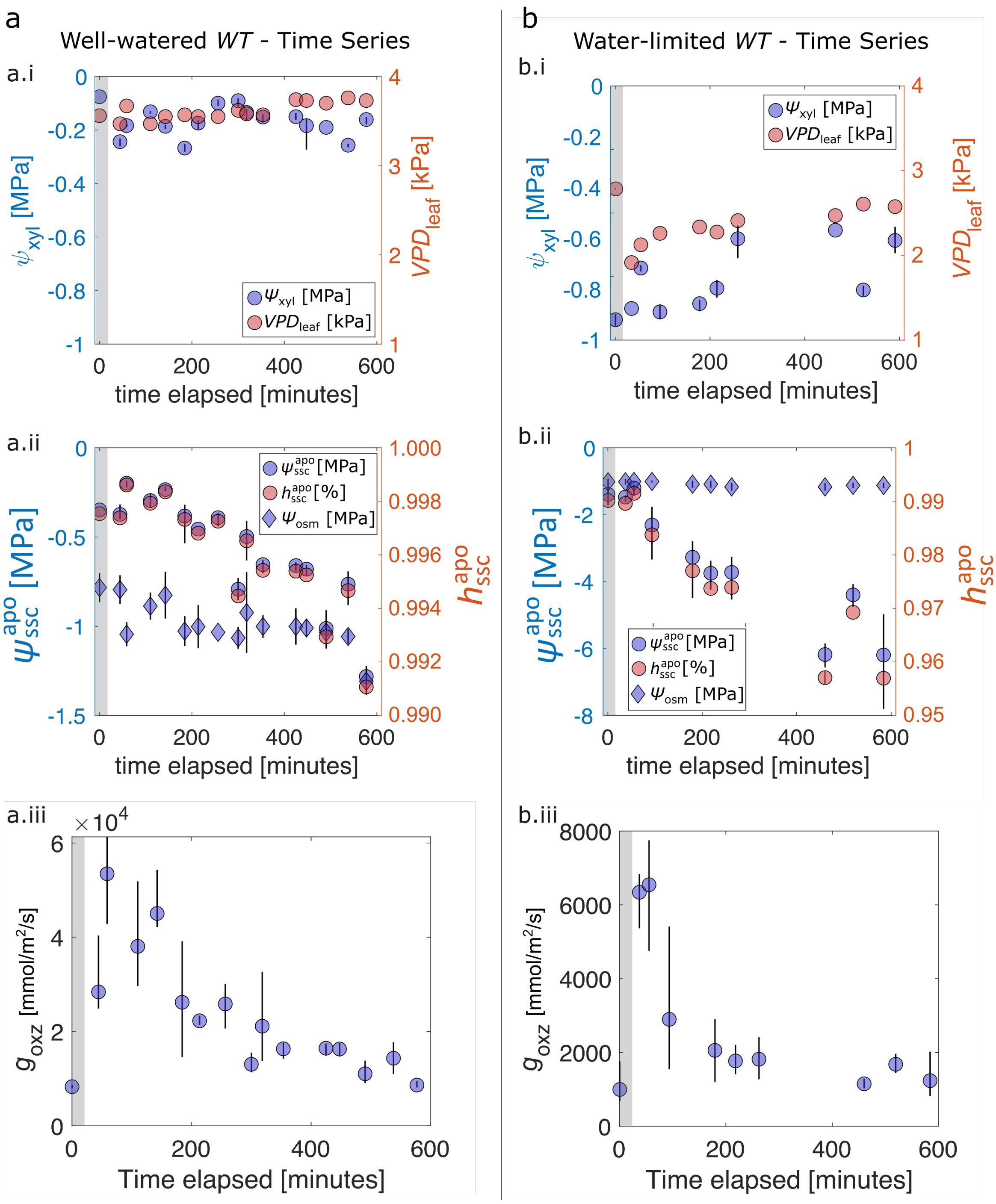

**Figure S18. Changes in bulk leaf osmotic potential in response to drought stress.** Diurnal changes in osmotic potential of maize leaf tissue with varying drought stress: For two cases, (A) with well-watered wild-type ($\boldsymbol{WT}$) plants (high $\boldsymbol{\psi_{xyl}}$) and (B) water-limited wild-type ($\boldsymbol{WT}$) plants (low $\boldsymbol{\psi_{\mathrm{xyl}}}$) and with relatively constant $\boldsymbol{VPD_{\mathrm{leaf}}}$, we report changes in $\boldsymbol{\psi_{apo}^{ssc}}$, $\boldsymbol{h_{apo}^{ssc}}$, $\boldsymbol{\psi_{\mathrm{osm}}}$, and corresponding $g_{\mathrm{oxz}}$. At t=0, plants were in dark (pre-dawn conditions, 6 AM, grey translucent band).

**S10. Model of water transport in the outside-xylem zone**

**Model description**

Here, we present a mathematical model of the steady transfer of transpiration through the OXZ and stomates. Qualitative and quantitative aspects of the model are informed by the main text Figs. 2 and 3 and we have used the model to make the predictions of variable conductance of the OXZ and of profiles of water status within this zone in Fig. 4. Relative to state-of-the-art models of the water relations of the OXZ,[11,12] this model explicitly accounts for a cell-to-cell path of liquid flow alongside a distinct apoplasmic path. This feature allows it to accommodate the disequilibrium documented in main text Fig. 3 between symplasm and apoplasm.

**Anatomy of the outside-xylem zone (OXZ) and modeled geometry.**
**Figure S19** provides an overview of our model for maize. For this study, we have only considered this maize-like architecture with adaxial-abaxial symmetry. We will consider generalizations to other architectures in future studies. We model the transfer of water through the outside xylem zone (OXZ) in a characteristic "repeat unit" shown as solid white square outlines superimposed on a confocal section in the plane of a maize leaf in Fig. **S19a**. The lateral extent ($A_{\text{tot}}$ [m$^2$] in $x$-$y$ plane) of a repeat unit spans from the mid-plane of one vessel to the mid-plane of an adjacent vessel and contains the mesophyll and bundle sheath cells and intercellular air spaces (IAS) through which transpired water flows from the xylem to one or more stomates. The lateral area of the repeat unit, $A_{\text{tot}}$ defines the area over which we evaluate the rate of transpiration from the unit based on macroscopic values of stomatal conductance, $g_{\text{s}}$ [mmol/m$^2$/s] with its standard scaling to surface area of leaf. We do not model the diffusion through discrete stomates explicitly.

**Fig. S19b** presents a 3-D schematic diagram of a quarter section of a repeat unit of the OXZ that informs our pseudo 1-D model (**Fig. S19c**). We account for two coupled axial paths (along $z$): a cell-to-cell path with area, $A_{\text{c}-\text{c}}$ within a repeat unit through which liquid water flows via symplasmic, transcellular, cell wall, or a combination of these modes; and an apoplasmic path with area, $A_{\text{apo}}$ within a repeat unit through which water vapor diffuses through the IAS and liquid water flows through the cell wall. In each of these paths through the OXZ, we distinguish a bundle sheath zone around the mid-plane of thickness, $L_{\text{bs}}$ [m] and mesophyll zones beneath the adaxial and abaxial surfaces, each of thickness, $(L_{\text{tot}} - L_{\text{bs}})/2$. The geometric, hydraulic, and mechanical parameters of our model can be distinct between these zones. Given the adaxial-abaxial symmetry of maize leaves, we represent the adaxial ($z > 0$) and abaxial ($z < 0$) halves of the leaves as mirror images of one another with the same biophysical parameters for corresponding paths and zones. The interface between the cell-to-cell path and IAS (green dashed line and expanded view) has a perimeter, $\mathcal{P}_{\text{int}}$ [m] and represents the plasma membrane and cell wall.

Fig. S19c presents our pseudo 1-D approximate representation of transport in the OXZ. In this representation, water status varies only in the through-leaf direction (water potential, $\psi(z)$ or relative humidity, $h(z) = w(z)/w_{\text{sat}}$) in the cell-to-cell domain and the apoplasmic domain. This representation accounts for lateral gradients and flow from the cell-to-cell domain to the apoplasmic domain through their interface (green dashed line) while neglecting lateral gradients and flows (in $x$ and $y$) in each of these domains. This type of approximation ("fin analysis"[13]) is appropriate for coupling of long-narrow conduits in which internal conductance is large compared to the conductance between the conduits. We check this approximation below.

We account for the actual, 3-D internal geometry of the vapor path with three geometric parameters: first, the porosity of the leaf with respect to internal air spaces, $\phi_{\mathrm{vap}}$ [-]:

$$\phi_{\mathrm{vap}} = \frac{V_{\mathrm{vap}}}{V_{\mathrm{tot}}} \cong \frac{V_{\mathrm{apo}}}{V_{\mathrm{tot}}} \quad \text{(S14)}$$

where $V_{\mathrm{vap}}$ [m$^3$] is the volume of the IAS and $V_{\mathrm{tot}} = A_{\mathrm{tot}}L_{\mathrm{tot}}$ is the total volume of the repeat unit. In the second equality, we neglect the contribution of the cell walls to the volume of the apoplasm. This porosity has been measured and approximated by various methods.[14,15]

Second, we account for the tortuosity, $\tau_{\mathrm{apo}}$ [-] of the apoplasmic domain:

$$\tau_{\mathrm{apo}} = \left(\frac{L_{\mathrm{apo}}^{\mathrm{actual}}}{L_{\mathrm{tot}}}\right)^2 \text{ and } \tau_{\mathrm{c-c}} = \left(\frac{L_{\mathrm{c-c}}^{\mathrm{actual}}}{L_{\mathrm{tot}}}\right)^2 \quad \text{(S15)}$$

where $L_{\mathrm{apo}}^{\mathrm{actual}}$ [m] and $L_{\mathrm{c-c}}^{\mathrm{actual}}$ [m] are the local, non-linear path lengths through the IAS and cell-to-cell path, and $L_{\mathrm{tot}}$ [m] is the geometric (straight line) axial distance of the path.[14,15] We introduce $\tau_{\mathrm{c-c}}$ here to assess the geometry of the cell-to-cell path but do not account for it explicitly in our calculations; its contribution is included in the conductivity of this domain. We do use $\tau_{\mathrm{vap}}$ in our calculations and allow $\tau_{\mathrm{vap}}$ and $\phi_{\mathrm{vap}}$ to take on different values in the bundle sheath and mesophyll zones (see Table S4).

Third is a ratio of the area of the interface between the cell-to-cell and apoplasmic paths, $A_{\mathrm{int}}$ to that of the repeat unit, $A_{\mathrm{tot}}$:

$$f = \frac{A_{\mathrm{int}}}{A_{\mathrm{tot}}} = \frac{\mathcal{P}_{\mathrm{int}}L_{\mathrm{actual}}}{A_{\mathrm{tot}}} = \frac{\mathcal{P}_{\mathrm{int}}L_{\mathrm{tot}}\sqrt{\tau_{\mathrm{apo}}}}{A_{\mathrm{tot}}} \quad \text{(S16)}$$

This ratio has been estimated independently and provides a basis to evaluate the microstructure of the cell-to-cell and apoplasmic paths through the OXZ. We take its value, $f \cong 10 - 13$.[16] In Eq. S16, $\mathcal{P}_{\mathrm{int}}$ [m] is the perimeter of the interface.

Assuming the pores of the IAS are principally oriented along $z$, we estimate the total cross-sectional areas of the vapor path and cell-to-cell paths in a repeat unit as:

$$A_{\mathrm{vap}} \cong \frac{\phi_{\mathrm{vap}}}{\sqrt{\tau_{\mathrm{apo}}}}A_{\mathrm{tot}} \text{ and } A_{\mathrm{c-c}} \cong \frac{1-\phi_{\mathrm{vap}}}{\sqrt{\tau_{\mathrm{c-c}}}}A_{\mathrm{tot}} \quad \text{(S17)}$$

We evaluate the cross-sectional area of the cell wall, $A_{\mathrm{cw}}$ for axial flow of water as:

$$A_{\mathrm{cw}} = d_{\mathrm{cw}}\mathcal{P}_{\mathrm{int}} \quad \text{(S18)}$$

where $d_{\mathrm{cw}}$ [m] is the thickness of the cell wall.

Treating the local apoplasmic and cell-to-cell paths as having circular cross-sections (shown in Fig. S19b for apoplasm), we estimate their typical lateral dimensions, $d_{\mathrm{apo}}$ [m] and $d_{\mathrm{c-c}}$ [m], and of the local perimeter of the interface between domains, $\mathcal{P}_{\mathrm{int}}$ [m] as:

$$d_{\mathrm{apo}} = \frac{4\phi_{\mathrm{vap}}L_{\mathrm{tot}}}{f};\ d_{\mathrm{c-c}} = \frac{4(1-\phi_{\mathrm{vap}})L_{\mathrm{tot}}}{f};\ \text{and}\ \mathcal{P}_{\mathrm{int}} = \frac{fA_{\mathrm{tot}}}{L_{\mathrm{tot}}\sqrt{\tau_{\mathrm{apo}}}} \quad \text{(S19)}$$

These lateral, microscale dimensions ($d_{\mathrm{apo}}$ and $d_{\mathrm{c-c}}$) only serve to evaluate our 1-D approximation of transport in the cell-to-cell and apoplasmic domains. We note that several of these typical apoplasmic and cell-to-cell paths occupy each repeat unit (e.g., $n_{\mathrm{apo}} = 4\phi A_{\mathrm{tot}}/\pi d_{\mathrm{apo}}^2\sqrt{\tau_{\mathrm{apo}}}$ apoplasmic paths). The perimeter, $\mathcal{P}_{\mathrm{int}}$ enters our calculations to scale the interfacial conductance between the cell-to-cell and the apoplasmic paths.

The permissible ranges for the parameters are as follows: $A_{\mathrm{tot}} = 100^2 - 120^2 = 10 - 14.4 \times 10^3$ (μm²) and typical ranges of porosity, $\phi_{\mathrm{vap}} = 0.2 - 0.5$, $\tau_{\mathrm{vap}}^{\mathrm{bs}} = 1 - 5$, $\tau_{\mathrm{vap}}^{\mathrm{mes}} = 1 - 4$, $d_{\mathrm{cw}} = 0.2$ (μm), and $f = 10 - 13$ (-) we have: $A_{\mathrm{apo}} = 1.4 - 6 \times 10^3$ (μm²), $A_{\mathrm{c-c}} = 3.6 - 9.4 \times 10^3$ (μm²), $d_{\mathrm{c-c}} = 31 - 64$ (μm), $\mathcal{P}_{\mathrm{int}} = 223 - 845$ (μm), and $A_{\mathrm{cw}} = 63 - 132$ (μm²). We note that $A_{\mathrm{cw}} \ll A_{\mathrm{vap}}$ such that $V_{\mathrm{apo}} \cong V_{\mathrm{vap}}$ and $A_{\mathrm{apo}} \cong A_{\mathrm{vap}}$, as assumed in Eqs. S14 and S18.

Table S4 provides details of the parameters for leaf anatomy used in this model.

**Coupled mass balance in cell-to-cell path and apoplasm:**
At steady state, we have the following molar mass balances for differential element, $\Delta z$ relating axial flows in the cell-to-cell path ($m_{\mathrm{z}}^{\mathrm{c-c}}$ [mol/s]), in the apoplasm ($m_{\mathrm{z}}^{\mathrm{apo}}$), and between these paths ($m_{\mathrm{s-a}}$) (Fig. S19c):

$$m_z^{\mathrm{c-c}}(z+\Delta z) - m_z^{\mathrm{c-c}}(z) = -m_{\mathrm{s-a}} \quad \text{(S20)}$$

$$m_z^{\mathrm{apo}}(z+\Delta z) - m_z^{\mathrm{apo}}(z) = m_{\mathrm{s-a}} \quad \text{(S21)}$$

All symbols are defined in Tables S2-S5.

**Constitutive relation for axial flow in the cell-to-cell path:**
We assume $m_{\mathrm{z}}^{\mathrm{c-c}}$ is driven by gradients in water potential, $\psi_{\mathrm{c-c}}(z)$ [MPa]. Assuming that osmotic potential is approximately uniform through the domain, this gradient in water potential approximates that in cell turgor pressure, $P_{\mathrm{c-c}}(z)$ [MPa]:

$$m_z^{\mathrm{c-c}} = -k_{\mathrm{c-c}}A_{\mathrm{c-c}}\frac{d\psi_{\mathrm{c-c}}}{dz} \cong -k_{\mathrm{c-c}}A_{\mathrm{c-c}}\frac{dP_{\mathrm{c-c}}}{dz} \quad \text{(S22)}$$

Here, $k_{\mathrm{c-c}}$ [mol/m/s/MPa] is the effective axial hydraulic conductivity and $A_{\mathrm{c-c}}$ [m²] (Eq. S17) is the cross-sectional area of the cell-to-cell pathway for water transport. With the assumption of uniform osmotic potential, the process modelled in Eq. S22 may be interpreted as: (i) purely symplasmic flow between cells via plasmodesmata driven by the difference in turgor pressure between adjacent cells; (ii) as a transcellular flow passing through the plasma membrane of the upstream cell, through the cell walls of the adjacent cells, and through the plasma membrane of the downstream cell driven by the difference in water potential between these cells; or (iii) a combination of flows along these two paths between adjacent cells.[12,17,18,19] Our model does not distinguish between these scenarios. As discussed below, we constrain the value of $k_{\mathrm{c-c}}$ to ensure that the water potentials of the mesophyll cell remain above the turgor loss point in maize ($\psi_{\mathrm{c-c}} > \psi_{\mathrm{TLP}} \cong -2$ MPa).

**Constitutive relations for axial flows in the apoplasm:**
In the apoplasm, we account for flow along parallel, axial (along $z$) paths through the cell wall as a liquid and through the IAS as a vapor (expanded view in Fig. S19b). The following relations hold for both the bundle sheath and mesophyll zones in the apoplasm (Fig. S19c) with distinct values of the porosity, $\phi_{\mathrm{vap}}$ [-] and tortuosity, $\tau_{\mathrm{vap}}$ [-] (Table S4). Firstly, for the vapor component, we model the transport of vapor through the IAS as driven by gradients in mole fraction, $w(z)$ for the dilute vapor mixture in air:[20]

$$m_{\mathrm{z}}^{\mathrm{vap}} = -cD\frac{A_{\mathrm{vap}}}{\tau_{\mathrm{vap}}}\frac{dw}{dz} \tag{S23}$$

where $c$ [mol/m$^3$] is the molar concentration of air, $D$ [m$^2$/s] is the molecular diffusivity of water vapor in air, and $w$ is the mole fraction of vapor.

We assume local equilibrium between the vapor and the liquid water in its adjacent cell walls: $\psi_{\mathrm{vap}}(z) = \psi_{\mathrm{cw}}(z) = \psi_{\mathrm{apo}}(z)$.[21] This assumption allows us to combine the fluxes in the cell wall and vapor path into an apoplasmic flux driven by a single gradient ($\frac{d\psi_{\mathrm{apo}}}{dz}$). Going forward, we will use $\psi_{\mathrm{apo}}$ to represent the water potential in this effective medium view of the apoplasm.

We further linearize Kelvin's equation (Eq. S6) to express gradients in vapor concentrations in terms of water potential:

$$\frac{dw_{\mathrm{vap}}}{dz} = \frac{dw}{d\psi_{\mathrm{vap}}}\frac{d\psi_{\mathrm{vap}}}{dz} \cong \left(\frac{\bar{v}p_{\mathrm{sat}}}{RTP_{\mathrm{atm}}}\right)\frac{d\psi_{\mathrm{vap}}}{dz} = \left(\frac{\bar{v}\mathrm{c}}{P_{\mathrm{atm}}}\right)\frac{d\psi_{\mathrm{apo}}}{dz} \tag{S24}$$

where $p_{\mathrm{sat}} = cRT$ [Pa] is the saturation vapor pressure at the leaf temperature, $T$ [K], $R$ [J/K/mol] is the ideal gas constant, $\bar{v}$ [mol/m$^3$] is the molar volume of liquid water, and $P_{\mathrm{atm}}$ is the air pressure. The linearization in Eq. (S24) is accurate for $\psi_{\mathrm{vap}}\bar{v} \ll RT_{\mathrm{leaf}}$, a condition that holds even for the largest undersaturation studied here.

With Eq. S24, we can express Eq. S23 as:

$$m_{\mathrm{z}}^{\mathrm{vap}} = -k'_{\mathrm{vap}}\frac{A_{\mathrm{vap}}}{\tau_{\mathrm{vap}}}\frac{d\psi_{\mathrm{apo}}}{dz} \tag{S25}$$

where $k_{\mathrm{vap}} = k'_{\mathrm{vap}}/\tau_{\mathrm{vap}} = cD\left(\frac{\bar{v}p_{\mathrm{sat}}}{\tau_{\mathrm{vap}}RTP_{\mathrm{atm}}}\right)$ [mol/m/s/MPa] is the vapor conductivity.

The flux of water in the porous cell wall is defined with Darcy's law:

$$m_z^{cw} = -k'_{cw}\frac{A_{cw}}{\tau_{vap}}\frac{d\psi_{cw}}{dz} = -k_{cw}A_{cw}\frac{d\psi_{apo}}{dz} \tag{S26}$$

where we use a value for the conductivity of the cell wall, $k'_{cw}$ [mol/m/s/MPa] adopted from Michael et al.[22] and used the same tortuosity as for the vapor path.

We combine Eqs. (S25) and (S26) to obtain the total apoplasmic flux as a sum of the flux through the cell wall and the vapor flux:

$$m_z^{apo} = m_z^{vap} + m_z^{cw} = -\left(A_{cw}k_{cw} + A_{vap}k_{vap}\right)\frac{d\psi_{apo}}{dz} \tag{S27}$$

**Water transport across plasma membrane:**
The transmembrane movement of water molecules is driven by a local difference in water potential between the cell-to-cell path and the cell wall (which we take to be in local equilibrium with the IAS):

$$m_{s-a} = \kappa_{mem}\left(\psi_{c-c} - \psi_{apo}\right)\Delta z \tag{S28}$$

where $\kappa_{mem}$ [mol/m/s/MPa] is the hydraulic conductivity of the plasma membrane. We define a range of values for $\kappa_{mem}$ based on osmotic permeabilities ($P_{os}$ [$\mu$m/s]) measured in protoplast-swelling assays.[23-25] The relationship between $\kappa_{mem}$ and $P_{os}$ is:

$$\kappa_{mem} = \frac{P_{os}d_{c-c}}{2RT}$$

We can now assess the appropriateness of our pseudo 1-D model by assessing the ratio of the lateral resistance within each domain to that of the interface between them. This ratio is called the Biot number, $\mathcal{B}$ [-]; we expect the pseudo 1-D approximation to be accurate for $\mathcal{B} \ll 1$.[13] For the cell-to-cell and vapor paths (Eq. S19), we have:

$$\mathcal{B}_{c-c} = \frac{\kappa_{mem}}{k_{c-c}}\,;\, \mathcal{B}_{vap} = \frac{\kappa_{mem}}{k_{vap}} \tag{S29}$$

For typical values of the parameters (see Table S4), we find $\mathcal{B}_{c-c} \sim 0.002$ and $\mathcal{B}_{vap} \sim 0.2$. These values indicate that our pseudo 1-D approximation is fully appropriate for the cell-to-cell path ($\mathcal{B}_{c-c} \ll 1$) and reasonably appropriate for the apoplasmic path ($\mathcal{B}_{vap} < 1$), although we expect that our treatment does fail to capture the effects of some weak lateral gradients in this path.

**Coupled governing equations:**
We substitute the expressions for the flows (Eqs. S22, S27, and S28) into the mass balances (Eqs. S20 and S21), divide through by $\Delta z$, and allow $\Delta z$ to go to zero to obtain the governing differential equations for the water potential distribution through the thickness of the leaf (Fig. 19c):

$$\frac{d^2\psi_{c-c}}{dz^2} = \frac{\kappa_{mem}}{k_{c-c}A_{c-c}}\left(\psi_{c-c} - \psi_{apo}\right) \tag{S30}$$

$$\frac{d^2\psi_{\text{apo}}}{dz^2} = -\frac{\kappa_{\text{mem}}}{A_{\text{cw}}k_{\text{cw}} + A_{\text{vap}}k_{\text{vap}}}(\psi_{\text{c-c}} - \psi_{\text{apo}}) \tag{S31}$$

These governing equations are similar in structure to the equations used to model heat transfer from extended surfaces such as fins and yield analytical solutions for $\psi_{\text{c-c}}(z)$ and $\psi_{\text{apo}}(z)$ (see below).[13]

**Boundary and interfacial conditions:**
We solve Eqs. S30 and S31 simultaneously in each of four domains (Fig. 19c): the adaxial and abaxial bundle sheath zones ($0 < z < L_{\text{bs}}/2$ and $-L_{\text{bs}}/2 < z < 0$) and the abaxial and abaxial mesophyll zones ($L_{\text{bs}}/2 < z < L_{\text{tot}}/2$ and $-L_{\text{tot}}/2 < z < -L_{\text{bs}}/2$). This abaxial-adaxial division allows us to apply boundary conditions independently on the two surfaces of a leaf, thereby, extending this formulation to model the OXZ in plants with hypostomatous leaves or obstruction on one side (as in Fig. 4h of main text). The division into bundle sheath and mesophyll zones allow us to define distinct geometric and transport parameters for these regions, a useful feature to account for variation in porosity and tortuosity with depth and to confront architectures that propose different hydraulic properties of bundle sheath membranes relative to mesophyll membranes.[26]

**Cell-to-cell path:** Water enters the bundle sheath zones of the cell-to-cell paths on both the abaxial and adaxial side at $z = 0$. We neglect any drop in potential as water moves from the xylem into these zones:

$$\psi_{\text{c-c}}^{\text{bs}}(z = 0^{\pm}) = \psi_{\text{xyl}} \tag{S32}$$

At the outer boundaries of the mesophyll zones of the cell-to-cell paths ($z = \pm L_{\text{tot}}/2$), we impose a no-flux boundary condition based on the assumption that cuticular transpiration is negligible:

$$\left.\frac{d\psi_{\text{c-c}}^{\text{mes}}}{dz}\right|_{z=\pm L_{\text{tot}}/2} = 0 \tag{S33}$$

At the interfaces between bundle sheath and mesophyll zones ($z = \pm L_{\text{bs}}/2$), we impose continuity of the water potentials:

$$\psi_{\text{c-c}}^{\text{bs}}(z = \pm L_{\text{bs}}/2) = \psi_{\text{c-c}}^{\text{mes}}(z = \pm L_{\text{bs}}/2) \tag{S34}$$

**Apoplasm**: At the stomates, we apply flux boundary conditions at both the abaxial and adaxial ends ($z = \pm\frac{L_{\text{tot}}}{2}$). The stomates are represented in this model as an interface of infinitesimal thickness such that its conductance to water vapor can be described by a discrete parameter $g_s$ (stomatal conductance). Here, we neglect any boundary layer resistance (typically, $< 0.05 \times (1/g_s)$).

$$\mp\left(A_{\text{cw}}^{\text{mes}}k_{\text{cw}} + A_{\text{vap}}^{\text{mes}}k_{\text{vap}}\right)\left.\frac{d\psi_{\text{apo}}^{\text{mes}}}{dz}\right|_{z=\pm\frac{L_{\text{tot}}}{2}} = A_{\text{tot}}g_s\left(w_{\text{apo}}\left(z = \pm\frac{L_{\text{tot}}}{2}\right) - w_{\text{a}}\right) \tag{S35}$$

At the mid-plane ($z = \pm L_{\text{bs}}/2$), we impose continuity of water potentials and fluxes:

$$\psi_{\text{apo}}^{\text{bs}}(z=0^{+}) = \psi_{\text{apo}}^{\text{bs}}(z=0^{-}) \tag{S36}$$

$$\left.\frac{d\psi_{\text{apo}}^{\text{bs}}}{dz}\right|_{z=0^{+}} = \left.\frac{d\psi_{\text{apo}}^{\text{bs}}}{dz}\right|_{z=0^{-}} \tag{S37}$$

At the interfaces between bundle sheath and mesophyll zones ($z = \pm L_{\text{bs}}/2$), we impose continuity of the water potentials:

$$\psi_{\text{apo}}^{\text{bs}}(z = \pm L_{\text{bs}}/2) = \psi_{\text{apo}}^{\text{mes}}(z = \pm L_{\text{bs}}/2) \tag{S38}$$

**Transpiration rate and conductance of OXZ:**
We calculate the transpiration rates at the two surfaces with the flows in Eq. S35:

$$E^{\text{ab/ad}} = A_{\text{tot}} g_s \left( w_{\text{apo}}\left(z = \pm \frac{L_{\text{tot}}}{2}\right) - w_{\text{a}} \right) = A_{\text{tot}} g_s \left( w_{\text{ssc}}^{\text{ab/ad}} - w_{\text{a}} \right) \tag{S39}$$

where $w_{\text{ssc}}^{\text{ab}}$ and $w_{\text{ssc}}^{\text{ad}}$ are the vapor fractions in the substomatal cavity that we associate with the values of water status measured by AquaDust (Fig. 2b in main text).

Here, we re-express the conductance of the OXZ (Eq. S1) within the model:

$$g_{\text{OXZ}} = \frac{E^{\text{ad}} + E^{\text{ab}}}{w_{\text{xyl}} - w_{\text{apo}}\left(z = \frac{L_{\text{tot}}}{2}\right)} = \frac{E^{\text{ad}} + E^{\text{ab}}}{w_{\text{xyl}} - w_{\text{ssc}}^{\text{ad}}} \tag{S40}$$

**Solution technique**
The coupled governing Eqs. (S30) and (S31), can be solved analytically. We assume trial solutions of the form:

$$\psi_{\text{c-c}} = \alpha e^{rz} \tag{S41}$$

$$\psi_{\text{apo}} = \beta e^{rz} \tag{S42}$$

Substituting these trial solutions in Eqs. (S30) and (S31) gives four roots of the auxiliary equations:

$$\{r_1, r_2, r_3, r_4\} = \left\{ 0, 0, \pm \sqrt{\kappa_{\text{mem}} \left( \frac{1}{k_{\text{c-c}} A_{\text{c-c}}} + \frac{1}{A_{\text{cw}} k_{\text{cw}} + A_{\text{vap}} k_{\text{vap}}} \right)} \right\} \tag{S43}$$

Substituting the roots back into each auxiliary equation provides a relationship between the coefficients $\alpha$ and $\beta$ appearing in Eqs. (S41) and (S42) with the following expressions describing the unknown potential fields:

$$\psi_{\text{c-c}} = \alpha_1 + \alpha_2 z + \alpha_3 e^{r_3 z} + \alpha_4 e^{-r_3 z} \tag{S44}$$

$$\psi_{\text{apo}} = \alpha_1 + \alpha_2 z - \frac{k_{\text{c-c}} A_{\text{c-c}}}{A_{\text{cw}} k_{\text{cw}} + A_{\text{vap}} k_{\text{vap}}} \alpha_3 e^{r_3 z} - \frac{k_{\text{c-c}} A_{\text{c-c}}}{A_{\text{cw}} k_{\text{cw}} + A_{\text{vap}} k_{\text{vap}}} \alpha_4 e^{-r_3 z} \tag{S45}$$

There are four unknown parameters ($\alpha_1, \alpha_2, \alpha_3, \alpha_4$) for each of the four domains (bundle sheath and mesophyll zones of both adaxial and abaxial sides) and they are obtained using the sixteen boundary conditions stated earlier (Eqs. S32-S38). The equations are solved in MATLAB using the nonlinear system of equations solver - fsolve.

**Predictions for four hypothetical scenarios**

We have made predictions for four hypothetical scenarios about hydraulic properties of the OXZ. In each case below, the hydraulic conductivity of one component of the OXZ (symplasm or, apoplasm or, plasma membrane) changes as a function of $\psi_{\mathrm{xyl}}$. For the component of the OXZ with variable hydraulic conductivity, a functional form of that hydraulic conductivity is provided as an input to the model. For components whose hydraulic conductivity does not change, a constant value is provided as an input to the simulation.

1. **Preferred hypothesis:** Limiting and variable resistance presented by plasma membrane at the interface between the cell-to-cell and apoplasmic paths (Fig. 4 in main text).
2. **Alternative hypothesis 1:** Limiting and variable resistance presented by cell-to-cell path with local equilibrium between liquid and vapor paths (Fig. S20).[11,12]
3. **Alternative hypothesis 2:** Limiting and variable resistance presented by axial flow through cell walls (Fig. S21)[12]
4. **Alternative hypothesis 3:** Limiting and variable resistance presented by the plasma membranes in the bundle sheath (Fig. S22).[26]

In each case, we find solutions of the forms in Eqs. S30 and S31 for the profiles of water potential, $\psi_{\mathrm{c-c}}(z)$ and $\psi_{\mathrm{apo}}(z)$ across the bundle sheath and mesophyll domains. We use these profiles to evaluate $g_{\mathrm{OXZ}}$ (Eq. S40) and saturation in the abaxial sub-stomatal cavity ($w_{\mathrm{ssc}}^{\mathrm{ab}} = w_{\mathrm{apo}}\left(z = \frac{L_{\mathrm{tot}}}{2}\right)$ from $\psi_{\mathrm{apo}}(z = \frac{L_{\mathrm{tot}}}{2})$ with the Kelvin equation, Eq. S6). We judge the compatibility of each scenario with experiments based on the following criteria:

1. **Criterion 1:** Ability to match variation in $g_{\mathrm{OXZ}}$ with $\psi_{\mathrm{xyl}}$ (panels d in figures).
2. **Criterion 2:** Ability to match variation in internal saturation, $h_{\mathrm{ssc}}^{\mathrm{ad}}$ with $\psi_{\mathrm{xyl}}$ (panels e in figures).
3. **Criterion 3:** Ability to predict disequilibrium between undersaturated apoplasm and turgid cells as documented in Fig. 3 of the main text.

**Preferred hypothesis: Large cell-to-cell conductance and both mesophyll and bundle sheath plasma membranes undergo loss of conductivity to water under drought stress (Fig. 4 in main text)**

Based on the observed nonlinear dependence of $g_{\mathrm{OXZ}}$ on $\psi_{\mathrm{xyl}}$, we define a response function for the hydraulic conductivity of both mesophyll and bundle sheath plasma membranes, $\kappa_{\mathrm{mem}}$, that is a sigmoidal function of the upstream xylem potential:

$$\kappa_{\mathrm{mem}} = \frac{\kappa_{\mathrm{mem}}^{\mathrm{max}}}{1+e^{-(a_1\psi_{\mathrm{xyl}}+\psi_{50})}} + \kappa_{\mathrm{mem}}^{\mathrm{min}} \qquad \text{(S46)}$$

Here, $\kappa_{\text{mem}}^{\text{max}}$ [mol/m/s/MPa] is the maximum conductivity of the plasma membrane, $\kappa_{\text{mem}}^{\text{min}}$ [mol/m/s/MPa] is the minimum conductivity of the plasma membrane, $a_1$ [MPa$^{-1}$] is a measure of the sensitivity of $\kappa_{\text{mem}}$ to $\psi_{\text{xyl}}$, and $\psi_{50}$ [MPa] is the characteristic xylem potential at which the plasma membrane undergoes loss of conductance to water.

To adjust the parameters in Eq. S46 and the governing Eqs. S30-31, we proceeded as follows:

1. **Geometry:** We constrained $A_{\text{c-c}}$ to $5 - 7 \times 10^3$ (μm$^2$), $A_{\text{vap}}$ to $5 - 6 \times 10^3$ (μm$^2$), $\tau_{\text{bs}}$ to $1 - 4$, and $\tau_{\text{mes}}$ to $1 - 4$. The values of $A_{\text{c-c}}$ and $A_{\text{vap}}$ used to generate predictions fall within the permissible range based on our confocal image data (see SI Fig. S19a) and the values of $\tau_{\text{bs}}$ and $\tau_{\text{mes}}$ correspond to a subset of those reported in the literature.[14,15] The values of geometric parameters were used across all four scenarios.
2. **Cell-to-cell and cell-wall conductances:** We held the hydraulic conductivities of the cell-to-cell path, $k_{\text{c-c}}$, and cell wall, $k_{\text{cw}}$ constant in this scenario in which we attribute all variability in $g_{\text{OXZ}}$ to variability in $\kappa_{\text{mem}}$. We constrained the value of $k_{\text{c-c}}$ with the observation of maintenance of turgor ($\psi_{\text{c-c}} > \psi_{\text{TLP}} \approx -2$ MPa) even as $\psi_{\text{xyl}}$ falls to $\sim -1.4$ MPa. All symbols, their nominal values, and relevant references are stated in Table S5. This vulnerability function has been plotted in Fig. 4c.
3. **Plasma membrane conductance:** We adjusted the values $\kappa_{\text{mem}}^{\text{max}}$ and $\kappa_{\text{mem}}^{\text{min}}$ such that the predicted values of $g_{\text{OXZ}}$ (Eq. S40) at $\psi_{\text{xyl}} = -0.2$ MPa and $\psi_{\text{xyl}} = -1.4$ MPa closely matched the mean measured values in the corresponding bins (Fig. 4d) while ensuring that the predicted values of $h_{ssc}^{apo}$ remained within the standard error in $h_{ssc}^{apo}$ measurements (Fig. 4e). Next, we used linear interpolation between the measured values of $g_{\text{OXZ}}$ at $\psi_{\text{xyl}} = -0.2$ MPa and $\psi_{\text{xyl}} = -0.6$ MPa (Fig. 4e) to estimate a value for $\psi_{50}$. Finally, with $\kappa_{\text{mem}}^{\text{max}}$, $\kappa_{\text{mem}}^{\text{min}}$, and $\psi_{50}$ held fixed, we varied $a_1$ and chose the value that minimized the least-squares difference between experimental and model-based $g_{\text{OXZ}}$. The resulting values of the parameters in Eq. S46) were: $\kappa_{\text{mem}}^{\text{max}} = 1.1 \pm 0.1 \times 10^{-5}$ (mol/m/s/MPa), $\kappa_{\text{mem}}^{\text{min}} = 4.1 \times 10^{-8} \pm 0.4 \times 10^{-8}$ (mol/m/s/MPa), $a_1 = 1.1 \times 10^{-5} \pm 0.2 \times 10^{-5}$ (MPa$^{-1}$), and $\psi_{50} = -0.22 \pm 0.02$ (MPa). We generate uncertainty intervals on the predictions by propagating the extreme values of the range of parameter values through the model (see Tables S4 and S5 for these values). Note that the values of $\kappa_{\text{mem}}$ are within the bounds obtained from maize leaf protoplast-swelling experiments[25] (Fig. 4c) and this range in maize lies with the range defined by measurements from protoplasts across multiple species.[23,24]

**Criterion 1:** Yes. Here, predicted $g_{\text{OXZ}}$ falls by a factor $\sim 25$ in the model prediction in agreement with our data (Fig. 4e).

**Criterion 2:** Yes. Apoplasmic potentials dropped from near saturation $h_{ssc}^{apo} \sim 1$ at well-watered conditions ($\psi_{\text{xyl}} = -0.2 \pm 0.2$ MPa) to $h_{ssc}^{apo} \sim 0.92$ under moderate drought ($\psi_{\text{xyl}} = -1.4 \pm 0.2$ MPa), matching our measurements (Fig. 4e).

**Criterion 3:** Yes. This model predicts significant undersaturation in the mesophyll apoplasm ($h_{ssc}^{apo} \sim 0.92$ ) while mesophyll cells remains at potentials above the turgor loss point of maize leaves, enabled by high cell-to-cell conductivity (Fig. 4g). Furthermore, the predicted profile in the IAS agrees with those measured via confocal (Fig. 4g).

**Conclusion**: We conclude that this hypothesis is compatible, qualitatively and quantitatively with our observations.

**Alternative hypothesis 1: Local equilibrium between cell-to-cell path and apoplasm and cell-to-cell path becomes less permeable to water under drought stress**

Here, the cell-to-cell path is assumed to be in local equilibrium with the apoplasm and the cell-to-cell path is the dominant, variable conductance path for water transport in the leaf. To maintain near equilibrium between the cell-to-cell path and apoplasm we take $\kappa_{\mathrm{mem}} = 7 \times 10^{-6}$ (mol/m/s/MPa). This value is near the highest conductivity measured for a plasma membrane from protoplast-swelling assays. This scenario forces the model presented here to mimic the conditions assumed within the effective medium, local equilibrium hydraulic model of Rockwell et al. 2014.[11] We have not come across a reference that explicitly proposes that the cell-to-cell path becomes less permeable to water with drought stress. However, there are reports of reduction in permeability of plasmodesmata in response to increased intracellular Calcium levels during drought,[27] so we proceed to implement variable conductance in this case by lowering the value of the hydraulic conductivity of the cell-to-cell path, $k_{\mathrm{c-c}}$ in our model. It should be noted that as the value of $k_{\mathrm{c-c}}$ decreases, the relative contribution of the apoplasm in OXZ water transport increases. Again, based on the observed nonlinear dependence of $g_{\mathrm{OXZ}}$ on $\psi_{\mathrm{xyl}}$, we define $k_{\mathrm{c-c}}$ to be a sigmoidal function of the upstream xylem potential:

$$k_{\mathrm{c-c}} = \frac{k_{\mathrm{c-c}}^{\max}}{1+e^{-(a_1\psi_{\mathrm{xyl}}+\psi_{50})}} + k_{\mathrm{c-c}}^{\min} \qquad \text{(S47)}$$

Here, $k_{\mathrm{c-c}}^{\max} = 1.1 \times 10^{-5} \pm 0.1 \times 10^{-5}$ (mol/m/s/MPa) is the maximum conductivity of the cell-to-cell path, $k_{\mathrm{c-c}}^{\min} = 7 \times 10^{-8} \pm 0.5 \times 10^{-8}$ (mol/m/s/MPa) is the minimum conductivity of the cell-to-cell path, $a_1 = 1 \times 10^{-5} \pm 0.1 \times 10^{-5}$ ($\mathrm{MPa}^{-1}$) is a measure of the sensitivity of $k_{\mathrm{c-c}}$ to $\psi_{xylem}$, and $\psi_{50} = -0.23 \pm 0.01$ (MPa) is the characteristic xylem potential at which the cell-to-cell path undergoes loss of conductance to water. The limits of $k_{\mathrm{c-c}}$ are chosen to give ourselves the best chance of capturing the range of $g_{\mathrm{OXZ}}$ observed in experiments while staying within physiological bounds set by protoplast-swelling experiments. Fine-tuning of hydraulic parameters was not performed since this architecture failed to agree with experiments qualitatively as shown below. This vulnerability function has been plotted in Fig. S20c. The values of geometric parameters are mentioned in table S4 'Alternative hypothesis 1'.

**Criterion 1**: No. Here, predicted $g_{\mathrm{OXZ}}$ falls by a factor $\sim 5$ in the model prediction. This is not in agreement with our data (Fig. S20d).

**Criterion 2:** No. the change in apoplasmic potentials do not match experimental data (Fig. S20e).

**Criterion 3:** No. The low conductance of the cell-to-cell pathway under drought-stressed conditions pulls the symplasmic potentials down to the apoplasm potentials (i.e., local equilibrium) which are below $\psi_{\mathrm{TLP}}$ through most of the thickness in the water-limited case (Figs. S20f-g).

**Conclusion**: This hydraulic architecture is not compatible qualitatively (turgor loss) or quantitatively (minimal undersaturation) with our observations.

**Alternative hypothesis 2: Cell wall is the dominant hydraulic pathway, and it becomes less permeable to water under drought stress**

Here, we assume that the cell wall within the apoplasm is the dominant, variable conductance path for water transport in the leaf, as suggested by Buckley.[12] Flow of water occurs predominantly in the apoplasm through the full thickness of the leaf ($k_{\mathrm{c-c}} < k_{\mathrm{apo}}$). Based on the observed nonlinear dependence of $g_{\mathrm{oxz}}$ on $\psi_{xylem}$, we define $k_{cw}$ to be a sigmoidal function of the upstream xylem potential:

$$k_{\mathrm{cw}} = \frac{k_{\mathrm{cw}}^{\max}}{1+e^{-(a_1\psi_{\mathrm{xyl}}+\psi_{50})}} + k_{\mathrm{cw}}^{\min} \quad \text{(S48)}$$

Here, $k_{\mathrm{cw}}^{\max} = 2.7 \times 10^{-6} \pm 0.2 \times 10^{-6}$ mol/m/s/MPa is the maximum conductivity of the cell wall, $k_{\mathrm{vap}}$ serves as the minimum conductivity of the cell wall when the pores are dry, $a_1 = 10^{-5} \pm 0.2 \times 10^{-5}$ MPa$^{-1}$ is a measure of the sensitivity of $k_{\mathrm{cw}}$ to $\psi_{\mathrm{xyl}}$, and $\psi_{50} = -0.25 \pm 0.02$ MPa is the characteristic xylem potential at which the cell wall undergoes loss of conductance to water. Fine-tuning of hydraulic parameters was not performed since this architecture failed to agree with experiments qualitatively as shown below. This vulnerability function has been plotted in Fig. S21c. The values of geometric parameters are mentioned in table S4 'Alternative hypothesis 2'.

**Criterion 1:** No. Here, the predicted variation in $g_{oxz}$ does not match the observed factor $\sim 25$ in experiments (Fig. S21d).

**Criterion 2:** No. Again, the change in apoplasmic potentials do not match experimental data (Fig. S21e).

**Criterion 3:** No. Since the cell wall is designed to carry most of the transpiration flux in this hydraulic architecture, the low conductance of the cell-to-cell pathway pulls the symplasmic potentials down to the apoplasm potentials. In other words, this architecture predicts local equilibrium between symplasm and apoplasm similar to alternative hypothesis 1 above (Fig. S21g).

**Conclusion**: This hydraulic architecture is not compatible qualitatively (turgor loss) or quantitatively (minimal undersaturation) with our observations. Conductance (= conductivity per unit length x cross-sectional flow area) of the cell wall pathway is not large enough to carry a flux that is commensurate with the transpiration rate. Buckley[12] proposed a hydraulic model where cell-to-cell transport is transmembrane, and their estimate of plasma membrane conductance led them to conclude that apoplasmic transport dominates in the leaf. Excluding the possibility of cell-to-cell transport via plasmodesmata may have been the reason why the symplasm conductance was underestimated in their model.

**Alternative hypothesis 3: Bundle-sheath membrane is the dominant hydraulic resistance in the outside-xylem tissue**

Here, the cell-to-cell path is assumed to be the dominant path for water transport in the leaf with the bundle sheath membranes having a variable conductance, as suggested by Shatil-Cohen et al.[26] A feature of this hypothesis is that the mesophyll membrane is not the dominant hydraulic resistance and maintains a relatively high value. Based on the observed nonlinear dependence of $g_{\mathrm{OXZ}}$ on $\psi_{\mathrm{xyl}}$, we define a response function for the hydraulic conductivity of bundle sheath plasma membranes, $\kappa_{\mathrm{mem}}^{\mathrm{bs}}$, that is a sigmoidal function of the upstream xylem potential:

$$\kappa_{\mathrm{mem}}^{\mathrm{bs}} = \frac{\kappa_{\mathrm{mem}}^{\max}}{1+e^{-(a_1\psi_{\mathrm{xyl}}+\psi_{50})}} + \kappa_{\mathrm{mem}}^{\min} \quad \text{(S49)}$$

Here, $\kappa_{\mathrm{mem}}^{\max} = 9 \times 10^{-6} \pm 10^{-6}$ is the maximum conductivity of the plasma membrane, $\kappa_{\mathrm{mem}}^{\min} = 2 \times 10^{-7} \pm 1 \times 10^{-7}$ is the minimum conductivity of the plasma membrane, $a_1 = 1.1 \times 10^{-5} \pm 0.1 \times 10^{-5}$ MPa$^{-1}$ is a measure of the sensitivity of $\kappa_{\mathrm{mem}}$ to $\psi_{\mathrm{xyl}}$, and $\psi_{50} = -0.23 \pm 0.01$ MPa is the characteristic xylem potential at which the plasma membrane undergoes loss of conductance to water. The limits of $\kappa_{\mathrm{mem}}^{\mathrm{bs}}$ are chosen to give ourselves the best chance of capturing the range of $g_{\mathrm{OXZ}}$ observed in experiments while staying within physiological bounds set by protoplast-swelling experiments.  Fine-tuning of hydraulic parameters was not performed since this architecture failed to agree with experiments qualitatively as shown below. This vulnerability function has been plotted in S.I. Fig. S22c. The values of geometric parameters are mentioned in table S4 'Alternative hypothesis 3'.

**Criterion 1:** Yes. Here, predicted $g_{oxz}$ falls by a factor $\sim 25$ in the model prediction in agreement with our data (Fig. S22d).

**Criterion 2:** Yes. Apoplasmic potentials plummeted from near saturation $h_{\mathrm{ssc}}^{\mathrm{apo}} \sim 1$ at well-watered conditions ($\psi_{xylem} = -0.2 \pm 0.2$ MPa) to $h_{\mathrm{ssc}}^{\mathrm{apo}}$~0.94 under moderate drought ($\psi_{xylem} = -1.4 \pm 0.2$ MPa). Again, this prediction matches our data (Fig. S22e).

**Criterion 3:** No. The low conductance of the bundle sheath membranes between the bundle sheath and mesophyll cells pulls the mesophyll symplasmic potentials below the turgor loss point in maize (Fig. S22f-g). Predicted loss of turgor in mesophyll cells makes this hydraulic architecture inconsistent with our data.

**Conclusion**: While this model can predict the observed drop $g_{\mathrm{OXZ}}$ and $h_{\mathrm{ssc}}^{\mathrm{apo}}$, its prediction of symplastic potentials beneath the TLP make it incompatible with our observations. This architecture may be true for water potentials close to saturation.

**Neglect of temperature-gradient-driven vapor transport:**
In our model, we ignore non-isothermal effects on the variation of water potential through the leaf thickness. To justify this simplification, we perform an energy balance in the leaf to obtain the characteristic temperature difference across the leaf in our controlled-environment cuvette experiments:

$$\frac{0.76k_{\mathrm{c-c}}^{T}}{L_{\mathrm{c}}}\Delta T_{\mathrm{c}}=\frac{N_{A}hc}{\lambda_{\mathrm{eff}}}\times PAR \tag{S50}$$

Here, 0.76 is the liquid volume fraction; $k_{\mathrm{c-c}}^{\mathrm{T}}\approx 0.62$ W/m/K is the thermal conductivity of liquid water at 36 °C; $L_{\mathrm{c}}=L_{\mathrm{tot}}$ is the characteristic length scale over which light absorption occurs in an amphistomatous maize leaf with near symmetry across the leaf mid-plane;[28] $\Delta T_{\mathrm{c}}$ is the characteristic temperature difference across the leaf; $N_A$ is Avogadro's number; $h$ is Planck's constant; $c$ [m/s] is the speed of light in vacuum; $\lambda_{\mathrm{eff}}$ = 550 nm is the effective wavelength of white light imposed on the leaf inside a PPSystems CIRAS-3 cuvette; $PAR=1000$ $\mu$mol/m$^2$/s is the characteristic intensity of white light imposed on the leaf in our experiments. Solving, we obtain $\Delta T_c\sim 0.1$ K. Gradients in temperature give rise to gradients in vapor pressure, and, in turn, influence the effective conductance of the vapor path as follows:[11]

$$k_{\mathrm{vap}}^{*}\cong k_{\mathrm{vap}}\left(1+\frac{0.8\ \mathrm{MPa}}{\Delta\psi_{\mathrm{apo}}}\right) \tag{S60}$$

where, $\Delta\psi_{\mathrm{apo}}$ is the isothermal difference in water potential from the mid-plane to the stomates. This effect on vapor transfer can be non-negligible under well-watered conditions for which $\frac{0.8\ \mathrm{MPa}}{\Delta\psi_{\mathrm{apo}}}\sim 1$ (e.g., Fig. 4f) but should be negligible for cases in which $g_{\mathrm{OXZ}}$ has fallen and we observe and predict large differences in potential ($\Delta\psi_{\mathrm{apo}}\sim 10$ MPa – e.g., Figs. 4g-4h). In summary, we do not expect that non-isothermal effects impact the principal conclusions of this study with its focus on cases in which large gradients occur across the OXZ. Nonetheless, we intend to extend this model to account for non-isothermal effects in a future study.

**Table S2: Transport variables**

| Symbol | Definition | Nominal value/Range | Units |
|---|---|---|---|
| $z$ | Axial position on the thickness dimension of the leaf | $-L_{ab}$ to $L_{ad}$ | $\mu$m |
| $\psi$ | Water potential | | MPa |
| $\psi_{xyl}$ | Xylem water potential | $-0.2\pm 0.2$ to $-1.8\pm 0.2$ | MPa |
| $\psi_{c-c}$ | Water potential in cell-to-cell path | $O(-1)$ | MPa |
| $\psi_{apo}$ | Apoplasm water potential | $O(-10)$ | MPa |
| $\psi_{cw}$ | Cell wall water potential | $O(-10)$ | MPa |
| $\psi_{TLP}$ | Turgor loss point in maize | $-2$ | MPa |
| $m_z$ | Axial mass flow rate along z | | Mol/s |
| $tm$ | Transmembrane | | Mol/s |

| | | | |
|---|---|---|---|
| $sym$ | Symplasm | | |
| $apo$ | Apoplasm | | |
| $c-c$ | Cell-to-cell path | | |
| $cw$ | Cell wall | | |
| $vap$ | Vapor | | |
| $air$ | Atmosphere | | |
| $bs$ | Bundle sheath | | |
| $mes$ | Mesophyll | | |
| $k$ | Hydraulic conductivity per leaf area | | mol/m/s/MPa |
| $\kappa$ | Hydraulic conductivity per membrane surface area | | mol/m/s/MPa |
| $w$ | Mole fraction of water vapor | | – |

**Table S3: Thermodynamic parameters**

| Symbol | Formula | Definition | Nominal value/Range | Units | References |
|---|---|---|---|---|---|
| $T$ | | Leaf temperature | 309 | K | – |
| $R$ | | Universal gas constant | 8.314 | Pa.m$^3$/mol/K | – |
| $w_{\text{sat}}$ | | Saturation pressure at leaf temperature | 5946.2 | Pa | – |
| $\bar{v}$ | | Molar volume of water | $18 \times 10^{-6}$ | m$^3$/mol | – |
| $P_{\text{atm}}$ | | Atmospheric pressure | 101325 | Pa | – |
| $rh$ | | Relative humidity | 0.36 or 0.29 | – | – |
| $VPD$ | $p_{sat} \times \frac{(1-rh)}{10^3}$ | Vapor pressure deficit | 3.8 or 4.2 | kPa | Mean VPD in Fig. 2 is 3.8 kPa; 4.2 kPa for Figs. 4g and 4h |
| $c$ | | Molar concentration of air | 39.45 | mol/m$^3$ | – |
| $D$ | | Diffusivity of water vapor in air | $2.5 \times 10^{-5}$ | m$^2$/s | – |
| $\chi_0$ | $\frac{w_{sat}}{w_{atm}}$ | Reference mole fraction of water vapor at leaf temperature | 0.0587 | – | – |

**Table S4: Leaf anatomical parameters**

| Symbol | Formula | Definition | Nominal value/Permissible Range | Nominal value/range used in predictions | Units | References |
|---|---|---|---|---|---|---|
| $\tau$ | – | Tortuosity of air spaces | – | – | – | – |

| | | | | | | |
|---|---|---|---|---|---|---|
| Preferred hypothesis | | | | | | |
| $\tau_{\mathrm{bs}}$ | – | Tortuosity of air spaces adjacent to bundle sheath cells | $1-5$ | $1-4$ | – | [14,15] |
| $\tau_{\mathrm{mes}}$ | – | Tortuosity of air spaces adjacent to mesophyll cells | $1-4$ | $1-4$ | – | [14,15] |
| Alternative hypothesis 1 | | | | | | |
| $\tau_{\mathrm{bs}}$ | – | Tortuosity of air spaces adjacent to bundle sheath cells | $1-5$ | $1-3$ | – | [14,15] |
| $\tau_{\mathrm{mes}}$ | – | Tortuosity of air spaces adjacent to mesophyll cells | $1-4$ | $1-3$ | – | [14,15] |
| Alternative hypothesis 2 | | | | | | |
| $\tau_{\mathrm{bs}}$ | – | Tortuosity of air spaces adjacent to bundle sheath cells | $1-5$ | $1-3$ | – | [14,15] |
| $\tau_{\mathrm{mes}}$ | – | Tortuosity of air spaces adjacent to mesophyll cells | $1-4$ | $1-3$ | – | [14,15] |
| Alternative hypothesis 3 | | | | | | |
| $\tau_{\mathrm{bs}}$ | – | Tortuosity of air spaces adjacent to bundle sheath cells | $1-5$ | $1-4$ | – | [14,15] |
| $\tau_{\mathrm{mes}}$ | – | Tortuosity of air spaces adjacent to mesophyll cells | $1-4$ | $1-4$ | – | [14,15] |
| Parameters common across hypotheses | | | | | | |
| $\phi$ | – | Porosity of air spaces | – | – | – | – |
| $\phi_{\mathrm{bs}}$ | – | Porosity of air spaces adjacent to bundle sheath cells | $0.2-0.5$ | $0.46-0.5$ | – | [14,15] |
| $\phi_{\mathrm{mes}}$ | – | Porosity of air spaces adjacent to mesophyll cells | $0.2-0.5$ | $0.46-0.5$ | – | [14,15] |
| $f$ | – | Ratio of cell surface area exposed to apoplasm to leaf surface area | $10-13$ | $10-13$ | – | [16] |
| $L_{\mathrm{tot}}$ | – | Total thickness of maize leaf | 200 | 200 | $\mu$m | Fig. 2 in [3] |
| $L_{\mathrm{ad}}$ | – | Thickness of the adaxial section | 100 | 100 | $\mu$m | Fig. 2 in [3] |
| $L_{\mathrm{ab}}$ | – | Thickness of the abaxial section | 100 | 100 | $\mu$m | Fig. 2 in [3] |
| $L_{\mathrm{bs}}$ | – | Thickness of the bundle sheath in each section (adaxial and abaxial) of the leaf | 33.3 | 33.3 | $\mu$m | Fig. 2 in [3] |
| $d_{\mathrm{c-c}}$ | $\frac{4(1-\phi_{\mathrm{vap}})L_{\mathrm{tot}}}{f}$ | Diameter of cell-to-cell path | $31-64$ | $31-43$ | $\mu$m | – |
| $d_{\mathrm{cw}}$ | – | Thickness of cell wall | 200 | 200 | nm | [29] |
| $A_{\mathrm{c-c}}$ | $\frac{1-\phi_{\mathrm{vap}}}{\sqrt{\tau_{\mathrm{c-c}}}}A_{\mathrm{tot}}$ | Cross-sectional area for flow in cell-to-cell path | $3.6-9.4\times10^{-9}$ | $5-7\times10^{-9}$ | m$^2$ | – |
| $A_{\mathrm{vap}}$ | $\frac{\phi_{\mathrm{vap}}}{\sqrt{\tau_{\mathrm{apo}}}}A_{\mathrm{tot}}$ | Cross-sectional area for flow in air spaces | $1.4-6\times10^{-9}$ | $5-6\times10^{-9}$ | m$^2$ | – |
| $A_{\mathrm{cw}}$ | $d_{\mathrm{cw}}\mathcal{P}_{\mathrm{int}}$ | Cross-sectional area for flow in cell wall | $7.1-15.3\times10^{-11}$ | $8\times10^{-11}$ | m$^2$ | – |
| $A_{\mathrm{tot}}$ | $A_{\mathrm{c-c}}+A_{\mathrm{vap}}+A_{\mathrm{cw}}$ | Leaf cross-sectional area | $10-14.4\times10^{-9}$ | $10-13\times10^{-9}$ | m$^2$ | – |
| $\mathcal{P}_{\mathrm{int}}$ | $\frac{fA_{\mathrm{tot}}}{L_{\mathrm{tot}}\sqrt{\tau_{\mathrm{apo}}}}$ | Perimeter of cell-to-cell domain (and apoplasm domain due to continuity) | $223-845$ | 400 | $\mu$m | – |

### Table S5: Transport parameters

| Symbol | Formula | Definition | Nominal value/Permissible Range | Nominal value/range used in predictions | Units | References |
|---|---|---|---|---|---|---|
| Preferred hypothesis | | | | | | |
| $k_{\mathrm{cw}}^{\mathrm{bs}}$ | – | Hydraulic conductivity of bundle sheath cell wall | $\frac{2.7\times10^{-6}}{\tau_{\mathrm{bs}}}$ | $\frac{2.7\times10^{-6}}{\tau_{\mathrm{bs}}}$ | mol/m/s/MPa | [22] |
| $k_{\mathrm{cw}}^{\mathrm{mes}}$ | – | Hydraulic conductivity of mesophyll cell wall | $\frac{2.7\times10^{-6}}{\tau_{\mathrm{mes}}}$ | $\frac{2.7\times10^{-6}}{\tau_{\mathrm{mes}}}$ | mol/m/s/MPa | [22] |
| $k_{\mathrm{c-c}}$ | – | Hydraulic conductivity of cell-to-cell path | $5\times10^{-5}$ | $5\times10^{-5}$ | mol/m/s/MPa | Maintain $\psi_{\mathrm{c-c}}$ above TLP |
| $\kappa_{\mathrm{mem}}^{\mathrm{bs}}$ | – | Hydraulic conductivity of bundle sheath membranes per membrane surface area | $2.6\times10^{-8}$ to $1.3\times10^{-5}$ | $3.7\times10^{-8}$ to $1.2\times10^{-5}$ | mol/m/s/MPa | [25] |
| $\kappa_{\mathrm{mem}}^{\mathrm{mes}}$ | – | Hydraulic conductivity of mesophyll membranes per membrane surface area | $2.6\times10^{-8}$ to $1.3\times10^{-5}$ | $3.7\times10^{-8}$ to $1.2\times10^{-5}$ | mol/m/s/MPa | [25] |
| $a_1$ | – | Sensitivity of $\kappa$ to $\psi_{xylem}$ | – | $1.1\times10^{-5}\pm0.2\times10^{-5}$ | MPa$^{-1}$ | Fit to $g_{\mathrm{oxz}}$ from experiments |
| $\psi_{50}$ | – | Characteristic $\psi_{xylem}$ at which $\kappa$ changes | – | $-0.22\pm0.02$ | MPa | Fit to $g_{\mathrm{oxz}}$ from experiments |
| $\kappa_{\mathrm{mem}}^{\mathrm{max}}$ | – | Maximum hydraulic conductivity of plasma membrane | $1.3\times10^{-5}$ | $1.1\times10^{-5}\pm0.1\times10^{-5}$ | mol/m/s/MPa | [25] |
| $\kappa_{\mathrm{mem}}^{\mathrm{min}}$ | – | Minimum hydraulic conductivity of plasma membrane | $2.6\times10^{-8}$ | $4.1\times10^{-8}\pm0.4\times10^{-8}$ | mol/m/s/MPa | [25] |
| Alternative hypothesis 1 | | | | | | |
| $k_{\mathrm{cw}}^{\mathrm{bs}}$ | – | Hydraulic conductivity of bundle sheath cell wall | $\frac{2.7\times10^{-6}}{\tau_{\mathrm{bs}}}$ | $\frac{2.7\times10^{-6}}{\tau_{\mathrm{bs}}}$ | mol/m/s/MPa | [22] |
| $k_{\mathrm{cw}}^{\mathrm{mes}}$ | – | Hydraulic conductivity of mesophyll cell wall | $\frac{2.7\times10^{-6}}{\tau_{\mathrm{mes}}}$ | $\frac{2.7\times10^{-6}}{\tau_{\mathrm{mes}}}$ | mol/m/s/MPa | [22] |
| $\kappa_{\mathrm{mem}}^{\mathrm{bs}}$ | – | Hydraulic conductivity of bundle sheath membranes per membrane surface area | $2.6\times10^{-8}$ to $1.3\times10^{-5}$ | $7\times10^{-6}$ | mol/m/s/MPa | [25] |
| $\kappa_{\mathrm{mem}}^{\mathrm{mes}}$ | – | Hydraulic conductivity of mesophyll membranes per membrane surface area | $2.6\times10^{-8}$ to $1.3\times10^{-5}$ | $7\times10^{-6}$ | mol/m/s/MPa | [25] |
| $a_1$ | – | Sensitivity of $\kappa$ to $\psi_{xylem}$ | – | $1\times10^{-5}\pm0.1\times10^{-5}$ | MPa$^{-1}$ | Fit to $g_{\mathrm{oxz}}$ from experiments |
| $\psi_{50}$ | – | Characteristic $\psi_{xylem}$ at which $\kappa$ changes | – | $-0.23\pm0.01$ | MPa | Fit to $g_{\mathrm{oxz}}$ from experiments |
| $\kappa_{\mathrm{c-c}}^{\mathrm{max}}$ | – | Maximum hydraulic conductivity of cell-to-cell path | $1.3\times10^{-5}$ | $1.1\times10^{-5}\pm0.1\times10^{-5}$ | mol/m/s/MPa | [25] |

| | | | | | | |
|---|---|---|---|---|---|---|
| $\kappa_{\text{c-c}}^{\text{min}}$ | – | Minimum hydraulic conductivity of cell-to-cell path | $2.6 \times 10^{-8}$ | $7 \times 10^{-8} \pm 0.5 \times 10^{-8}$ | mol/m/s/MPa | [25] |
| | | | Alternative hypothesis 2 | | | |
| $\kappa_{\text{mem}}^{\text{bs}}$ | – | Hydraulic conductivity of bundle sheath membranes per membrane surface area | $2.6 \times 10^{-8}$ to $1.3 \times 10^{-5}$ | $10^{-5}$ | mol/m/s/MPa | [25] |
| $\kappa_{\text{mem}}^{\text{mes}}$ | – | Hydraulic conductivity of mesophyll membranes per membrane surface area | $2.6 \times 10^{-8}$ to $1.3 \times 10^{-5}$ | $10^{-5}$ | mol/m/s/MPa | [25] |
| $a_1$ | – | Sensitivity of $k$ to $\psi_{xylem}$ | – | $1 \times 10^{-5} \pm 0.2 \times 10^{-5}$ | $MPa^{-1}$ | Fit to $g_{\text{oxz}}$ from experiments |
| $\psi_{50}$ | – | Characteristic $\psi_{xylem}$ at which $k$ changes | – | $-0.25 \pm 0.02$ | MPa | Fit to $g_{\text{oxz}}$ from experiments |
| $k_{\text{cw}}^{\text{max}}$ | – | Maximum hydraulic conductivity of cell wall | $3 \times 10^{-6}$ | $2.7 \times 10^{-6} \pm 0.2 \times 10^{-6}$ | mol/m/s/MPa | [22] |
| | | | Alternative hypothesis 3 | | | |
| $k_{\text{cw}}^{\text{bs}}$ | – | Hydraulic conductivity of bundle sheath cell wall | $\frac{2.7 \times 10^{-6}}{\tau_{\text{bs}}}$ | $\frac{2.7 \times 10^{-6}}{\tau_{\text{bs}}}$ | mol/m/s/MPa | [22] |
| $k_{\text{cw}}^{\text{mes}}$ | – | Hydraulic conductivity of mesophyll cell wall | $\frac{2.7 \times 10^{-6}}{\tau_{\text{mes}}}$ | $\frac{2.7 \times 10^{-6}}{\tau_{\text{mes}}}$ | mol/m/s/MPa | [22] |
| $k_{\text{c-c}}$ | – | Hydraulic conductivity of cell-to-cell path | – | $\kappa_{\text{mem}}^{\text{bs}}$ | mol/m/s/MPa | Here, the limiting resistance in the cell-to-cell path is the bundle sheath resistance |
| $\kappa_{\text{mem}}^{\text{mes}}$ | – | Hydraulic conductivity of mesophyll membranes per membrane surface area | $2.6 \times 10^{-8}$ to $1.3 \times 10^{-5}$ | $8 \times 10^{-6}$ | mol/m/s/MPa | [25] |
| $a_1$ | – | Sensitivity of $\kappa$ to $\psi_{xylem}$ | – | $1.1 \times 10^{-5} \pm 0.1 \times 10^{-5}$ | $MPa^{-1}$ | Fit to $g_{\text{oxz}}$ from experiments |
| $\psi_{50}$ | – | Characteristic $\psi_{xylem}$ at which $\kappa$ changes | – | $-0.23 \pm 0.01$ | MPa | Fit to $g_{\text{oxz}}$ from experiments |
| $\kappa_{\text{mem}}^{\text{bs-max}}$ | – | Maximum hydraulic conductivity of bundle sheath plasma membrane | $1.3 \times 10^{-5}$ | $9 \times 10^{-6} \pm 1 \times 10^{-6}$ | mol/m/s/MPa | [25] |
| $\kappa_{\text{mem}}^{\text{bs-min}}$ | – | Minimum hydraulic conductivity of bundle sheath plasma membrane | $2.6 \times 10^{-8}$ | $2 \times 10^{-7} \pm 1 \times 10^{-7}$ | mol/m/s/MPa | [25] |
| | | | Parameters common across hypotheses | | | |
| $k_{\text{vap}}^{\text{bs}}$ | $\frac{cD\bar{v}\chi_0\phi_{\text{bs}}}{RT\tau_{\text{bs}}}$ | Isothermal hydraulic conductivity of vapor in air spaces adjacent to bundle sheath | $\frac{4.05 \times 10^{-7}}{\tau_{\text{bs}}}$ | $\frac{4.05 \times 10^{-7}}{\tau_{\text{bs}}}$ | mol/m/s/MPa | – |
| $k_{\text{vap}}^{\text{mes}}$ | $\frac{cD\bar{v}\chi_0\phi_{\text{mes}}}{RT\tau_{\text{mes}}}$ | Isothermal hydraulic conductivity of vapor in air spaces adjacent to mesophyll | $\frac{4.05 \times 10^{-7}}{\tau_{\text{mes}}}$ | $\frac{4.05 \times 10^{-7}}{\tau_{\text{mes}}}$ | mol/m/s/MPa | – |
| $g_s^{ad}$ | – | Stomatal conductance on adaxial surface | 90 | 90 (or 75) | $mmol/m^2/s$ | Gas exchange data; 75 for Fig. 4h only |

| | | | | | | |
|---|---|---|---|---|---|---|
| $g_s^{ab}$ | – | Stomatal conductance on abaxial surface | 90 | 90 (or 0) | mmol/m$^2$/s | Gas exchange data; 0 for Fig. 4h only |
| $g_{\mathrm{OXZ}}$ | Eq. S27 | OXZ conductance | $[1.5 \times 10^3, 3.8 \times 10^4]$ | $[1.5 \times 10^3, 3.8 \times 10^4]$ | mmol/m$^2$/s | Data |

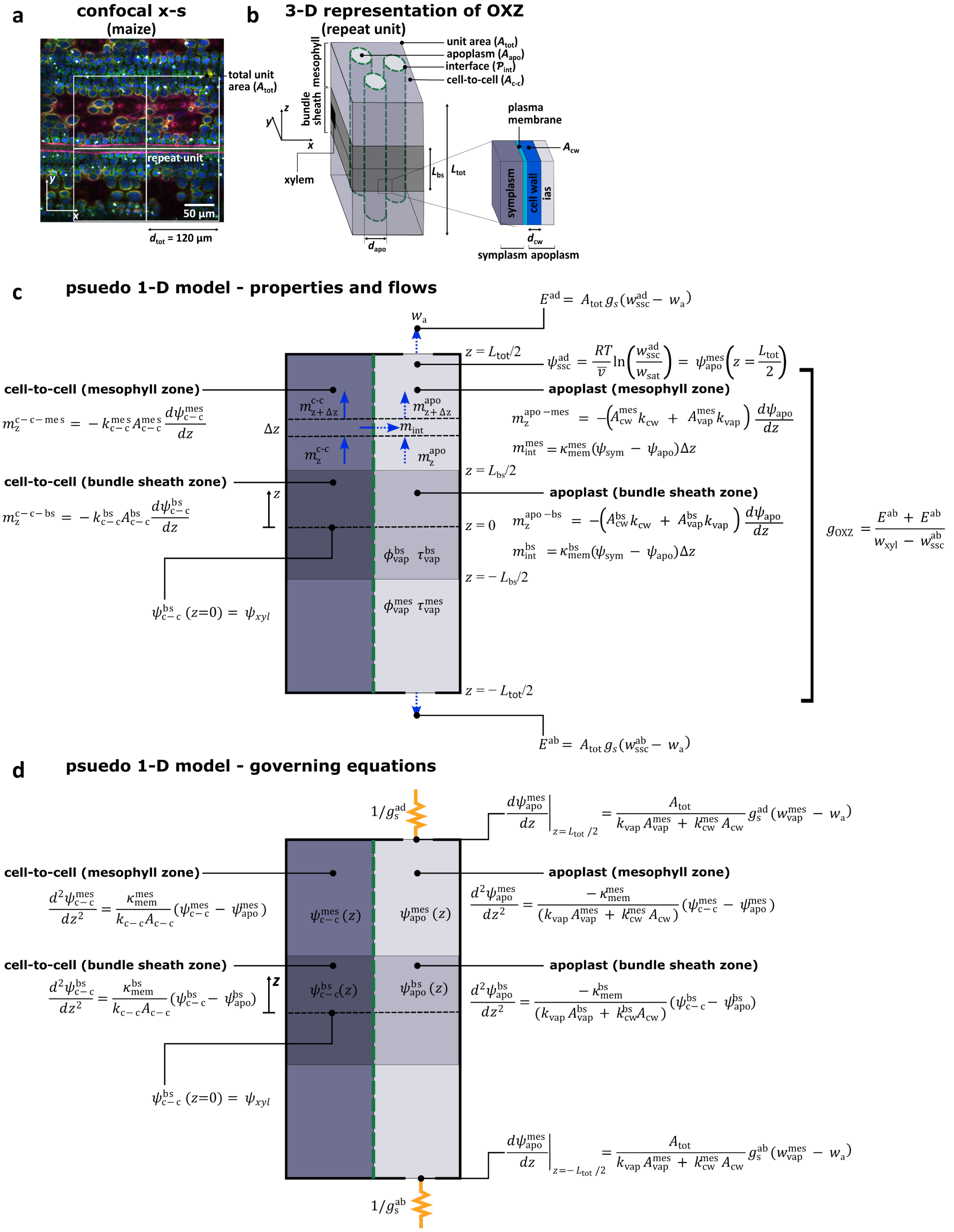

**Figure S19. Pseudo 1-D model of OXZ.**

**a.** Repeat unit defined on a confocal section (in-plane of leaf; $x$-$y$ plane) of a maize leaf showing native autofluorescence. In the vertical direction ($y$-axis), the repeat unit spans the mid-points of adjacent vessels (dense bands of cells in green and blue). In the horizontal direction ($x$-axis), it captures one or two stomates (dark semi-circular regions pass through sub-stomatal cavities). The area of the repeat unit, $A_{tot}$ serves as the reference surface area of leaf for which all mass flows of water through the OXZ are calculated. We take the lateral dimension of the repeat unit, $d_{tot}$ to be $100 - 120$ μm for modelling maize. **b.** 3-D representation of a repeat unit of the OXZ used to define geometry considered in our pseudo 1D model (c and d). Each plane ($x$-$y$ plane) contains cell-to-cell (darker shading) and apoplasmic (lighter shading) paths. To assess local pore geometry in our effective medium treatment, we consider interspersed apoplasmic ($A_{\mathrm{apo}}$ cross-sectional area per repeat unit) and cell-to-cell ($A_{\mathrm{c-c}}$ cross-sectional area per repeat unit) paths of cylindrical cross-section that are principally oriented along the through-leaf axis ($z$-axis); the apoplasmic paths are represented explicitly (diameter, $d_{\mathrm{apo}}$). The interface between these zones (green dashed lines) has perimeter, $\mathcal{P}_{\mathrm{int}}$ [m] per repeat unit and represents the plasma membrane and cell wall (expanded view). The cell wall has cross-sectional area, $A_{\mathrm{cw}}$. Through the thickness (along $z$ – total thickness, $L_{\mathrm{tot}}$), both apoplasmic and cell-to-cell paths are divided into four zones: abaxial and adaxial mesophyll and abaxial and adaxial bundle sheath (thickness, $L_{\mathrm{bs}}$) zones. The geometry and transport parameters of the mesophyll and bundle sheath zones have different values (see Tables S2-S5). For maize, the leaf is treated as having abaxial-adaxial symmetry (same parameters) except with respect to the flux boundary condition defined by stomatal conductance. **c-d.** Representation of pseudo 1-D domains in which transport is modelled with relevant equations for flows in c and for mass balances and boundary conditions in d. Note: while the domains are presented graphically as having lateral extent, mathematically, we account only for gradients and flows along $z$ within each domain and for local exchange between the cell-to-cell and apoplasmic domains at each axial position (flows shown in c for a differential axial zone of length $\Delta z$. See associated text and equations for additional details on model.

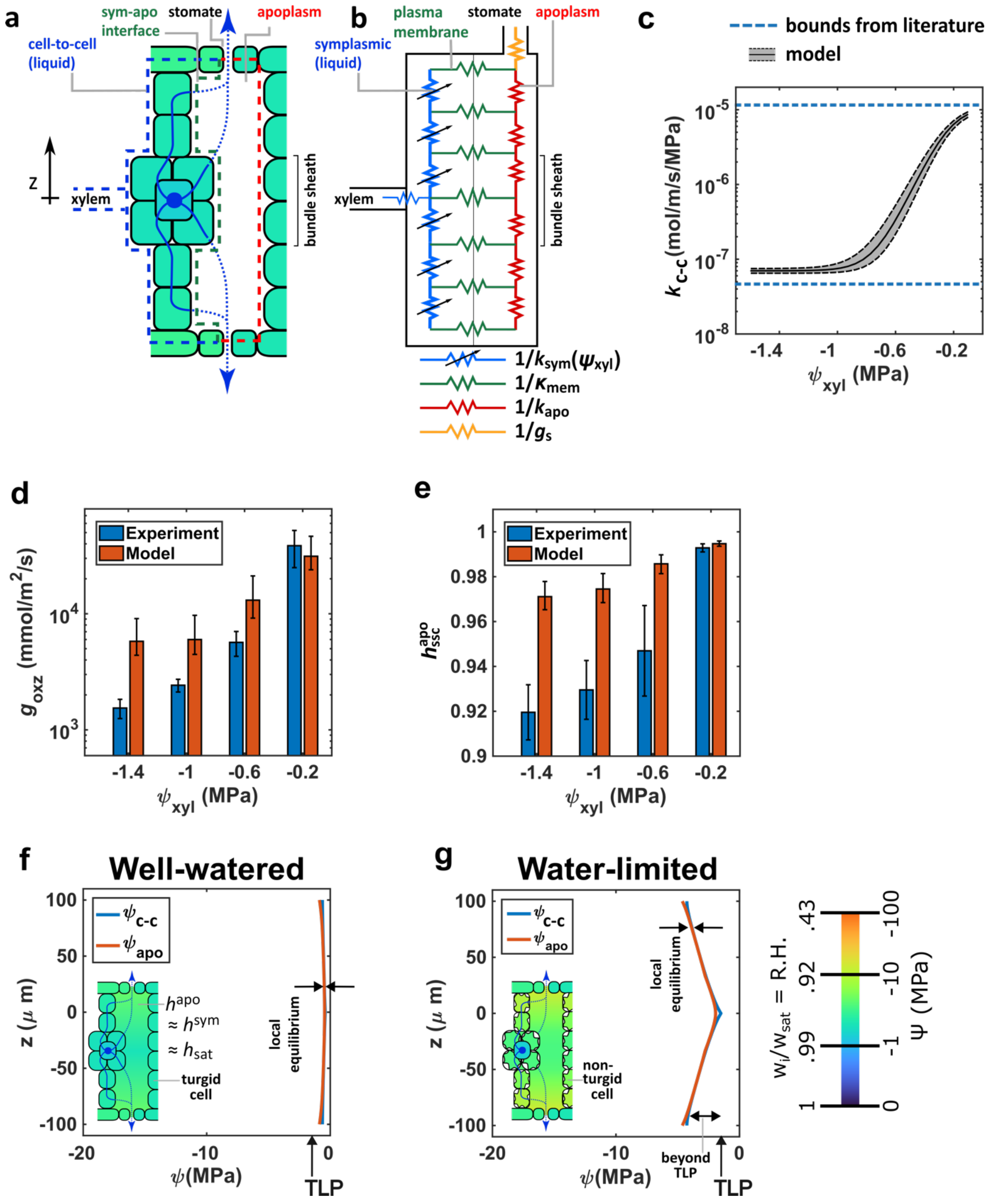

**Figure S20. Alternative hydraulic architecture 1: 'local equilibrium'.**

**a**: Schematic diagram of architecture and flow paths. **b**. Hydraulic architecture in which the symplasm and apoplasm are forced to be in near equilibrium at all times by setting $\kappa_{mem} = 5 \times 10^{-6}$ mmol/m/s/MPa (approximately the highest conductivity measured for a plasma membrane from protoplast-swelling assays). This scenario mimics that captured by the an effective medium with local equilibrium as developed by Rockwell et al.[11] **c**: Functional dependence of conductance of the cell-to-cell path ($k_{c-c}$) on $\psi_{xyl}$ (Eq. S47); this tissue transport parameter function is provided as input to the solver that predicts the distribution of water potential in the symplasm ($\psi_{\mathrm{c-c}}$) and apoplasm ($\psi_{\mathrm{apo}}$) as a function of depth in the leaf z. **d-e**: Comparison of prediction from model of $g_{\mathrm{oxz}}$ (d) and relative humidity, $h_{\mathrm{ssc}}^{\mathrm{apo}}$ (e) based on variable conductance in (c) with experimentally observed values of this tissue conductance (d) and intercellular air space relative humidity (e) as a function of $\psi_{\mathrm{xyl}}$. **f-g**: Profiles of water potential in along the cell-to-cell path ($\psi_{\mathrm{c-c}}(z)$ – blue curve) and apoplasmic path ($\psi_{\mathrm{apo}}(z)$ – orange curve) for well water (f) and water-limited (g) conditions. Schematic representation of local potentials at the interfaces between symplasm and apoplasm. Predicted loss of turgor in mesophyll cells and disagreement in (d) and (e) makes this hydraulic architecture inconsistent with our data.

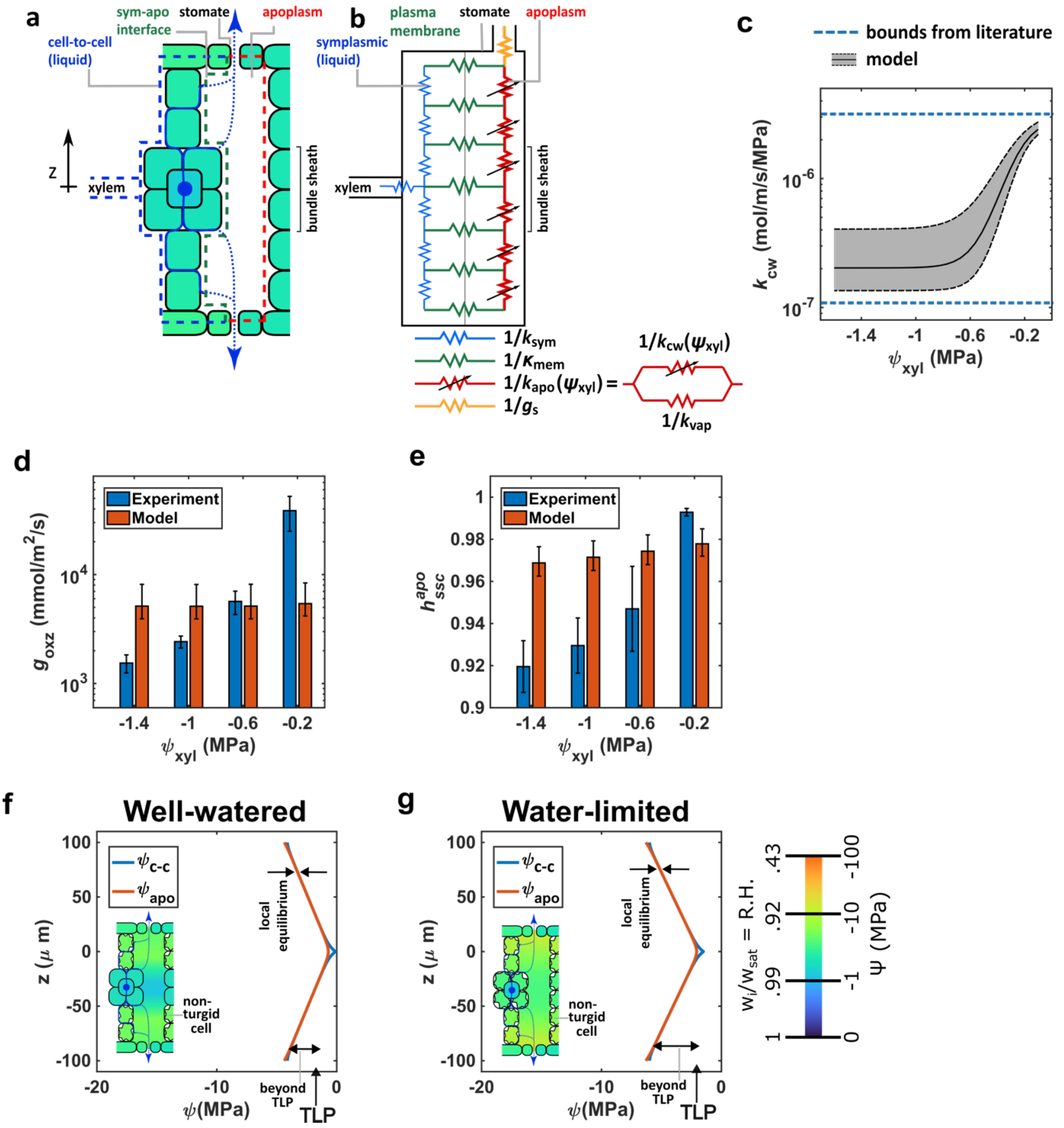

**Figure S21. Alternative hydraulic architecture 2: 'vulnerable cell wall'.**

**a**: Schematic diagram of architecture and flow paths. **b**. Hydraulic architecture in which transport through the cell wall is assumed to be the dominant, variable conductance path for water transport in the leaf as suggested by Buckley et al.[12] Here, flow of water occurs predominantly in the apoplasm through the full thickness of the leaf ($k_{\text{c-c}} < k_{\text{apo}}$). **c**: Functional dependence of conductance of cell wall ($k_{\text{cw}}$) on $\psi_{\text{xyl}}$ (Eq. S48); this tissue transport parameter function is provided as input to the program that predicts the distribution of water potential in the symplasm ($\psi_{\text{c-c}}$) and apoplasm ($\psi_{\text{apo}}$) as a function of depth in the leaf z. **d-e**: Comparison of prediction from model of $g_{\text{oxz}}$ (d) and relative humidity, $h_{\text{ssc}}^{\text{apo}}$ (e) based on variable conductance in (c) with experimentally observed values of this tissue conductance (d) and intercellular air space relative humidity (e) as a function of $\psi_{\text{xyl}}$. **e-g**: Profiles of water potential in along the cell-to-cell path ($\psi_{\text{c-c}}(z)$ – blue curve) and apoplasmic path ($\psi_{\text{apo}}(z)$ – orange curve) for well water (f) and water-limited (g) conditions. Schematic representation of local potentials at the interfaces between symplasm and apoplasm. Predicted loss of turgor in mesophyll cells makes this hydraulic architecture inconsistent with our data. Predicted loss of turgor in mesophyll cells and disagreement in (d) and (e) makes this hydraulic architecture inconsistent with our data.

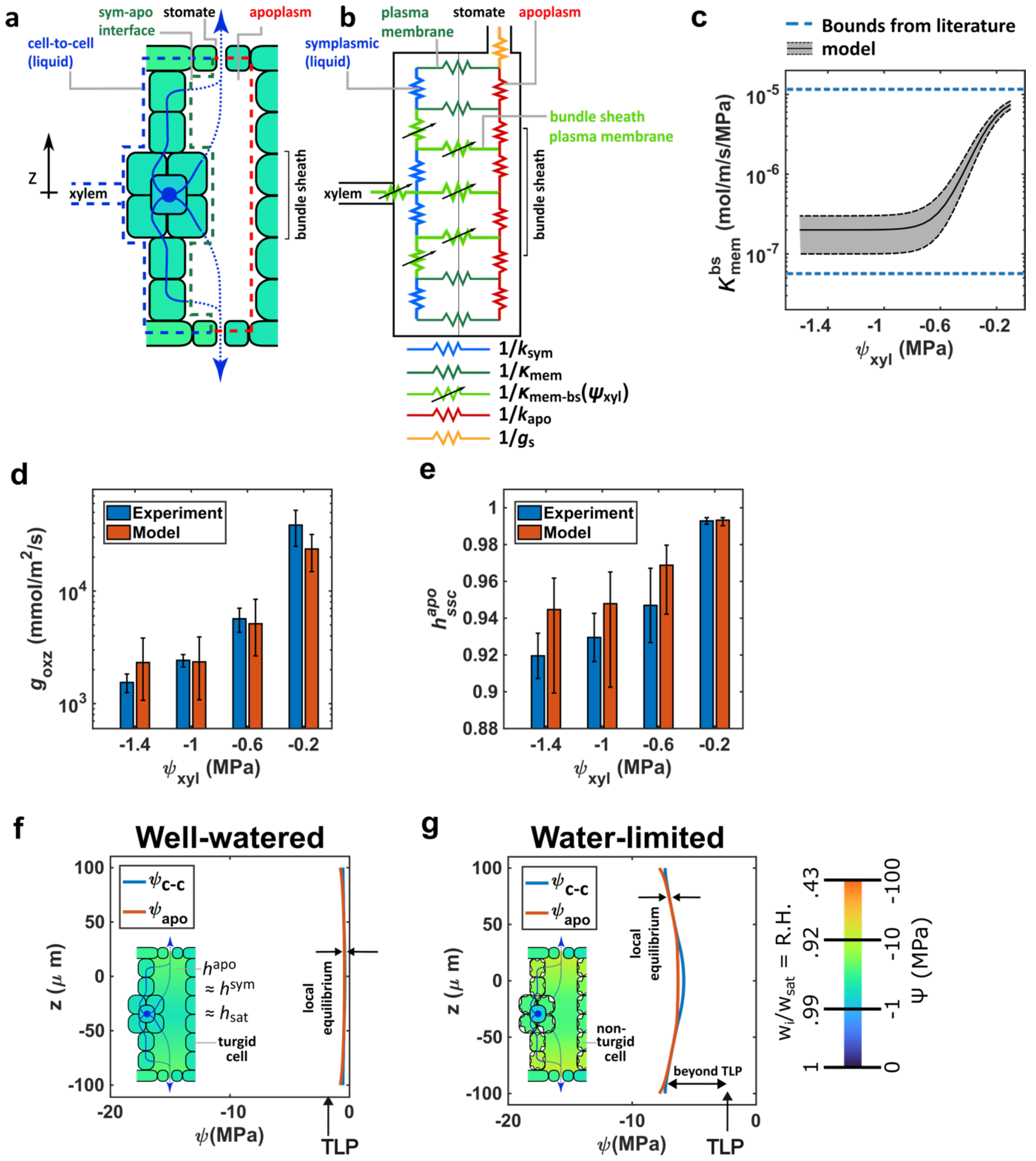

**Figure S22. Alternative hydraulic architecture 3: 'vulnerable bundle sheath'.**

**a**: Schematic diagram of architecture and flow paths. **b**. Variable limiting resistances are attributed to the plasma membrane of the bundle sheath (light green zigzag lines), including in the connection of the OXZ to the xylem, as proposed by Shatil-Cohen et al. 2011.[26] **c**: Functional dependence of conductance of bundle sheath plasma membrane hydraulic conductivity ($\kappa_{\mathrm{mem}}^{\mathrm{bs}}$) on $\psi_{\mathrm{xyl}}$ (Eq. S49); this tissue transport parameter function is provided as input to the program that predicts the distribution of water potential in the symplasm ($\psi_{\mathrm{c-c}}$) and apoplasm ($\psi_{\mathrm{apo}}$) as a function of depth in the leaf z. **d-e**: Comparison of the prediction from model of $g_{\mathrm{oxz}}$ (c) and relative humidity, $h_{\mathrm{ssc}}^{\mathrm{apo}}$ based on variable conductance in (c) with experimentally observed values of this tissue conductance (d) and intercellular air space relative humidity (e) as a function of $\psi_{\mathrm{xyl}}$. **e-g**: Profiles of water potential in along the cell-to-cell path ($\psi_{c-c}(z)$ – blue curve) and apoplasmic path ($\psi_{apo}(z)$ – orange curve) for well water (f) and water-limited (g) conditions. Schematic representation of local potentials at the interfaces between symplasm and apoplasm. Predicted loss of turgor in mesophyll cells makes this hydraulic architecture inconsistent with our data.

**S11. Depth profile of AquaDust measurements from confocal microscopy**

AquaDust measurements from confocal microscopy as a function of depth inside the leaf offer a second check on model predictions of $\psi_{ssc}^{apo}$ described in Section S8. Qualitatively, the water potentials become more negative as one approaches the stomates from inside the leaf, consistent with our expectation of the direction of the water potential gradient. Interestingly, the quantitative assessment points to the existence of a large gradient in water potential (of the order of a few Megapascals) in the vapor phase within the leaf. However, the gradient does not seem to be large enough to bring midplane potentials to a state of saturation, as suggested by Wong et al. 2022.[30] Future studies should aim to resolve gradients of water potential in deeper planes within the leaf.

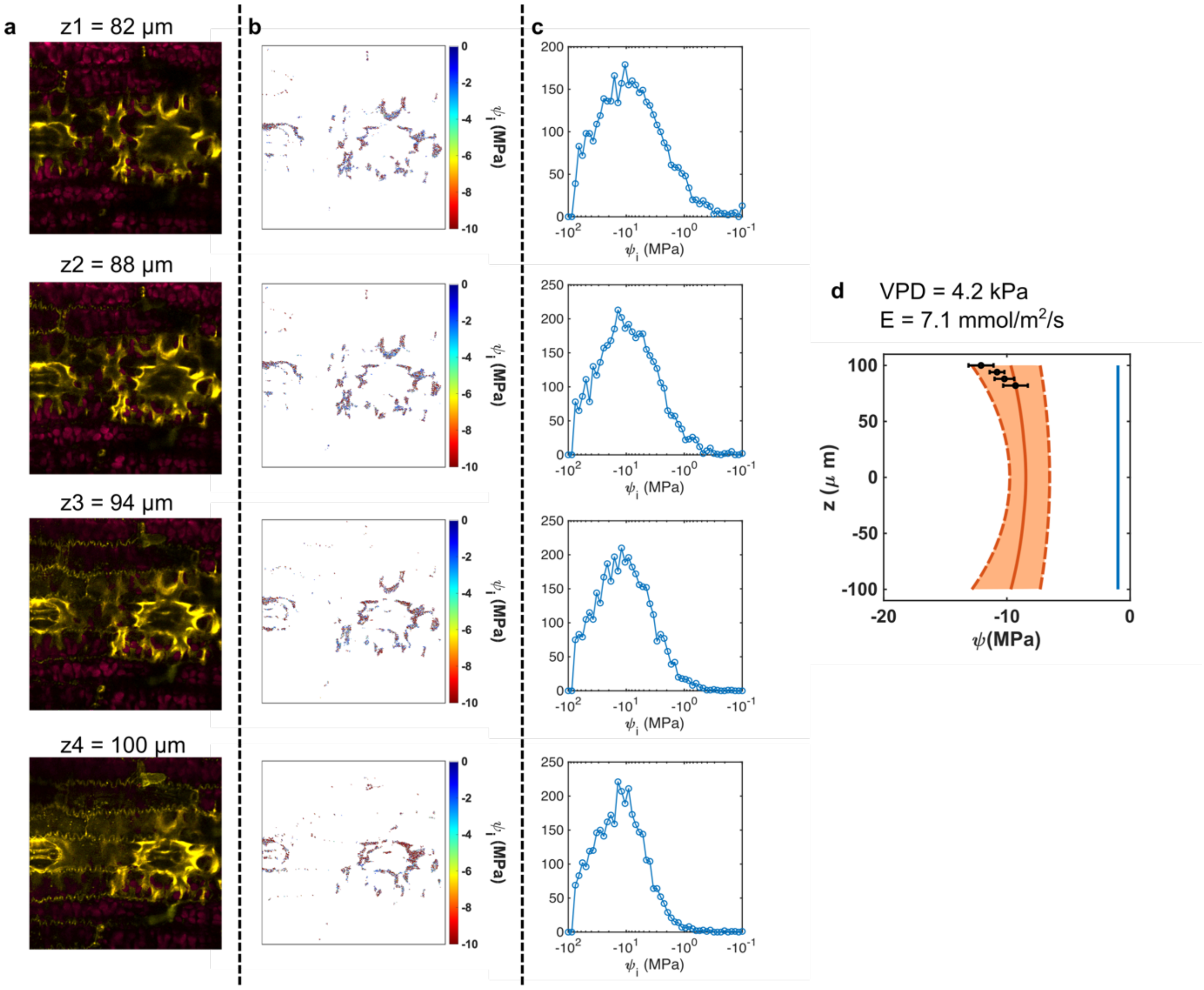

**Figure S23. Cellular scale measurement of $\psi_{scc}^{apo}$ at varying depths.**

**a.** Autofluorescence from chloroplast (false coloured as magenta) overlain with AquaDust signal (false coloured yellow). **b**. Pixel-scale water potential distribution. Thresholding is used to segment AquaDust and non-AquaDust pixels and corresponding water potential is calculated as described by Eq. S1. **c**. Frequency distribution of pixels at different water potential values, peak corresponds to the mode of the distribution. **d**. Prediction from model (orange curve) described in Section S10 overlain on mode from (c). The parameters used here are described in SI Section S10 'Preferred hypothesis'.

**S12. Response of hydraulic conductance of outside-xylem zone $g_{\mathrm{oxz}}$ to Mercury (II) Chloride.**

**NOTE:** Mercuric Chloride ($HgCl_2$) is an acutely toxic chemical that can lead to death upon ingestion or contact with the skin. Review the MSDS and consult with Environmental Health and Safety before use.

Considering the information that $g_{\mathrm{oxz}}$ is solely determined by $\psi_{\mathrm{xyl}}$, we anticipated that its regulation is under biological control and may involve aquaporins. To test this hypothesis, we poisoned aquaporins by feeding aquaporin blocker, $HgCl_2$ through the xylem tissue and measured changes in $g_{\mathrm{oxz}}$ as a function of time.[31–33] All experiments were conducted in the fume hood with proper PPE (see Sec. S13 for full protocol). All plants used were well-watered. One leaf each from 3 biological replicates was cut under water in a wide glass container (closed symbols – Fig. S24). The cut end of the leaf inside the water was scooped into a 20 ml glass vial and appropriate amounts of $HgCl_2$ is added to the 20 ml vial to make up a 1.5 mM $HgCl_2$ solution. The top end of the glass vial was sealed with a parafilm securing the leaf in place. This process was repeated for 1 leaf each on 3 other biological replicates where we did not add any $HgCl_2$ to the glass vial (control – open symbols – Fig. S24). The glass vial was secured in place using a lab stand. The exposed end of the leaf, containing an AQD infiltrated zone was clamped with a gas exchange cuvette, kept at 36 C, VPD 3.5 kPa, to collect simultaneous AquaDust and gas-exchange measurements. The fold change in $g_{\mathrm{oxz}}$ achieved by poisoning aquaporins agrees well with fold-change observed in $g_{\mathrm{oxz}}$ in response to induced drought stress (Fig. 2e – main text).

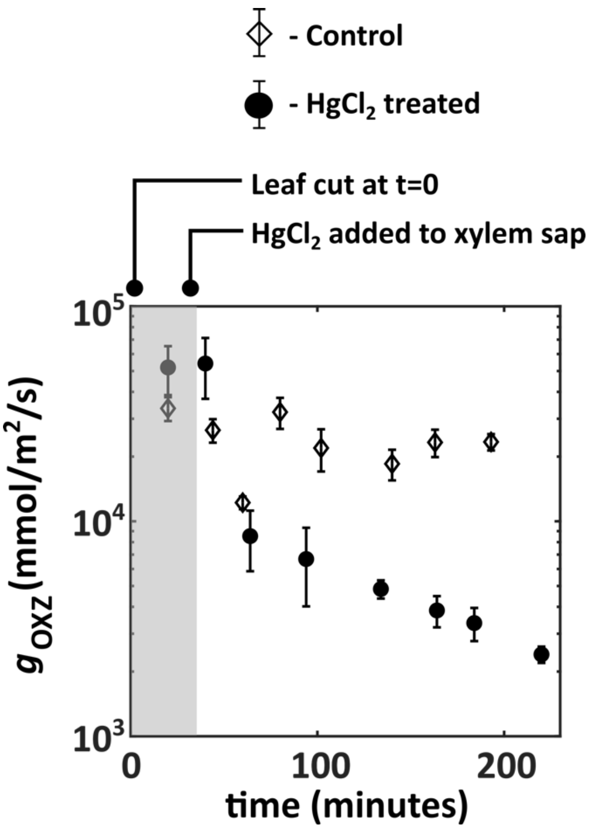

**Figure S24. Response of hydraulic conductance of outside-xylem zone $g_{\mathrm{oxz}}$ to Mercury (II) Chloride.**

**S13. Standard Operating Procedure (SOP) for handling Mercury (II) Chloride ($HgCl_2$) solution for gas-exchange and AquaDust experiments**

**Introduction.**

**NOTE:** Mercuric Chloride ($HgCl_2$) is an acutely toxic chemical that can lead to death upon ingestion or contact with the skin. Review the MSDS and consult with Environmental Health and Safety before using this protocol.

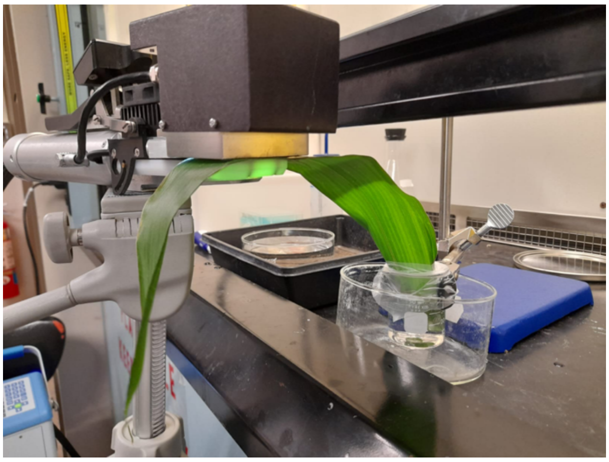

**Figure S25. Photo of experiment in progress with an excised maize leaf with the cut end dipped inside artificial xylem sap solution with 0.5 mM $HgCl_2$.**

**Supplies.**

- $HgCl_2$ stock solution (5% w/v ~ 0.184 M – Ricca Chemical: **MDL Number:** MFCD00011041; **CAS Number:** 7487-94-7) stored in container with an absorbent jacket
- Artificial xylem sap (20 mM KCl, 1 mM CaCl2)
- Mercury spill kit (Uline; Model No. S-19487)
- Lab stand
- Parafilm
- Disposable wooden stirrer to mix $HgCl_2$ with artificial xylem sap
- 40 mL glass beaker for containing xylem sap that will be fed to the excised leaf (a 40 mL beaker is appropriate for an excised maize leaf)
- 2 crystallizing dishes (one for containing artificial xylem sap and one for use as secondary container)
- Mechanical pipette with disposable tips (volume ~ 200 µL)
- Gas exchange system with stable tripod

**Personal Protective Equipment (PPE).**

- Silver shield gloves (Honeywell Safety; VWR Catalog Number: 11000-652)
- Lab coat
- Nitrile gloves

- Face shield
- Goggles
- Impermeable footwear (e.g., snow boots)
- Full-sleeve shirt, full-length pants

**Procedure.**
**NOTE:** All following steps require participation of two persons.
Preparations:

1. Ensure that the use of an acute toxin is indicated on the HASP form of the lab door.
2. Inform other users of the room in which this work will occur about its use.
3. Install sign on hood indicating with GHS labels:

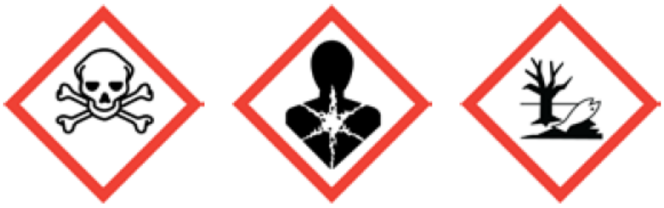

GHS06 GHS08 GHS09

4. Prepare wide-mouth hazardous waste containers (500 mL plastic reagent bottles will suffice) for both solid and liquid waste exposed to $HgCl_2$. These containers MUST be placed inside a secondary container inside the fume hood.
5. Make ~1.5-2 L artificial xylem sap solution in a large crystallizing dish.
6. Cut maize leaf in this crystallizing dish ensuring cut end is submerged in artificial xylem sap solution at all times.
7. Insert the 40 mL beaker into the large crystallizing dish containing artificial xylem sap.
8. Guide the cut end of the maize leaf into the 40 mL beaker ensuring cut end is submerged at all times.
9. Scoop up the artificial xylem sap along with the excised leaf into the 40 mL beaker ensuring cut end is submerged at all times.
10. Adjust volume of solution in 40 mL beaker containing the excised leaf to just above the 40 mL mark.
11. Clamp the 40 mL beaker to a lab stand and secure leaf in position with parafilm, always ensuring that the cut end of the leaf is submerged in sap solution.
12. Place the leaf inside a gas exchange cuvette.
13. The setup at this stage is shown in Figure 1.
14. Perform gas exchange measurements and AqD potential once gas exchange reaches steady state values (~ 25 minutes after placing leaf inside cuvette). At this stage, $HgCl_2$ has not been added to xylem sap. So, this measurement gives us the $K_{ox}$ in the absence of $HgCl_2$.

**NOTE**: all the following manipulations MUST occur inside the fume hood.

15. Before proceeding, the following PPE must be worn:
    a. Face shield
    b. Lab coat

    c. Triple-gloving: inner-most layer: nitrile; second layer: silver shield; third layer: nitrile.

16. Take out the $HgCl_2$ stock solution container from the absorbent jacket.
17. Add $HgCl_2$ (~109 µL of 0.184 M stock solution) to artificial xylem sap in the 40 mL beaker such that final solution has a concentration of 0.5 mM $HgCl_2$.
18. Put the $HgCl_2$ stock solution container back inside the the absorbent jacket.
19. Use wooden stirrer to mix the solution for approximately 30 seconds.
20. Dispose of stirrer in hazardous waste container for solids.
21. Rinse the third/outer-most layer of nitrile gloves thoroughly and dispose of in regular trash can.
22. Dispose of silver shield gloves and use fresh set of gloves for next steps.
23. Lower fume hood sash to safe height.
24. Allow gas exchange to reach steady state values before taking the next set of measurements:
    a. Measure steady state gas exchange (PAR=750: 100% white light, cuvette temperature=36 C, VPD=4 kPa, [CO2-reference]=500 ppm)
    b. Wearing nitrile gloves, remove light module to measure AqD signal through cuvette window. Re-insert light module on to the cuvette for next set of measurements to be taken after ~25-30 minutes.

**Disassembly of setup.**

25. Before proceeding, the following PPE must be worn:
    a. Face shield
    b. Lab coat
    c. Triple-gloving: inner-most layer: nitrile; second layer: silver shield; third layer: nitrile.
26. Remove leaf from cuvette.
27. Raise fume hood sash such that the setup can be handled.
28. Dispose of the parafilm into the hazardous waste container for solids.
29. Unclamp the 40 mL beaker from the lab stand, making sure the solution in the beaker is not spilled.
30. The leaf is taken out of the beaker and disposed of in the hazardous waste container for solids.
31. The solution containing $HgCl_2$ in the 40 mL beaker is disposed of in the hazardous waste container for liquids.
32. Rinse the 40 mL beaker thoroughly (at least 3 times) with water and dispose of the water in the hazardous waste container for liquids.
33. Rinse the third/outer-most layer of nitrile gloves thoroughly and dispose of in regular trash can.
34. Dispose of silver shield gloves and use fresh set of gloves for next steps.
35. Lower fume hood sash to safe height.
36. Leave hazard signage on hood so long a chemical is stored there.

**Completion of experiments involving $HgCl_2$.**

37. Have waste containers collected as hazardous waste with appropriate labels.

38. If additional experiments are NOT planned with $HgCl_2$ within 1 month, have remaining regent collected as hazardous waste for disposal with appropriate labels.
